# Boosting Antitumour Efficacy and Immunity by Boron Neutron Capture Therapy with Size-Controlled Nanoparticles

**DOI:** 10.1101/2025.04.02.646946

**Authors:** Weian Huang, Heon Gyu Kang, Xu Han, Jie Yu, Xiaoxiao Chen, Takushi Takata, Yoshinori Sakurai, Minoru Suzuki, Naoki Komatsu

## Abstract

Boron neutron capture therapy (BNCT) is emerging cancer radiotherapy requiring ^10^B sensitizer. Although boronophenylalanine (BPA)–BNCT is approved clinically in Japan, the low tumour selectivity and retentivity result in the long-time infusion of high doses. Based on our finding of the size-controllable mechanochemical synthesis of boron-10 carbide nanoparticles (^10^B_4_C NPs), the 50 nm size NPs grafted with poly(glycerol), ¹⁰B₄C(50)-PG, is found to show superior tumour selectivity and retentivity to enhance the eradication efficacy at much lower dosage (5 mg [^10^B] / kg (mouse)). The dosage is further reduced by twice neutron irradiation or combination with an immune checkpoint inhibitor (ICI). Antitumour immunity is found to be boosted by ¹⁰B₄C(50)-PG–BNCT to induce abscopal effect to treat remote or metastatic tumours and long-term memory to prevent cancer recurrence. Additionally, minimal side effects and gradual NP excretion are observed for one year. The ^10^B_4_C(50)-PG is concluded to be a promising ^10^B carrier for clinical application of BNCT due to the prominent antitumour efficacy and immune-activation with minimal toxicity.

---

NPs hold significant promise in clinical practice especially as drug carriers to cancer^1^. Currently, various lipid- and protein-based “soft” NPs such as Doxil^®^ and Abraxane^®^ have been approved in clinical cancer therapies^2,3^. However, “hard” inorganic NPs have faced challenges^4,5^, in spite of their unique properties such as high atomic density, robust size and shape, and high extensibility in the surface functionality^6,7^. Typical limitations are low tumour selectivity to result in low therapeutic efficacy^8^ and the potential toxicity to raise safety concerns^9^.

Meanwhile, boron neutron capture therapy (BNCT) is much less harmful than the other radiotherapies^10^, because ^10^B atom generates α-particle (He^2+^) and recoiled lithium nucleus (Li^3+^) very locally within a range of 10 μm upon neutron irradiation^11^. Although BPA and sodium borocaptate (BSH) have been used clinically^11^, they have inherent issues such as low tumour targetability and retentivity^12^. Especially, BPA–BNCT, recently approved in Japan, suffers from the long-time infusion (3 h) of high doses (500 mg / kg)^13^. To overcome these challenges, these boron agents have been conjugated with polymers, encapsulated in micelles or liposomes, and incorporated in inorganic NPs, which still required relatively high dosage (Supplementary Table 1), increasing the potential toxicity *in vivo*^9,14^. Additionally, BNCT’s localized nature limits its efficacy against metastatic cancers^15^. Recent studies suggest combining BNCT with immunomodulators (PD-L1 siRNA^16^ or anti-PD-1 antibodies^17^) induce abscopal effects to distant tumours, though such responses remain elusive with single-modality BNCT via intravenous administration, or even the other radiotherapy alone.

Herein, we firstly establish the size-controllable mechanochemical process to synthesize ^10^B_4_C NPs, which are grafted with PG^18^. Then, we evaluated the size effect of ^10^B_4_C-PG to find that 50 nm core size, or ^10^B_4_C(50)-PG, shows the higher ^10^B delivery efficiency than the other core sizes; ^10^B_4_C(*Y*)-PG (*Y* = 35, 80 and 110), realizing 88% complete regression (CR) by single bolus intravenous injection at much lower dosage (12 mg / kg (mouse)) than BPA in preclinical and clinical practice, as mentioned above. Since millions of boron-10 atoms are densely packed in one ^10^B_4_C(50)-PG particle, its ^10^B dosage is about five times less than that of BPA in clinical use. The dosage is further reduced to half by adding neutron irradiation one more time. Besides, antitumour immunity is boosted by ^10^B_4_C(50)-PG–BNCT to exhibit abscopal effect, in which the distant tumours are suppressed or even eradicated. Therefore, anti-PD-1 antibody is combined with ^10^B_4_C(50)-PG–BNCT at lower dosage to eradicate tumour. Furthermore, long-term immune memory is observed in cured mice which rejects the same and other kinds of murine tumour cells. Moreover, no side effects are observed in the long-term safety studies. Overall, ^10^B_4_C(50)-PG demonstrates a great promise as ^10^B carrier in BNCT for clinical trials.

## Size-controllable mechanochemical synthesis of ^10^B4C NPs and their PG grafting

Although ball milling process has been employed for B_4_C NP syntheses (Supplementary Table 2), the size control has not been realized so far. We controlled the size to be 35, 50, 80 and 110 nm by pretreating both magnesium and ^10^B-enriched bulk boron oxide by ball-milling (Fig. 1a and Extended Data Fig. 1a), giving *p*-Mg and *p*-^10^B_2_O_3_, respectively. They were subjected to the second ball-milling in the presence of graphite under various conditions (Supplementary Table 3). The smaller sizes with 35 and 50 nm, designated as ^10^B_4_C(35) and ^10^B_4_C(50) NPs, were synthesized through pretreatment of bulk ^10^B_2_O_3_ for 20 and 60 min by ball-milling, respectively. The resulting *p*-^10^B_2_O_3_ sizes qualitatively depend on the pretreatment duration (Supplementary Fig. 1) and reflect the ^10^B_4_C NP sizes (Extended Data Fig. 1b). Meanwhile, the unpretreated bulk ^10^B_2_O_3_ gave ^10^B_4_C(80) and ^10^B_4_C(110) NPs with 80 and 110 nm sizes through the second ball milling with new and worn balls, respectively, probably due to the difference in the ball weights and the surface roughness (Supplementary Fig. 2)^19^. The ball milling duration is another parameter to control the size; B_4_C(100) NP was synthesized under the conditions for ^10^B_4_C(80) by changing the duration from 8 h to 6 h with new balls (Supplementary Table 3 and Supplementary Fig. 3).

**Fig. 1.**
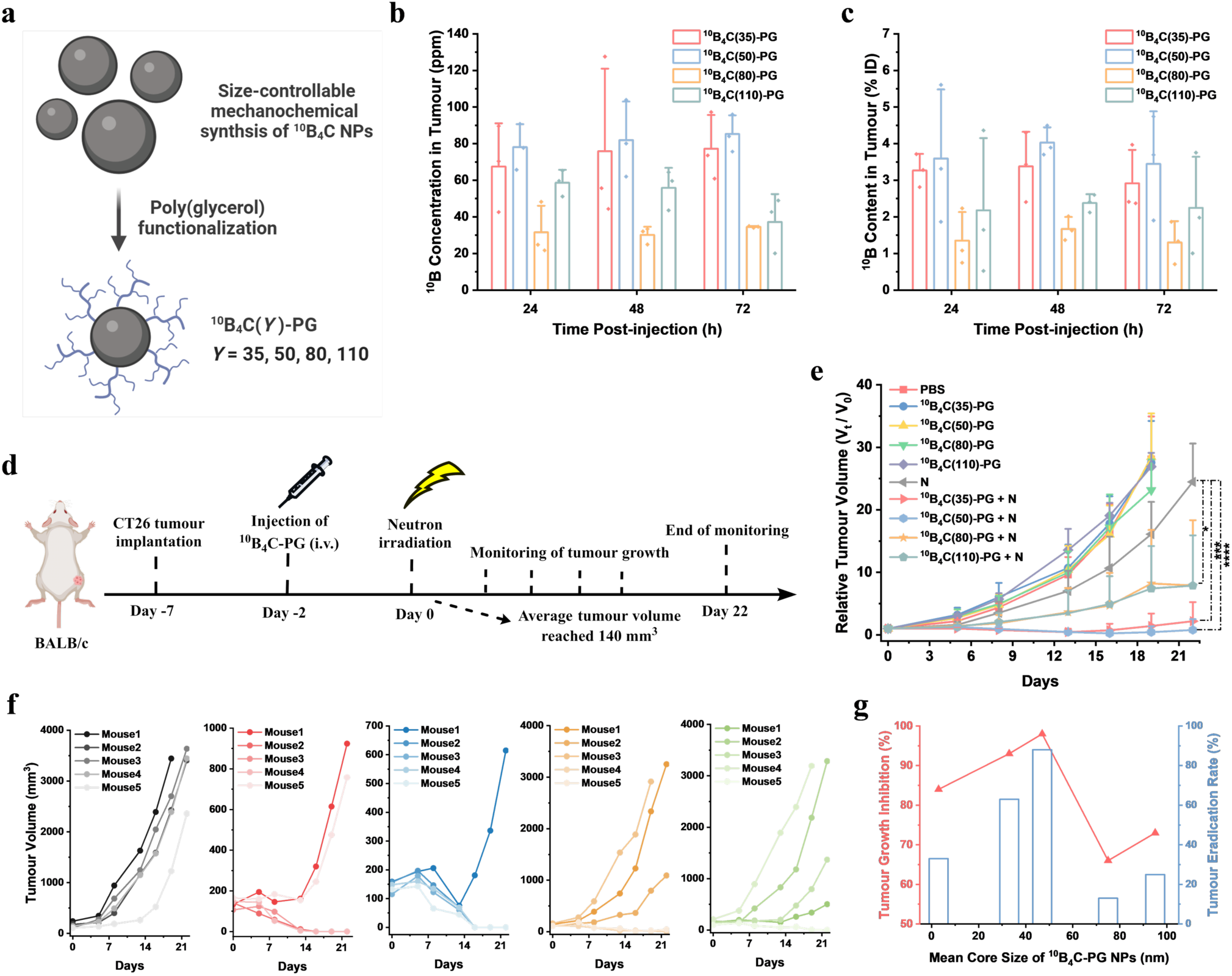
Size effect of ^10^B_4_C-PG on biodistribution and BNCT efficacy. **a,** Size-controlled synthesis of ^10^B_4_C NPs and their PG functionalization. **b,** ^10^B concentration in tumour and blood at 24 - 72 h post-injection of ^10^B_4_C(*Y*)-PG (*Y* = 35, 50, 80 and 110) at a dosage of 5.1 mg [^10^B]/kg (mouse). **c,** %ID of NPs in tumour and blood at 24 - 72 h post-injection. Data are given as the mean ± SD (*n* = 3). **d,** Schematic of CT26 tumour treatment process with dosage of 5.1 mg [^10^B]/kg (mouse). **e,** Relative tumour volume monitored for 22 days. Data are given as the mean ± SD (*n* = 4 in PBS group and *n* = 5 in other groups). **f,** Growth curves of individual tumour mouse in N and ^10^B_4_C(*Y*)-PG + N (*Y* = 35, 50, 80 and 110) groups from left to right. **g,** Summary of size effect of NPs on TGI and tumour eradication rate of ^10^B_4_C-PG–BNCT (*n* = 3 for ^10^B_4_C(5)-PG and *n* = 8 for others). Mean core size of NPs was measured by TEM. Statistical analysis of relative tumour volume with one-way ANOVA post Bonferroni test; \**p* < 0.05, \*\**p* < 0.01, \*\*\**p* < 0.001, \*\*\*\**p* < 0.0001.

The resulting ^10^B_4_C NPs are characterized by X-ray diffraction (XRD), Fourier-transform infrared (FTIR) spectroscopy, scanning electron microscopy (SEM), and dynamic light scattering (DLS). The XRD profiles of the ^10^B_4_C NPs are consistent with that of B_4_C powder reported previously^20^, showing the crystallite sizes of 14.0 – 21.4 nm by Scherrer equation (Extended Data Fig. 1c), which are the same order as that in the DLS sizes (Extended Data Fig. 1b). Therefore, we designate them as B_4_C, though molar ratio of boron / carbon atoms in the NPs is not consistent with the theoretical one, which will be described below. FTIR spectra of ^10^B_4_C NPs are also consistent with the previous report^20^; the bands at 1617 and 1120 cm^−1^ in ^10^B_4_C(50) shift to 1573 and 1090 cm^−1^ in B_4_C(50) including ^10^B and ^11^B at natural abundance, supporting ^10^B enrichment in the samples (Extended Data Fig. 1d and Supplementary Fig. 3)^21^. In addition, the band at 3400 cm^−1^ is attributed to O-H stretching, suggesting high probability to initiate the ring-opening polymerization of glycidol for the subsequent PG functionalization^22^. In the SEM images (Supplementary Fig. 4), ^10^B_4_C NPs have not spherical, but rounded shapes with the same average size order as that determined by DLS.

In phosphate buffer saline (PBS), all the ^10^B_4_C NPs exhibited much larger median diameters and wider distributions than those in water by DLS (Supplementary Fig. 5 and Extended Data Fig. 1b), indicating their aggregation in a physiological environment. For their individualization to enhance the dispersibility and stability in view of *in vivo* applications, ^10^B_4_C NPs were grafted with PG according to the reported method^23^, and the resulting ^10^B_4_C-PG were characterized by FTIR, DLS, thermogravimetric analysis (TGA), transmission electron microscopy (TEM), and zeta potential. In the FTIR (Extended Data Fig. 1d and Supplementary Fig. 6a), the bands at 1114, 2875 and 3340 cm^−1^ are corresponding to C-O-C, C-H and O-H stretching in PG, which are consistent with those reported previously^24^. In the DLS, the median sizes of ^10^B_4_C NPs in water become larger in water and PBS after PG coating (Extended Data Fig. 1e and Supplementary Fig. 6b). This indicates that ^10^B_4_C-PG with all the sizes is individually dispersed in PBS owing to the 35 – 49 wt% of PG coating measured by TGA (Supplementary Fig. 6c). The hydrodynamic sizes of ^10^B_4_C-PG in PBS did not change for a week (Supplementary Fig. 7), showing high colloidal stability of these individualized ^10^B_4_C-PG. The mean core sizes determined by TEM (Supplementary Fig. 8) are almost similar to the median diameters of ^10^B_4_C NPs by DLS (Fig. 1h). The zeta potentials of ^10^B_4_C-PG in water and PBS decreased after PG-functionalization (Extended Data Fig. 1f and Supplementary Fig. 9), due to the conversion of the surface protic functional groups such as boronic acid (–B(OH)_2_) on the ^10^B_4_C NPs to the aprotic boronic esters by initiating the ring-opening polymerization of glycidol^25,26^. Since the ^10^B_4_C NPs are found not to be single crystalline, but polycrystalline, by XRD (Fig. 1c), we determined the actual ^10^B contents to be 33 – 43 wt% in ^10^B_4_C-PG (Extended Data Fig. 1f and Supplementary Fig. 10) by the prompt γ-ray microanalysis (PGRA)^27^. Taking the PG contents into consideration, the ^10^B contents in the cores are calculated to be 65 – 67 wt%, which are smaller than the theoretical value; 77 wt% in ^10^B_4_C. This is probably due to the oxygen-containing functional groups such as hydroxy ones on the NP surface, which is confirmed by FTIR mentioned above. In the Raman spectra (Supplementary Fig. 11)^28^, the amorphous carbon is detected, reducing the ^10^B content. However, the boron-10 carbide NPs synthesized mechanochemically in this work are designated as ^10^B_4_C based on their XRD profiles (Extended Data Fig. 1c).

## Size effect of ^10^B4C-PG on biodistribution and BNCT efficacy

Since the NP size is known to affect the tumour targetability^1^, we tried to find the optimum size of ^10^B_4_C-PG *in vivo* based on the size-selective syntheses of ^10^B_4_C NPs mentioned above. After no obvious cytotoxicity of ^10^B_4_C-PG with all the sizes was confirmed *in vitro* by murine colon cancer CT26 cells (Supplementary Fig. 12), biodistribution was investigated by CT26 bearing mice at 24, 48 and 72 h after intravenous injection. All the core sizes from 35 to 110 nm of the ^10^B_4_C-PG showed tumour ^10^B concentration ([^10^B]) above the BNCT criteria (20 ppm, Fig. 1b) at all the time points (24, 48 and 72 h) and tumour %ID (injection dose) higher than 0.7%ID (median) based on the literature survey (Fig. 1c)^8^. The ^10^B_4_C-PG with each size keeps a similar range of %ID and [^10^B] in tumour at all the time points, and all the %ID increases from 24 h to 48 h (Fig. 1b, c). These phenomena clearly indicate higher tumour retentivity and longer blood circulation of ^10^B_4_C-PG (Supplementary Fig. 13a – 13d). However, %ID in tumour decreased in all the sizes from 48 to 72 h, implying that ^10^B_4_C-PG are retentive in tumour, but excreted gradually (Fig. 1c)^31^. The [^10^B] ratio in tumour and blood (T/B ratio) increased significantly in all sizes from 3 – 7 at 24 h to 17 – 21 at 72 h (Supplementary Fig. 14), which are matched with the BNCT criteria (> 3) to secure less side effect. On the other hand, clear size effect is observed in tumour accumulation. More than 4.0 and 3.4%ID, or 80 and 75 ppm [^10^B], were detected at 48 h by ^10^B_4_C(50)-PG and ^10^B_4_C(35)-PG, respectively, while ^10^B_4_C(80)-PG and ^10^B_4_C(110)-PG accumulated less amounts; 1.6 – 2.4%ID or 30 – 56 ppm [^10^B] at 48 h (Fig. 1b, c). The above %ID of ^10^B_4_C(50)-PG and ^10^B_4_C(35)-PG are the highest among all the boron-10 drugs for cancered mice in BNCT reported so far, including recently reported poly(vinyl alcohol)-BPA and fructose-BPA complexes (1.4 and 0.7%ID, respectively)^32^ as summarized in Supplementary Table 1. In the organs, all the sizes showed low %ID in kidney regardless of time due to their sizes larger than the pore sizes of the glomerular filtration membrane^33^, meaning little excretion through urine. The two smaller NPs are much stealthier than the larger ones based on the less amounts accumulated in liver and spleen (Supplementary Fig. 13). These phenomena should result in more tumour accumulation through longer blood circulation. Based on the highest %ID in tumour for all the sizes (Fig. 1d), we determined the neutron irradiation at 48 h post-administration in the following *in vivo* BNCT experiments.

According to the procedure (Fig. 1d), the tumour volume of the CT26 implanted mice were monitored after intravenous injection of PBS or ^10^B_4_C-PG with four sizes with and without neutron irradiation (N); that is, PBS, ^10^B_4_C(*Y*)-PG, N, and ^10^B_4_C(*Y*)-PG + N (*Y* = 35, 50, 80, and 110). The tumour growth was more efficiently suppressed in the ^10^B_4_C(*Y*)-PG + N groups as compared with the other groups (Fig. 1e). Especially, tumours were eradicated at 60% and 80% of the mice in the ^10^B_4_C(35)-PG + N and ^10^B_4_C(50)-PG + N groups, respectively (Fig. 1f). These results were reproduced at 67% and 100% of the tumour eradication in ^10^B_4_C(35)-PG + N and ^10^B_4_C(50)-PG + N groups, respectively (Supplementary Fig. 15). The hematoxylin and eosin (H&E) staining shows that the tissue slices at the tumour region in ^10^B_4_C(35)-PG + N and ^10^B_4_C(50)-PG + N groups are different from those in the other groups (Supplementary Fig. 16), supporting tumour eradication in the two groups. In addition, pronounced pyknosis, fragmentation and dissolution of cells are observed, reflecting the necroses^34^. Besides, the tissues of liver, spleen and kidney in all the groups were verified to be healthy by H&E staining after 22 days (Supplementary Fig. 17). Although eradiation rates were much lower with larger ^10^B_4_C-PG (Supplementary Fig. 17), about four times higher dosage enhanced the eradiation rate to 100% (Supplementary Fig. 18). In addition, ^10^B_4_C(5)-PG with 5 nm core size was separated from ^10^B_4_C(35)-PG dispersion by centrifugation, characterized by TGA, DLS and TEM (Supplementary Fig. 19), and used in BNCT. The lower therapeutic efficacy and eradiation rate were shown as compared to those of ^10^B_4_C(35)-PG and ^10^B_4_C(50)-PG (Supplementary Fig. 20). No intrinsic toxicity was observed in all the *in vivo* experiments including NP injection and/or neutron irradiation, as evidenced by no mouse fatalities and no significant changes in body weight (Supplementary Fig. 21). Overall, ^10^B_4_C(50)-PG show the highest ^10^B delivering efficiency to tumour (Fig. 1c), tumour eradication rate (Fig. 1g), and tumour growth inhibition (TGI, %) (see Methods) upon neutron irradiation at the dosage of 5.1 mg [^10^B]/kg (mouse) (Fig 1c and g).

The boron-10 containing drugs such as BPA have relied on high dosages to compensate for poor delivery efficiency to and poor retentivity in tumour in BNCT (Extended Data Fig. 2a). Our analysis based on the previous reports shows that only 32% and 19% of the boron drugs achieve high delivery efficiency to tumour with low dosage (HDLD) and high TGI with low dosage (HTLD) as shown in Extended Data Fig. 2a and 2b, respectively. While only 17% of the drugs achieve tumour eradication rate above 50%, ^10^B_4_C(50)-PG is the only drug to eradicate tumour consistently across models (Extended Data Fig. 2c). Among them, ^10^B_4_C(35)-PG and ^10^B_4_C(50)-PG can achieve HDLD, HTLD and tumour eradication simultaneously due to optimal size and the protein shielding property of PG grafting to result in the better stealth efficiency to slip through mononuclear phagocyte system (MPS)^18^. ^10^B_4_C(50)-PG is better than ^10^B_4_C(35)-PG probably due to better tumour penetration and retentivity with 50 nm size in addition to higher [^10^B] in tumour^31,35^. Furthermore, review of the studies on size effect of NPs shows that 30 – 60 nm NPs achieve superior tumour delivery efficiency (Supplementary Table 4), aligning with our findings.

## ^10^B_4_C(50)-PG–BNCT for various tumour models and conditions

For scope and limitation of ^10^B_4_C(50)-PG–BNCT, antitumour efficacy was evaluated by the mouse models with larger initial tumour volumes of CT26 and different murine tumour cell types of breast cancer (4T1), sarcoma (Meth-A), melanoma (B16-F10) and Lewis lung carcinoma (LLC) according to the processes shown in Fig. 2a. The tumour growth was efficiently suppressed in the mice with about 1.2 times larger initial tumour volumes by ^10^B_4_C(50)-PG + N (Fig. 2b, Supplementary Fig. 22a), though the eradication rate slightly decreased from 88% (Fig. 1g) to 50% (Supplementary Fig. 22b). In the mice with more than four times larger initial volumes, the tumour growth suppression was not so efficient in the ^10^B_4_C(50)-PG + N group at the same NP and neutron dosages (Fig. 2d), and accordingly, no tumour was eradicated (Supplementary Fig. 23). While antitumour efficacy depends on the initial tumour volume, the ^10^B and neutron dosages as well as irradiation times can adjust for tumour initial volume to enhance the efficacy.

**Fig. 2.**
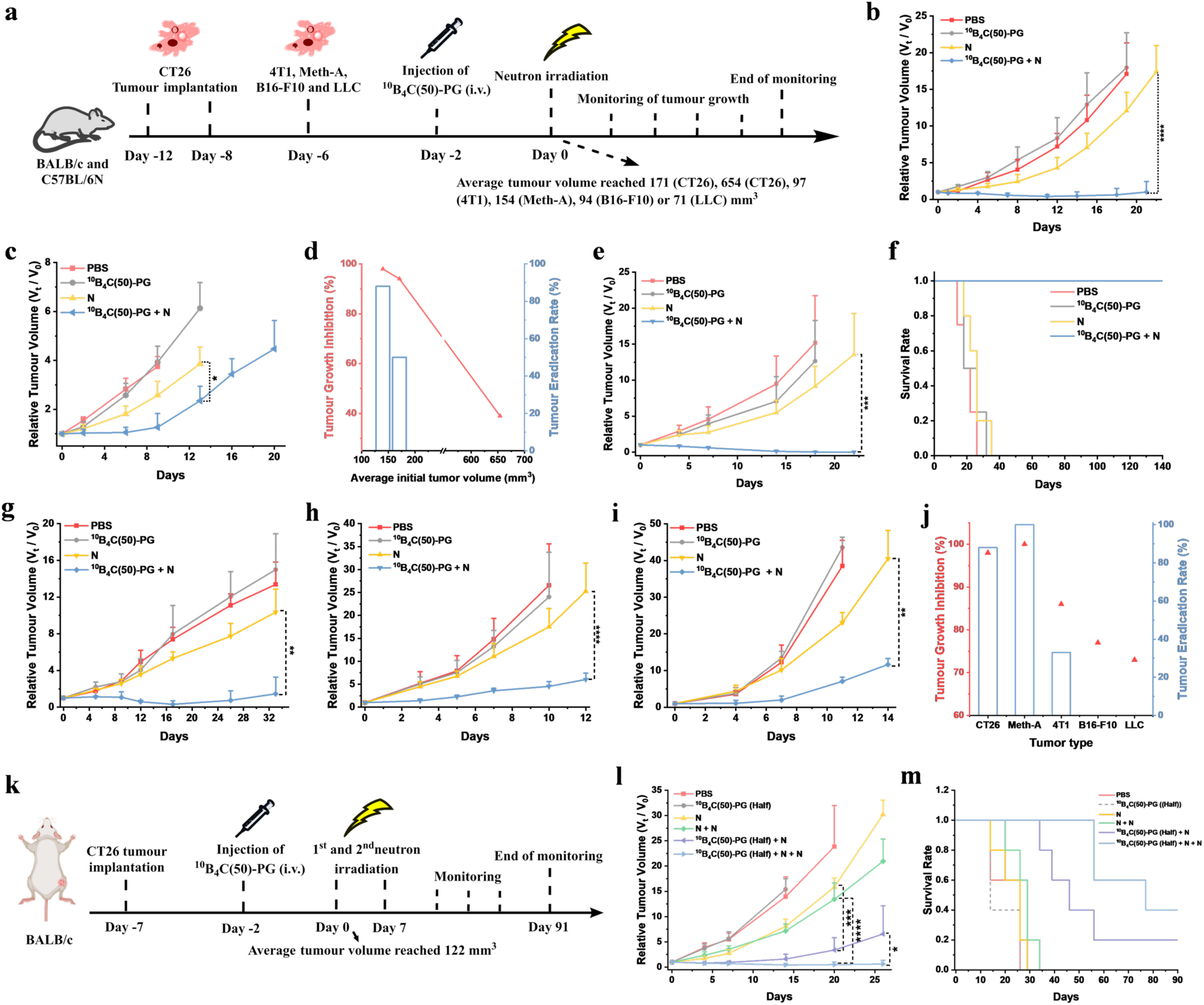
BNCT antitumour efficacy of ^10^B_4_C(50)-PG depending on tumour initial volumes, tumour cell types and neutron irradiation times. **a**, Treatment processes of CT26, Meth-A, B16-F10 and LLC tumours at dosage of 5.1 mg [^10^B]/kg (mouse) and 4T1 tumour at dosage of 5.2 mg [^10^B]/kg (mouse). **b**, Relative tumour volume monitored for 21 and 22 days (*n* = 12 in ^10^B_4_C(50)-PG + N group and *n* = 6 in other groups with average initial volume of 171 mm^3^). **c**, Relative CT26 tumour volume monitored for 20 days (*n* = 4 with average initial volume of 645 mm^3^). **d**, Initial tumour volume dependence of ^10^B_4_C(50)-PG on therapeutic efficacy and eradication rate. **e**, Relative Meth-A tumour volume monitored for 22 days (*n* = 5). **f**, Survival rate of mice in four groups in Meth-A tumour model. **g**, Relative 4T1 tumour volume monitored for 33 days (*n* = 3). **h**, Relative B16-F10 tumour volume monitored for 12 days (*n* = 15 in ^10^B_4_C(50)-PG + N group and *n* = 6 in other groups). **i**, Relative LLC tumour volume monitored for 14 days (*n* = 3). **j**, Summary of tumour cell type dependence on therapeutic efficacy and eradication rate. **k**, Treatment process of CT26 tumour model by twice neutron irradiation at dosage of 2.5 mg [^10^B]/kg (mouse). **l,** Relative CT26 tumour volume monitored for 26 days (*n* = 5). **m**, Survival rate of mice in six groups in **l**. Statistical analysis of relative tumour volume with one-way ANOVA post Bonferroni test; \**p* < 0.05, \*\**p* < 0.01, \*\*\**p* < 0.001, \*\*\*\**p* < 0.0001.

For the other tumour models than CT26, biodistribution of ^10^B_4_C(50)-PG was investigated by 4T1, B16-F10 and LLC mouse models. As shown in Supplementary Fig. 24, their trends are similar to that of CT26 (Fig. 1b – 1c and Supplementary Fig. 13). Although the [^10^B] in these tumours are lower than that of CT26 at 48 h post-injection, all of them are much higher than 20 ppm. In addition, T/B ratios are much higher than 3 (Supplementary Table 1). The antitumour efficacy was evaluated by *in vivo* BNCT with four kinds of tumour models including Meth-A in addition to the above three models (Fig. 2a). Although the tumour growth was suppressed in ^10^B_4_C(50)-PG + N groups in all the models, the efficacy was different significantly (Fig. 2e – 2i). All the Meth-A mice in ^10^B_4_C(50)-PG + N group reached complete remission and survived over 140 days (Fig. 2e, 2f and Supplementary Fig. 25). Meanwhile, the eradication rates were 33% and less in the other models (Fig. 2g - 2j, and Supplementary Fig. 26 and 27), indicating that the therapeutic parameters should be optimized for different tumour models. In addition, no intrinsic toxicity was observed following NP injection and neutron irradiation, as evidenced by the absence of mouse fatalities and significant changes in body weights (Supplementary Fig. 28).

To utilize the high retentivity of ^10^B_4_C(50)-PG in tumour, neutron was irradiated twice with the interval of one week and one day as shown in Fig. 2k and Supplementary Fig. 30a, respectively. The double neutron irradiation at the half dosage (2.5 mg [^10^B]/kg (mouse)), ^10^B_4_C(50)-PG (Half) + N + N, exhibited better tumour growth suppression and higher survival rate than the other conditions (Fig. 2l, 2m and Supplementary Fig. 29) with gradual increase of the weights (Supplementary Fig. 31). On the other hand, higher eradication rate 83% was observed at the double irradiation with one day interval, though the mice suffered from significant decrease in their body weights (Supplementary Fig. 30).

## ^10^B_4_C(50)-PG–BNCT for antitumour immunity

Since radiotherapy is not systemic, but local cancer treatment, it is not effective for metastatic cancer. Recently, a distant tumour has been responded by radio-irradiation directed at a primary one, designated as abscopal or bystander effect, in combination with immunotherapy^15^; for example, BNCT with ^10^B-containing polymer based NPs loaded with PD-L1 siRNA^16^ and with BPA combined with anti-PD-1 antibody^17^. However, abscopal effect has been rarely observed by single modality of BNCT with intravenous injection of ^10^B drug and even the other radiotherapies alone^36^.

To treat distant and primary tumours simultaneously by ^10^B_4_C(50)-PG–BNCT, the CT26 tumours were established at both right and left thighs of the mice, and neutron was irradiated only at the right thigh after intravenous injection of ^10^B_4_C(50)-PG (Supplementary Fig. 36a). While CT26 tumour in the right thigh was eradicated at 29% in the ^10^B_4_C(50)-PG + N group, the tumour growth in the left thigh was significantly suppressed despite no direct neutron irradiation (Supplementary Fig. 32b and c). As a result, the mice of the ^10^B_4_C(50)-PG + N group showed milder weight increase and longer median survival time (Supplementary Fig. 32d and e). The CT26 tumours with smaller volumes significantly increased the eradication rates to 100% and 60% in the right and left thighs, respectively, and survival ratio to 60% over 100 days (Supplementary Fig. 33). Then, we investigated the tumour immune microenvironment (TIM) to confirm immune stimulation by ^10^B_4_C(50)-PG–BNCT. The immunohistochemical staining was performed to primary and distant tumours after 22 days post-irradiation to count the numbers of CD3^+^, CD4^+^ and CD8^+^ T cells with aid of machine learning (Fig. 3a, Supplementary Fig. 34 and 35). The results show that the CD3^+^, CD4^+^ and CD8^+^ T cells in the ^10^B_4_C(50)-PG + N group are much more than those in the other groups (Fig. 3b and Supplementary Fig. 35b and d).

**Fig. 3.**
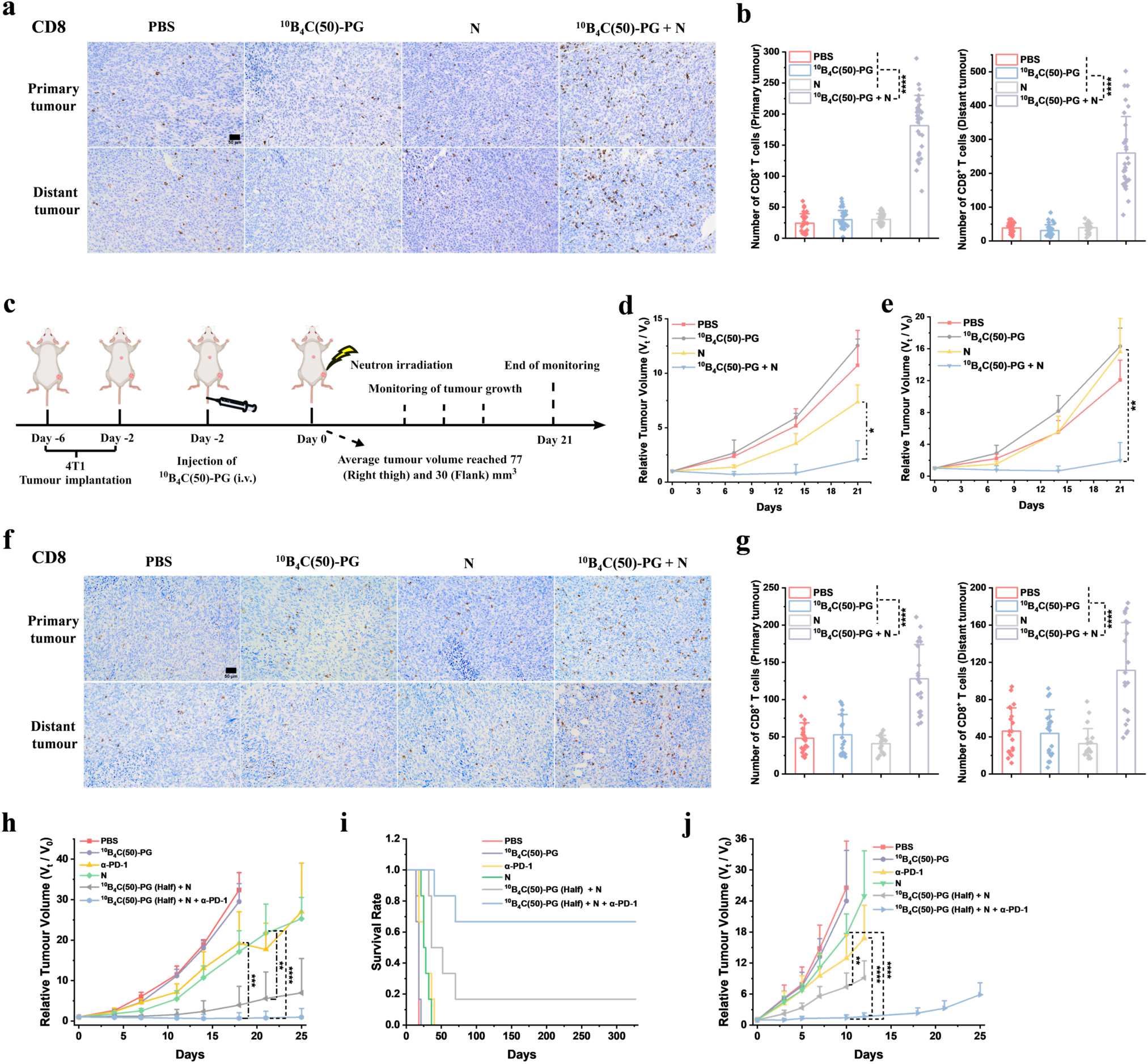
^10^B_4_C(50)-PG–BNCT enhances antitumour immunity, induces abscopal effect, and synergizes well with immunotherapy. **a,** Representative images of tumour sections examined by immunohistochemistry at day 22 in dual CT26 tumour treatment in Supplementary Fig. 33. **b**, Quantification of CD8^+^ T cell numbers in primary and distant CT26 tumours in different groups from immunohistochemistry images after recognizing T cells using machine learning tool FIJI (*n* = 30). **c**, Schematic of dual 4T1 tumour treatment process with dosage of 5.1 mg [^10^B]/kg (mouse) and neutron irradiation only at right thigh in *in vivo* BNCT (*n* = 3). **d,** Relative primary tumour volume at right thigh monitored for 21 days. **e,** Relative distant tumour volume at flank monitored for 21 days. **f,** Representative images of the tumour sections examined by immunohistochemistry at day 21 in dual 4T1 tumour treatment. **g,** Quantification of CD8^+^ T cell numbers in primary and distant 4T1 tumours in different groups from immunohistochemistry images after recognizing T cells using machine learning tool FIJI (*n* = 20). **h,** Relative CT26 tumour volume monitored for 25 days after BNCT at dosage of 2.5 mg [^10^B]/kg (mouse) with αPD-1 (*n* = 6). **i,** Survival rate of mice in different groups in CT26 tumour model *in vivo* BNCT with αPD-1 (*n* = 6). **j,** Relative B16-F10 tumour volume monitored for 25 days after BNCT at dosage of 2.5 mg [^10^B]/kg (mouse) with αPD-1 (*n* = 5 for αPD-1 group and n = 6 for others). Scale bar in immunostaining, 50 µm. Data are given as the mean ± SD. Statistical analysis of relative tumour volume with one-way ANOVA post Bonferroni test; \**p* < 0.05, \*\**p* < 0.01, \*\*\**p* < 0.001, \*\*\*\**p* < 0.0001.

Although these results indicates that the TIM becomes “hot” from “cold” after ^10^B_4_C(50)-PG– BNCT, the tumour growth suppression in left thighs (Supplementary Fig. 32c) can not be concluded solely as a result of the abscopal effect due to the poor linearity of neutron (Supplementary Fig. 36). Since much less thermal neutron fluence was counted in the flank (Supplementary Fig. 36c), 4T1 tumours were established at the flank together with the right thigh of the mice. Neutron was irradiated only at the right thigh after intravenous injection of ^10^B_4_C(50)-PG (Fig. 3c). The tumour growth was significantly suppressed in both flank and right thigh at the eradication rates of 33% and less metastatic areas were observed in lung (Fig. 3d, e and Supplementary Fig. 37), implying the abscopal effect. The immunohistochemical staining of primary and distant tumours (Fig. 3f) indicates that populations of CD4^+^ and CD8^+^ T cells in the ^10^B_4_C(50)-PG + N group are much more than those in the other groups (Fig. 3g). These results imply that the abscopal effect can be caused by the TIM boosted by ^10^B_4_C(50)-PG–BNCT.

Since the ICIs are known to be more effective in the “hot” rather than “cold” TIM^37^, anti-PD-1 antibody (αPD-1) was administrated intraperitoneally to various tumour mouse models after ^10^B_4_C(50)-PG–BNCT to confirm synergy in these modalities (Supplementary Fig. 39). Even at the half dosage in BNCT, 67% of the CT26 mice responded completely and survived over 300 days at the ^10^B_4_C(50)-PG (Half) + N + αPD-1 group (Fig. 3h, 3i and Supplementary Fig. 40). Although the other tumour models bearing B16-F10, 4T1 and LLC were not eradicated, the synergistic therapy showed higher efficacy than BNCT or ICI alone (Fig. 3j and Supplementary Fig. 41 - 43). In all the experiments, no intrinsic toxicity was observed as evidenced by the body weight (Supplementary Fig. 41 – 44).

## ^10^B_4_C(50)-PG–BNCT for immune memory

Preventing tumour recurrence is an integral focus of cancer therapy. Despite significant advancements in treatment, recurrence remains major concern, particularly for certain cancer types^38^. One of the reasons for recurrence is the lack of well-trained immunity^39^. Since the antitumour immunity was activated by ^10^B_4_C(50)-PG–BNCT as mentioned above, we investigated long-lasting antitumour effect or immune memory. CT26 and 4T1 tumour cells were implanted at the left thigh in the mice whose CT26 tumour in the right thigh was eradicated by BNCT (Fig. 2b and Supplementary Fig. 30), designated as Cured (CT26) and Cured (4T1), respectively, as illustrated in Fig. 4a. In parallel, the same tumour cells were implanted at the left thigh in the age-matched untreated mice, designated as Naive (CT26) and Naive (4T1), respectively. While rapid tumour growth was observed in Naive (CT26) and Naive (4T1), CT26 and 4T1 cells did not engraft at all in Cured (CT26) and mostly in Cured (4T1), respectively (Fig. 4b and Supplementary Fig. 45), exhibiting 100% and 67% survival rates after 120 days post tumour cell implantation (Fig. 4c).

**Fig. 4.**
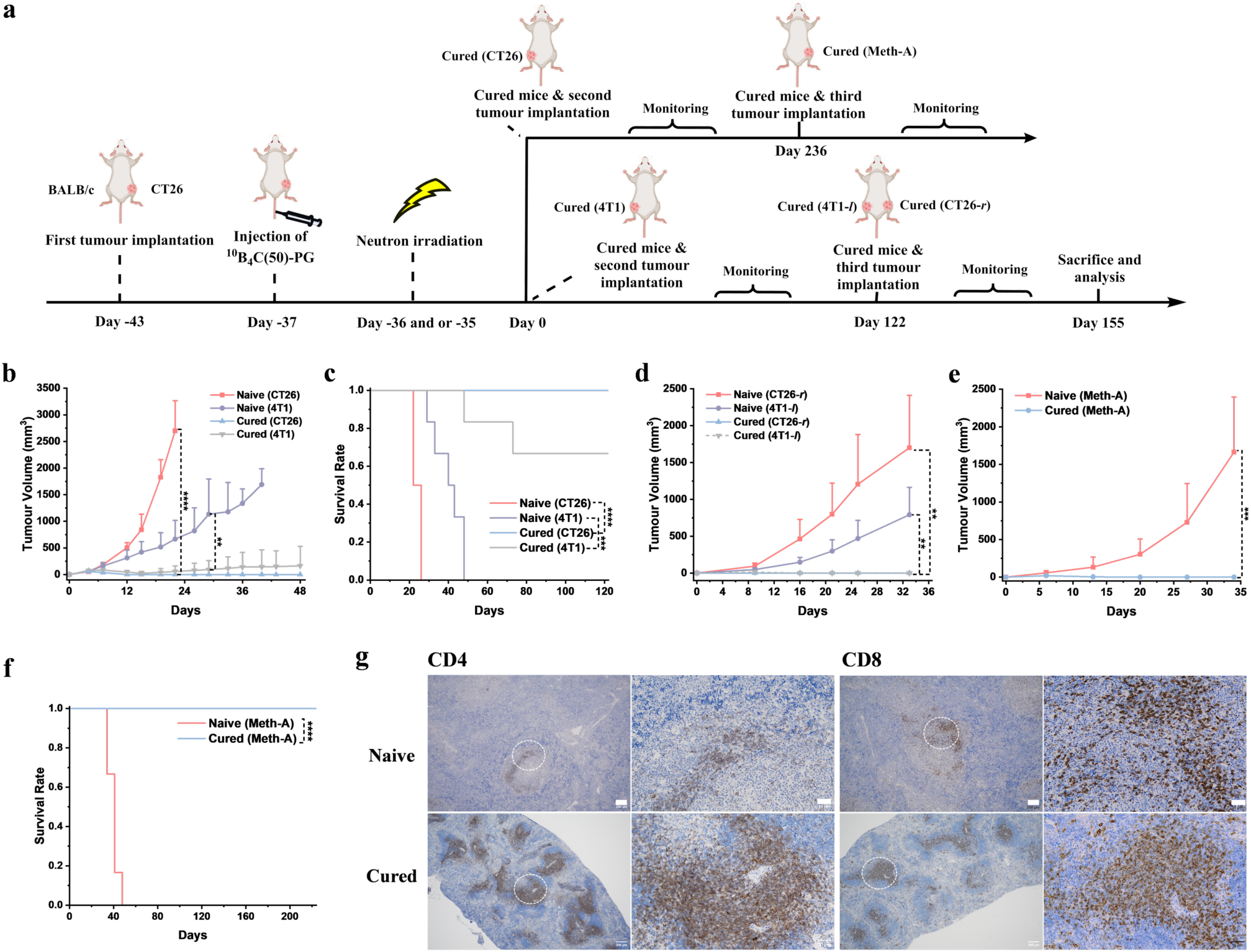
Immune memory induced by ^10^B_4_C(50)-PG–BNCT. **a**, Schematic of various tumour rechallenge processes in mice with CT26 tumour in right thigh cured by ^10^B_4_C(50)-PG–BNCT. **b**, CT26 and 4T1 tumour volumes in CT26 cured mice rechallenged by CT26 and 4T1, Cured (CT26) and Cured (4T1), respectively, in addition to CT26 and 4T1 implanted mice, Naive (CT26) and Naive (4T1), respectively (*n* = 6), monitored for 48 days. **c**, Survival rate of Naive (CT26), Naive (4T1), Cured (CT26) and Cured (4T1) for 120 days. **d**, Volumes of CT26 (Naive (CT26-*r*) and Cured (CT26-*r*)) and 4T1 (Naive (4T1-*l*) and Cured (4T1-*l*)) tumours in right and left thighs, respectively, in Naive (CT26 + 4T1) and Cured (CT26 + 4T1) mice monitored for 33 days, after their simultaneous implantation in Cured (4T1) rejecting 4T1 rechallenge completely (*n* = 4). **e**, Meth-A tumour volumes monitored for 34 days at rechallenge (Cured (Meth-A)) in Cured (CT26) rejecting CT26 at rechallenge experiment (*n* = 6). **f**, Survival rate of Cured (Meth-A) in comparison with Naive (Meth-A) where Meth-A was implanted in untreated mice. **g**, Typical images of spleen sections in Naive (CT26 + 4T1) and Cured (CT26 + 4T1) mice by immunohistochemical staining after 33 days post both CT26 and 4T1 implantation (Scale bar, 200 µm for the left image and 50 µm for magnified image). Statistical analysis of relative tumour volume with one-way ANOVA post Bonferroni test; \**p* < 0.05, \*\**p* < 0.01, \*\*\**p* < 0.001, \*\*\*\**p* < 0.0001.

To further examine the persistence of the immune memory, both CT26 and 4T1 cells were implanted simultaneously at the right and left thighs respectively (Cured (CT26 + 4T1) mice) in the Cured (4T1) mice rejecting 4T1 cells completely after 122 days post first 4T1 implantation (Fig. 4a). Neither of the cells in the Cured (CT26 + 4T1) mice grew at all in the right and left thighs (Cured (CT26-*r*) and Cured (4T1-*l*), respectively) (Fig. 4d and Supplementary Fig. 46). In addition, all the Cured (CT26 + 4T1) mice kept healthy with little inflammation based on the low IFN-γ and undetected TNF-α levels (Supplementary Fig. 47), and with no metastases in lung tissue in contrast to Naive (CT26 + 4T1) ones based on the H&E staining (Supplementary Fig. 48). For further applicability, Cured (CT26) mice rejecting CT26 in the left thigh recognized Meth-A cells immunologically not to engraft at the right thigh in the Cured (Meth-A) mice (Fig. 4e), which survived over 220 days after the Meth-A implantation (Fig. 4f and Supplementary Fig. 49). These results indicate that the immunity acquired by ^10^B_4_C(50)-PG–BNCT prevents the recurrence of the same cancer completely and also resists to the other types of murine cancer. In all the experiments, normal body weight increment was observed in the Cured (CT26) and Cured (4T1) even with additional implantation of CT26, 4T1 and Meth-A (Supplementary Fig. 50).

Following direct DNA damage in tumour cells caused by ^10^B_4_C(50)-PG–BNCT^26^, the antigens from the damaged tumour cells may be displayed on the antigen-presenting cells (APCs) like dendritic cells^40^. The APCs then activate the naive T cells through cross-presentation of the antigens in nearby lymphoid organs and the activated T cells infiltrate the tumour. Importantly, memory T cells form and persist even after pathogen clearance, circulating in the bloodstream and/or residing in tissues for long time^41^. To investigate this mechanism, the CD4^+^ and CD8^+^ T cell populations in the spleen and lymph node were evaluated in the Naive (CT26 + 4T1) and Cured (CT26 + 4T1) mice by immunohistostaining after 33 days post simultaneous CT26 and 4T1 implantation. Both CD4 and CD8 expressions were higher in the spleen of Cured (CT26 + 4T1) mice compared to Naive (CT26 + 4T1) mice, while CD4 expression was higher in the lymph nodes (Fig. 4g and Supplementary Fig. 51). These are probably memory T cells primed for rapid and effective responses upon re-exposure to tumour antigens, producing effector cytokines and cytolytic mediators^41^. Given that tumour-specific antigens, such as AH1, are shared across different murine tumour types^42,43^, these memory T cells are considered to not only recognize the CT26 cells, but also the other murine tumour cells.

## Long term *in vivo* safety assessment of ^10^B4C(50)-PG

Finally, to evaluate the biosafety of ^10^B_4_C(50)-PG, body weight and food consumption were monitored in the mice administered by PBS and ^10^B_4_C(50)-PG for one year (Fig. 5a and Supplementary Fig. 52). In addition, major blood cell populations and blood chemistry were analyzed after 381 days post-administration (Fig. 5b – 5d). All the differences were negligible in the body weight, food consumption, blood cell population and blood chemistry. Furthermore, structural integrity of major organs and muscle was confirmed by H&E staining after one year (Fig. 5e). To assess long term *in vivo* behaviour of ^10^B_4_C(50)-PG, biodistribution was compared in the mice after 14 and 381 days post administration (Fig. 5f, 5g). Both [^10^B] and %ID in liver and spleen decreased from 43 ppm and 38.5% to 16 ppm and 20.0%, and from 88 ppm and 9.9% to 49 ppm and 8.1%, respectively. These results indicate that ^10^B_4_C(50)-PG is gradually excreted from these organs; especially from liver probably through hepatobiliary^44^, and is harmless at this dosage at least for a year. Since ^10^B_4_C(50)-PG consists solely of main group elements with a rounded shape in addition to the small dosage mentioned above, we can avoid the typical concerns of inorganic nanomaterials such as heavy metal and asbestos-like toxicities even in terms of long term safety^1^.

**Fig. 5.**
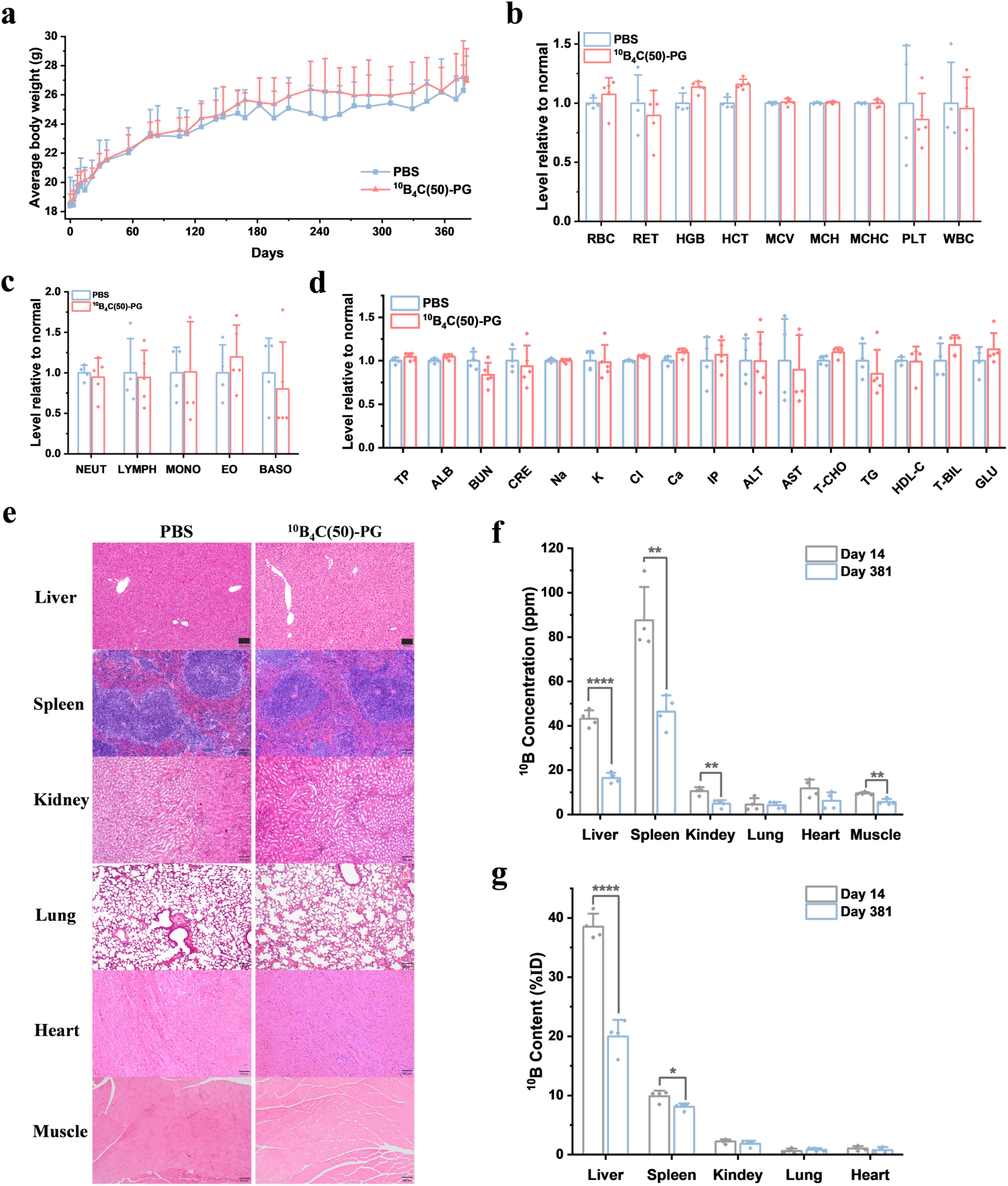
Biosafety of ^10^B_4_C(50)-PG. **a**, Time course of average body weights monitored for 381 days after intravenous injection of PBS (*n* = 4) and ^10^B_4_C(50)-PG at a dosage of 5.2 mg [^10^B]/kg (mouse)] (*n* = 5). **b**, Blood cell levels of red blood cell (RBC), reticulocyte (RET), hemoglobin (HGB), hematocrit (HCT), mean corpuscular volume (MCV), mean corpuscular hemoglobin (MCH), mean corpuscular hemoglobin concentration (MCHC), platelet (PLT), and white blood cell (WBC) after 381 days post injection. **c**, White blood cell levels of neutrophils (NEUT), lymphocytes (LYMPH), monocytes (MONO), eosinophils (EO), and basophils (BASO) after 381 days post injection. **d**, Comprehensive blood chemistry panel including total protein (TP), albumin (ALB), blood urea nitrogen (BUN), creatinine (CRE), sodium (Na), potassium (K), chloride (Cl), calcium (Ca), inorganic phosphorus (IP), alanine aminotransferase (ALT), aspartate aminotransferase (AST), total cholesterol (T-CHO), triglyceride (TG), high-density lipoprotein cholesterol (HDL-C), total bilirubin (T-BIL), and glucose (GLU) after 381 days post injection (*n* = 4 in PBS and *n* = 5 in ^10^B_4_C(50)-PG). **e**, H&E staining of histology sections from major organs and muscle after 381 days post injection. scale bar; 100 µm. **f**, **g** [^10^B] and %ID in major organs and muscle after 14 and 381 days post-injection of ^10^B_4_C(50)-PG at 5.2 mg [^10^B]/kg (mouse)] (*n* = 4). Statistical analysis with one-way ANOVA post Bonferroni test; \**p* < 0.05, \*\**p* < 0.01, \*\*\**p* < 0.001, \*\*\*\**p* < 0.0001.

## Outlook

Although mechanochemical synthesis has been attracting considerable interest from the viewpoint of “green chemistry”, it is still in the infant stage for the synthesis of pharmaceuticals^29,30^. In this study, size-controlled synthesis of ^10^B_4_C NPs has been realized by mechanochemical process in solid phase, which is advantageous ecologically and economically in terms of scalability and quality control in view of clinical application. This is one of the characteristics of the inorganic NP based drugs, which can be differentiated clearly from conventional small molecular drugs synthesized through several steps of typical organic processes. In the following step, since the facile process of the PG functionalization on nanodiamond was reported by our group in 2011, we have revealed various advantages of the PG-coated NPs not only from viewpoint of chemistry such as scalability, applicability and extensibility^7,45^, but also biomedical viewpoints such as high protein-corona shielding efficiency resulting in high stealth efficiency and long blood circulation^18,46^. Although some challenges still remain, we believe that the good manufacturing practice process of ^10^B_4_C(50)-PG can be established for its clinical use.

In BNCT, the accelerator-based neutron generator has been developed to improve its prospect in clinical practice, because they can be installed in hospitals, not like nuclear reactor. At present, 29 facilities installed or under construction and development all over the world including Japan, China, Italy, South Korea, Finland, Spain, Russia, UK, Israel, and Argentina. Among them, the 17 facilities in hospitals and centres are for clinical purpose, and two Japanese hospitals have already started patient treatment covered by insurance^47^. Since the ^10^B drug and the neutron generator work together inseparably for BNCT, the novel nanodrug ^10^B_4_C(50)-PG presented here can make a great progress in BNCT and even radiotherapy. We believe that ^10^B_4_C(50)-PG–BNCT can make a paradigm shift in cancer therapy due to the immune activation to induce abscopal effect and immune memory for metastatic and recurrent tumours, respectively.

## Methods

### Chemicals and materials

H_3_^10^BO_3_ (95% ^10^B) was purchased from Sigma-Aldrich. Magnesium (210 - 841 μm), 35 wt% hydrochloric acid, sodium chloride and 4% paraformaldehyde were purchased from Nacalai Tesque Inc., Japan. Graphite (4 μm), glycidol and CCK-8 kit were purchased from Ito Graphite Co., Ltd., Kanto Chemical Co., Inc. and Dojindo Molecular Technologies, respectively. PBS (−) buffer, RPMI-1640 with L-glutamine and phenol red 0.25w/v%, trypsin-1mmol/L EDTA ·4Na solution with phenol red and 10% neutral buffered formalin were purchased from Wako Pure Chemical, Inc., Japan. InVivoMAb anti-mouse PD-1 (CD279) Clone: RMP1-14 (BioXCell) was purchased from Wako Pure Chemical, Inc., Japan. Primary antibodies were CD3 (CST: #99940), CD4 (CST: #25229) and CD8 (Abcam: ab217344).

### Equipment

Dehydration of H_3_^10^BO_3_ was conducted in AF-2 type heating mantle (Taika Corporation) under a temperature controller TR-KN (As One Corporation). Ball milling synthesis was performed on Pulverisette 7 (Fritsch). Stainless steel 304 milling vials (45 and 12 cm^3^) were purchased from Changsha Mitr Instrument Equipment Co., Ltd. Stainless 304 steel balls (10 and 5 mm) were obtained from Ohashi Steelballs Co., Ltd. FTIR measurements were conducted on IRSpirit (Shimadzu Co. Ltd.), and 32 scans of the spectra were collected with a resolution of 4 cm^−1^ in the wavenumber range of 4000 to 400 cm^−1^ for each sample. TGA was run from room temperature to 1000 °C at a heating rate of 20 °C/min under nitrogen atmosphere on a Q-50 analyzer (TA Instruments). DLS analysis was performed five times for each sample on a Nanotrac UPA-UT151 system (Microtrac, Inc.) in number mode. The zeta potentials were measured 20 times for each sample on a Zetasizer Nano Series (Malvern Instruments, UK). Powder X-ray diffraction (XRD) measurements were conducted with a Rigaku rint2200u/pc-lh diffractometer with CuKα (λ = 1.5418 Å). Scanning electron microscopy analysis and energy dispersive X-ray measurements were carried out on SU8220 (Hitachi, Japan). Raman spectra were recorded by LabRam HR800 (Horiba) on four different spots for the final curve with 488 nm laser. Centrifugation was conducted on CF15RN (Himac) and Avanti J-E (Beckman). The absorbance was recorded at 570 nm on a microtiter plate reader (MTD-310, Corona Electric Co., Japan). The immunohistochemistry slides were examined by Caltivation Microscopy CKX53 Degital Camera DP23 (Olympus).

### Synthesis of ^10^B_4_C

^10^B_2_O_3_ was obtained from the dehydration of ^10^H_3_BO_3_ (4.0 g) at 130 °C for 40 min and 330 °C for 80 min according the method reported before^48^. Then, magnesium (2.2 g) was pretreated by ball milling at ball to powder ratio (BPR) of 10 and rotation speed (RS) at 800 rpm for 30 min with five 10 mm new balls in the container (12 mL). To obtain ^10^B_4_C(35) NP, ^10^B_2_O_3_ (1.0 g) was pretreated at BPR of 20 and RS at 800 rpm for 20 min with four 10 mm new balls in the 12 mL container. Then, after pausing for 5 min, the agglomerated powder was scraped off on the bottom or top of the vial and then same milling was repeated for two more times. To obtain ^10^B_4_C(50) NP, ^10^B_2_O_3_ was pretreated at BPR of 20 and RS of 800 rpm for 20 min with 10 mm new balls (4 balls) in the 12 mL container. For ^10^B_4_C(80), there was no pretreatment to ^10^B_2_O_3_. After that, ^10^B_2_O_3_, magnesium and graphite at mass ratio of 10:11:1, were mill at BPR of 30 and rotation speed of 800 rpm for 8 h with 10 and 5 mm new balls (mass ratio between them was 4) in the 45 mL container to obtain ^10^B_4_C with size of 35, 50 and 80 nm. To synthesize ^10^B_4_C with size above 100 nm, there was no pretreatment to ^10^B_2_O_3_ and the procedure was like former except that the balls were worn which were used for many times. Or just decreasing the milling time from 8 to 6 h could also obtain B_4_C with size of 100 nm (Supplementary Table. 3 and Fig. 3). Then, after 8 h milling finished, NaCl (mass ratio between NaCl and total powder was 5:1) was added into containers and mill for one more hour under the same condition to deagglomerate the powders. Then, leaching process was carried out in 15 wt% hydrochloric acid (40 mL) for 3 h at 90 ℃ with a magnetic stirrer. After that, the dispersion was firstly washed with Milli-Q water by centrifugation at 22260*g* and 25 ℃ for 50 min (Himac). Then after redispersion, the dispersion was secondly washed with Milli-Q water by centrifugation at 50400*g* and 25 ℃ for 30 min (Beckman) by repeated redispersion/ centrifugation cycles until the supernatant became clear and the pH value was 7.

### Synthesis of ^10^B_4_C-PG

^10^B_4_C NPs with different core size were functionalized with PG following the reported procedure^22^. ^10^B_4_C NPs (90 mg) with core size of 35, 50 and 80 nm were once reacted with glycidol (10 mL) while ^10^B_4_C (110) NPs were twice (20 h for first time and 4 h for second time) with magnetic stirrer at 140 ° C. After PG functionalization, the dispersion of ^10^B_4_C with core size of 50, 80 and 110 nm were washed by Milli-Q water for 4 times at conditions of 50400*g* and 25 ℃ for 30 min (Beckman) by repeated redispersion/ centrifugation until the supernatant became transparent, while ^10^B_4_C(35) was washed under the same conditions except that the time increased to 40 min. To obtain ^10^B_4_C(5)-PG NPs, ^10^B_4_C(35)-PG NPs dispersed in water were centrifugated at 15000*g* (Himac) for 10 min at 25 °C in 2 mL centrifugation tube and the supernatant was collected.

### In vitro cytotoxicity studies

CT26 cell line were cultivated in RPMI cell culture media supplemented with 10% fetal bovine serum and 1% of antibiotics at 37°C in a 5% CO_2_ humidified incubator. Cells were used from passage 4 to 30. For viability experiments, CT26 cells were seeded in 96-well plates with a density of 1 × 10^4^ cells/well. After 24 h, the cells were washed with PBS buffer, fresh culture medium containing different concentration (30, 60 and 120 ppm) of ^10^B_4_C NPs or ^10^B_4_C-PGs with different core size was added into each well. After incubation for 24 h, the cells were washed with PBS buffer twice and cell viability was measured by CCK-8 kit. The absorbance was recorded at 450 nm on a microtiter plate reader (MTD-310, Corona Electric Co., Japan). Experiments were carried out three times in quadruplicate.

### Animals and Tumour Models

Female BALB/c and C57BL/6 mice aged 8 to 10 weeks were maintained in the KUR animal facility (Kumatori campus, Kyoto University) with free access to water and standard food before injection and irradiation. Approximately CT26 (1.0 × 10^6^), 4T1 (1.0 × 10^6^), Meth-A (1.7 × 10^6^ in BNCT and 1.5 × 10^6^ in immune memory study), B16-F10 (1.0 × 10^6^) or LLC (0.5 × 10^6^) tumour cells suspended in PBS (100 µL) were subcutaneously implanted into the right leg of mice before the injection of the ^10^B_4_C-PG. The mice were kept under specific pathogen-free conditions, handled, and maintained in accordance with the guidelines of the Science Council of Japan. All animal experiments were conducted in accordance with the regulations of the Experimental Animal Center of Kyoto University and the standards approved by the Kyoto University Ethics Committee. The tumour volume of all mice was measured continuously with the same caliper and the same observer throughout the experiment. When the tumour volume reached 3000 mm^3^ or the body weight loss was over 20%, the mice were removed from the experimental group and euthanized. In some cases, this limit has been exceeded the last day of measurement and the mice were immediately euthanized. The tumour volume was calculated and statistically analyzed according to the equations shown below.

### Calculation of tumour volume

The volume of the tumour was calculated based on the following equation.

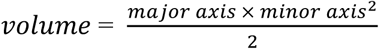

When the tumour volume data were statistically analyzed, all the data were normalized according to the following equation.

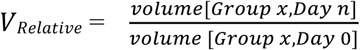

The tumour growth inhibition (TGI) compared to neutron control group (N) was calculated based on the following equation.

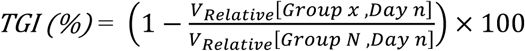

### Size effect of ^10^B_4_C-PG on biodistribution and tumour therapeutic efficacy

For pharmacokinetic studies in CT26 tumour model, female BALB/c mice with average body weight of 22 g (*n* = 3) were i.v. administered with PBS dispersion (200 µL) of ^10^B_4_C-PG with different size at ^10^B dosage of 5.1 mg / kg 24, 48 and 72 h before sacrificing. The mice were euthanized to dissect the organs when the average tumour volume reach around 120 mm^3^. The ^10^B contents were assessed based on the water included in the blood, tumour, and organs. Samples were put into Teflon tubes and ^10^B concentration in each sample was measured by the prompt γ-ray microanalysis (PGRA) system to obtain a ppm-order concentration (µg [^10^B]/g [biomaterial]). The %ID was the measured ^10^B mass of each sample divided by the measured ^10^B mass of the injected dose. The ^10^B mass of the injected dose was calculated from the quantified content of ^10^B in ^10^B_4_C.

For size-effect of NPs *in vivo* antitumour study, female BALB/c mice (*n* = 4 in PBS group and *n* = 5 in the other groups) were i.v. administered with PBS and ^10^B_4_C-PG with different size at ^10^B dosage of 5.1 mg / kg two days before irradiation (*n* = 5) with average weight of 22 g. The neutron was irradiated to the mice with average tumour volume of 140 mm^3^ placed on an acrylic holder, with a circular hole in the center, which is covered with a 5-mm-thick thermoplastic plate containing 40 wt% ^6^LiF (96% ^6^Li) to block thermal neutrons. The thigh with the tumour was stretched over the hole and was irradiated with neutron at flux of 5 × 10^9^ cm^−2^ s^−1^ for 12 min at Kyoto University Research Reactor (KUR). The day when neutron was irradiated was 0. Replicated experiment under the similar conditions was performed (*n* = 3). For demonstrating high dosage of ^10^B_4_C(80)-PG and ^10^B_4_C(110)-PG to eradicate tumour, mice with average weight of 21 g were i.v. administered with PBS or ^10^B_4_C-PG with different size at ^10^B dosage of 21.4 mg / kg two days before irradiation (*n* = 2, in N group and *n* = 3 in the other groups). For demonstrating the therapeutic efficacy of ^10^B_4_C(5)-PG NPs, mice with average body weight of 20 g were i.v. administered with PBS or ^10^B_4_C(5)-PG at ^10^B dosage of 5.2 mg / kg two days before irradiation (*n* = 2 in ^10^B_4_C(5)-PG group and *n* = 3 in the other groups).

### ^10^B_4_C(50)-PG–BNCT in larger or different tumour model

For evaluating the therapeutic efficacy of ^10^B_4_C(50)-PG to larger CT26 tumour in single irradiation, mice were i.v. administered with PBS or ^10^B_4_C(50)-PG at whole material dosage of 12 mg / kg or ^10^B dosage of 5.1 mg / kg two days before irradiation (*n* = 12 in ^10^B_4_C(50)-PG + N group and *n* = 6 in the other groups) when the average weight of mice was 21 g. The mice were irradiated with neutron at flux of 5 × 10^9^ cm^−2^ s^−1^ for 12 min with average tumour volume of 171 mm^3^. In even larger CT26 tumour case, mice (*n* = 4) were processed under the same conditions with average weight of 22 g. The mice were irradiated with neutron at flux of 2 × 10^9^ cm^−2^ s^−1^ for 30 min at day 0 with average tumour volume of 645 mm^3^.

Before performing similar studies to different tumour model, biodistribution studies were carried out. In 4T1 tumour model, female BALB/c mice with average body weight of 20 g (*n* = 3) were i.v. administered with ^10^B_4_C(50)-PG NPs 24, 48 and 72 h before sacrificing at ^10^B dosage of 5.2 mg / kg. In B16-F10 tumour model, female C57BL/6 mice with average body weight of 22 g (*n* = 3) were i.v. administered with ^10^B_4_C(50)-PG NPs 6, 24 and 48 h before sacrificing at ^10^B dosage of 5.1 mg / kg. In LLC tumour model, female C57BL/6 mice with average body weight of 22 g (*n* = 3) were i.v. administered with ^10^B_4_C(50)-PG NPs 48 h before sacrificing at ^10^B dosage of 5.1 mg / kg. The 4T1, B16-F10 and LLC mice were sacrificed to dissect the organs with average tumour volume of 149, 143 and 255 mm^3^, respectively. The ^10^B contents were assessed based on the PGRA system mentioned above.

For evaluating the therapeutic efficacy of ^10^B_4_C(50)-PG to 4T1 (*n* = 3), Meth-A (*n* = 4 in PBS and ^10^B_4_C(50)-PG groups, and *n* = 5 in the other groups), B16-F10 (*n* = 15 in ^10^B_4_C(50)-PG + N group and *n* = 6 in the other groups) and LLC (*n* = 3) tumours in single irradiation, mice were i.v. administered with PBS or ^10^B_4_C(50)-PG at same dosages two days before irradiation with average weight of 17, 20, 20 and 19 g respectively. The mice were irradiated with neutron at flux of 5 × 10^9^ cm^−2^ s^−1^ for 12 min with average 4T1, Meth-A, B16-F10 and LLC tumour volumes of 97, 154, 94 and 71 mm^3^, respectively. The day when neutron was irradiated was 0.

### Neutron fluence and radiation dose quantification

Mice with tumour (*n* = 3) were attached with thermoluminescent dosimeters (TLD) and gold foils on right, left thigh and flank to be irradiated with thermal neutron at flux of 5 × 10^9^ cm^−2^ s^−1^ for 12 min. Then the TLD and gold foils were analyzed to quantify the thermal, epithermal, and fast neutron fluences, and total radiation dose (Gy).

### The abscopal effect in ^10^B_4_C(50)-PG–BNCT

The bilateral CT26 tumour model was established by subcutaneously injecting 1.0 × 10^6^ and 0.5 × 10^6^ cells into the right and left thighs of mice 6 days before irradiation to mimic primary and distant tumours. Female BALB/c mice (*n* = 7) were i.v. administered with PBS and ^10^B_4_C(50)-PG at dosage of 5.1 mg [^10^B]/kg (mouse) two days before irradiation with average weight of 22 g. The mice were irradiated with neutron at flux of 5 × 10^9^ cm^−2^ s^−1^ for 12 min with average tumour volume of 78 and 88 mm^3^ at the right and left thighs. A replicated experiment was performed, with the average body weight at the time of injection being 20 g and the average tumour volumes in the right and left thighs being 58 mm³ and 46 mm³ when irradiating, respectively (*n* = 5). The bilateral 4T1 tumour model was established by subcutaneously injecting 1.0 × 10^6^ cells into the right thigh and flank of mice 6 days and 2 days before irradiation respectively to mimic primary and distant tumours. Mice (*n* = 3) were processed under the same conditions with average weight of 22 g. When the average tumour volume on right and thigh and flank reached 77 and 30 mm^3^ respectively, the mice were irradiated with neutron at same condition.

### Lower dosage of ^10^B_4_C(50)-PG–BNCT with double irradiation

For evaluating the therapeutic efficacy of ^10^B_4_C(50)-PG at lower dosage to CT26 tumour in double irradiation, mice (*n* = 6) were i.v. administered with PBS or ^10^B_4_C(50)-PG at dosage of 2.5 mg [^10^B]/kg (mouse) one day before irradiation (*n* = 5 in N + N group and *n* = 6 in the other groups) with average weight of 21 g. The mice were irradiated with neutron at flux of 5 × 10^9^ cm^−2^ s^−1^ for 12 min on day 0 and 1 respectively with average tumour volume of 166 mm^3^. In another experiment, mice (*n* = 5) were administered with PBS or ^10^B_4_C(50)-PG at the same dosage two day before irradiation when the average weight was 21 g. When the average CT26 tumour volume reached 122 mm^3^, the mice were irradiated with neutron at the same condition on day 0 and 7 respectively.

### The synergistic therapy of ^10^B_4_C(50)-PG NPs–BNCT with anti-PD-1

For evaluating the therapeutic efficacy of ^10^B_4_C(50)-PG to CT26 tumour in combination with anti-PD-1, female BALB/c mice (*n* = 6) were i.v. administered with PBS or ^10^B_4_C(50)-PG at dosage of 2.5 mg [^10^B]/kg (mouse) two day before irradiation when the average weight of mice was 21 g. In 4T1 (*n* = 3), B16-F10 (*n* = 5 in anti-PD-1 group and *n* = 6 in the other groups) and LLC (*n* = 3) tumour models, the mice were processed under the same conditions with average weight of 17, 20 and 19 g respectively. When the average CT26, 4T1, B16-F10 and LLC tumour volume reached 114, 99, 90 and 88 mm^3^, the mice were irradiated with neutron at flux of 5 × 10^9^ cm^−2^ s^−1^ for 12 min. The day when neutron was irradiated was 0. After irradiation, 200 μg of anti-PD-1 was intraperitoneally injected to the mice on day 0, 4, 7 and 11 in CT26 and LLC tumour models, on day 0, 3, 7 and 10 in 4T1 tumour model and on day 3, 6, 9 and 12 in B16-F10 tumour model.

### Immune memory induced by ^10^B_4_C(50)-PG–BNCT

To study the immune memory effect, CT26 and 4T1 cells (1.0 × 10^6^) suspended in PBS were subcutaneously injected into the left thighs of the BALB/c mice (*n* = 6) with CT26 tumour cured by ^10^B_4_C(50)-PG–BNCT in the previous studies. The age-matched naive mice were used as control. Four months later, four mice without tumour in Cured (4T1) group were subcutaneously reimplanted with CT26 and 4T1 cells (1.0 × 10^6^) suspended in PBS in the left and right thighs respectively (*n* = 4). The naive mice with age of 9 weeks were used as control. Six mice without tumour in Cured (CT26) group were subcutaneously implanted with Meth-A cells (1.5 × 10^6^) suspended in PBS in the right thigh (*n* = 6). The naive mice with age of 9 weeks were used as control.

### Immunohistochemistry

CT26 tumour mice were dissected at one day post-irradiation and fixed in 4% paraformaldehyde for over 48 h at 4 °C and then were made into paraffin blocks. The fixed tissues were then sent to the Centre for Anatomical, Pathological and Forensic Medical Researchers, Gradual school of Medicine, Kyoto University to perform staining. The blocks were sectioned at a 3 μm thickness. After deparaffinization and antigen retrieval, endogenous peroxidase activity was blocked by 0.3% H_2_O_2_ in methanol for 30 min. The glass slides were washed in PBS (6 times for 5 min in each washing) and mounted with 1% normal serum in PBS for 30 min. Subsequently primary antibody (CD4 and CD8) was applied overnight at 4 °C. They were incubated with biotinylated second antibody diluted to 1:300 in PBS for 40 min, followed by washing in PBS (6 times, 5 min). Avidin-biotin-peroxidase complex (ABC-Elite, Vector Laboratories, Burlingame, CA) at a dilution of 1:100 in BSA was applied for 50 min. After washing in PBS (6 times, 5 min), coloring reaction was carried out with DAB and nuclei were counterstained with hematoxylin. The slides were examined by Caltivation Microscopy CKX53 Degital Camera DP23 (Olympus). The data were processed and quantified by FIJI.

For the immune memory study, spleen and lymph node form naïve and cured mice were dissected and fixed in 4% paraformaldehyde for over 48 h at 4 ℃. The CD4 and CD8 staining method was the same as that mentioned above.

### *In vivo* safety studies

For the long-term biocompatibility study *in vivo*, female BALB/c mice at age of 6 weeks were i.v. administered with PBS and ^10^B_4_C(50)-PG at dosage of 12 mg / kg (*n* = 4 in PBS group and *n* = 5 in ^10^B_4_C(50)-PG group). The body weight and food consumption of mice in each week were monitored for over one year. At day 381, 0.5 mL of whole blood was collected into tubes with K_2_EDTA (BD Microtainer). To evaluate serum chemistry, 0.5 mL of blood was collected without an anticoagulant and allowed to clot for 30 min at room temperature before centrifuging at 2000*g* (Himac) for 10 min to isolate the serum at 4 °C. All analyses were performed by Oriental Yeast Co., Ltd.

### Histological tissue analysis

Sections of liver, spleen, kidney, lung, heart, muscle and tumour were fixed in 10% neutral buffered formalin for over 48 h. The fixed tissues were then sent to the Centre for Anatomical, Pathological and Forensic Medical Researches, Gradual school of Medicine, Kyoto University to perform H&E staining.

### Quantification and statistical analysis

Statistical analyses were performed using Origin 2024 software (OriginLab Corporation). Statistical parameters are expressed as the mean ± standard error of the mean in figures. No samples nor animals were excluded from data analyses except for death. One-way analysis of variance (ANOVA) was used to analyze data as indicated. Statistical values are indicated according to the following scale: \**p* < 0.05, \*\**p* < 0.01, \*\*\**p* < 0.001, and \*\*\*\**p* < 0.0001.

## Data availability

The data supporting the findings of this study are available within the paper and its Supplementary Information. The raw and analyzed datasets generated during the study are available for research purposes from the corresponding author on reasonable request. Source data are provided with this paper.

## Acknowledgements

The authors acknowledge Dr. Noriko Tsuda (Kyoto University) for experimental help of *in vivo* experiments, and Dr. Hiroki Takahashi, Dr. Guoqing Cheng and Ms. Xinyi Fu (Kyoto University) for guidance of serval equipment. We are also grateful to Dr. Hiroyuki Ueno (Kyoto University Innovation Capital Co., Ltd.) for valuable suggestion in rechallenge experiments, Dr. Jean-Marc Leveque (University of Savoie Mont-Blanc, France) for helpful discussion of ball-milling synthesis, and Dr. Masatoshi Chiba and Dr. Hiroshi Kawai (RadioNano Therapeutics Inc.) for proof-reading the manuscript. We also acknowledge Centre for Anatomical, Pathological and Forensic Medical Researchers, Gradual school of Medicine, Kyoto University to perform Immunohistochemistry, and H&E staining. This work is funded by Japan Society for the Promotion of Science (JSPS, Project No. 23KJ1245, 23K26509 and 21K19906) and JST Fund program for creating research-based startups from academia (JPMJSF23AV). K.H.G., J.Y. and X.C. acknowledge JST SPRING (ID No. JPMJSP2110), JSPS DC2 (ID No. 23KJ1245) and JSPS postdoctoral fellowships (ID No. P21705), respectively.

## Author contributions

N.K., M.S. and W.H. conceptualized the project. W.H. designed and performed the nanoparticle synthesis and characterization. W.H. and K.H.G. performed the cell experiments. W.H., K.H.G., J.Y., X.H., X.C., Y.S. and T.T. performed the biodistribution experiments. W.H, K.H.G, X.H, J.Y, M.S., X.C., Y.S. and T.T. performed the BNCT experiments. W.H., K.H.G. and X.H. performed long-term immune memory studies. W.H., X.H., K.H.G. and J.Y. performed the biocompatibility study of nanoparticles *in vivo*. W.H. performed the data analysis. W.H. and N.K. wrote the initial manuscript. All authors contributed to editing and revising the manuscript.

## Additional information

**Extended Data Fig. 1.**
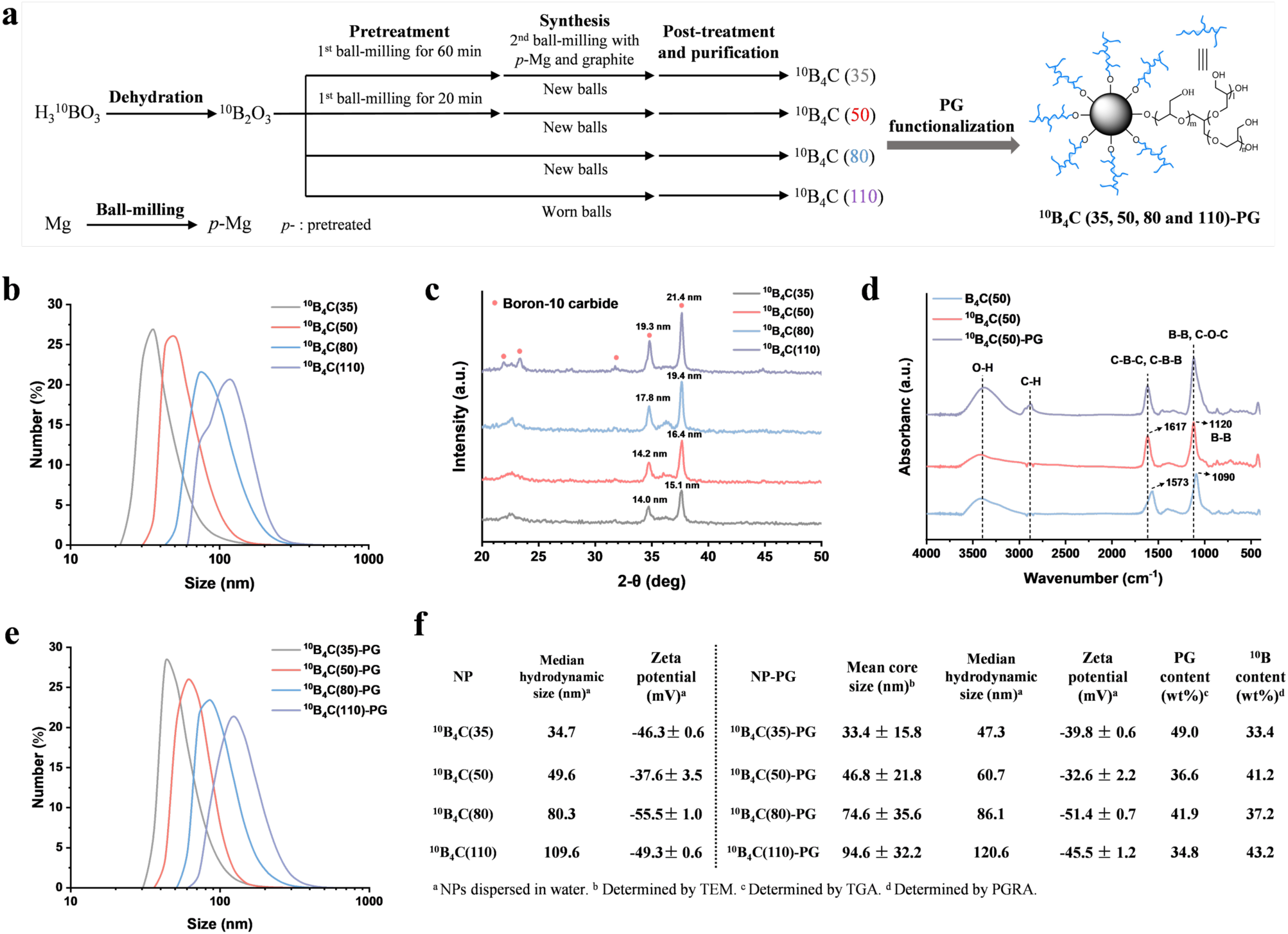
Synthesis of ^10^B_4_C NPs and ^10^B_4_C-PG, and their characterization. **a,** Schematic of procedure to synthesize ^10^B_4_C(*Y*) NPs and ^10^B_4_C(*Y*)-PG with core sizes *Y* = 35, 50, 80 and 110 nm. **b,** Powder XRD profiles of ^10^B_4_C(*Y*) NPs. **c,** FTIR spectra of ^10^B_4_C(*Y*) NPs **d,** Hydrodynamic size distribution of ^10^B_4_C(*Y*) NPs dispersed in Milli-Q water (*n* = 5). **e,** FTIR spectra of ^10^B_4_C(*Y*)-PGs. **f,** Hydrodynamic size distribution of ^10^B_4_C(*Y*)-PGs dispersed in PBS (*n* = 5). **g,** TGA curves of ^10^B_4_C(*Y*)-PGs in nitrogen atmosphere. **h,** Summary in characterization of ^10^B_4_C(*Y*) NPs and ^10^B_4_C(*Y*)-PGs.

**Extended Data Fig. 2.**
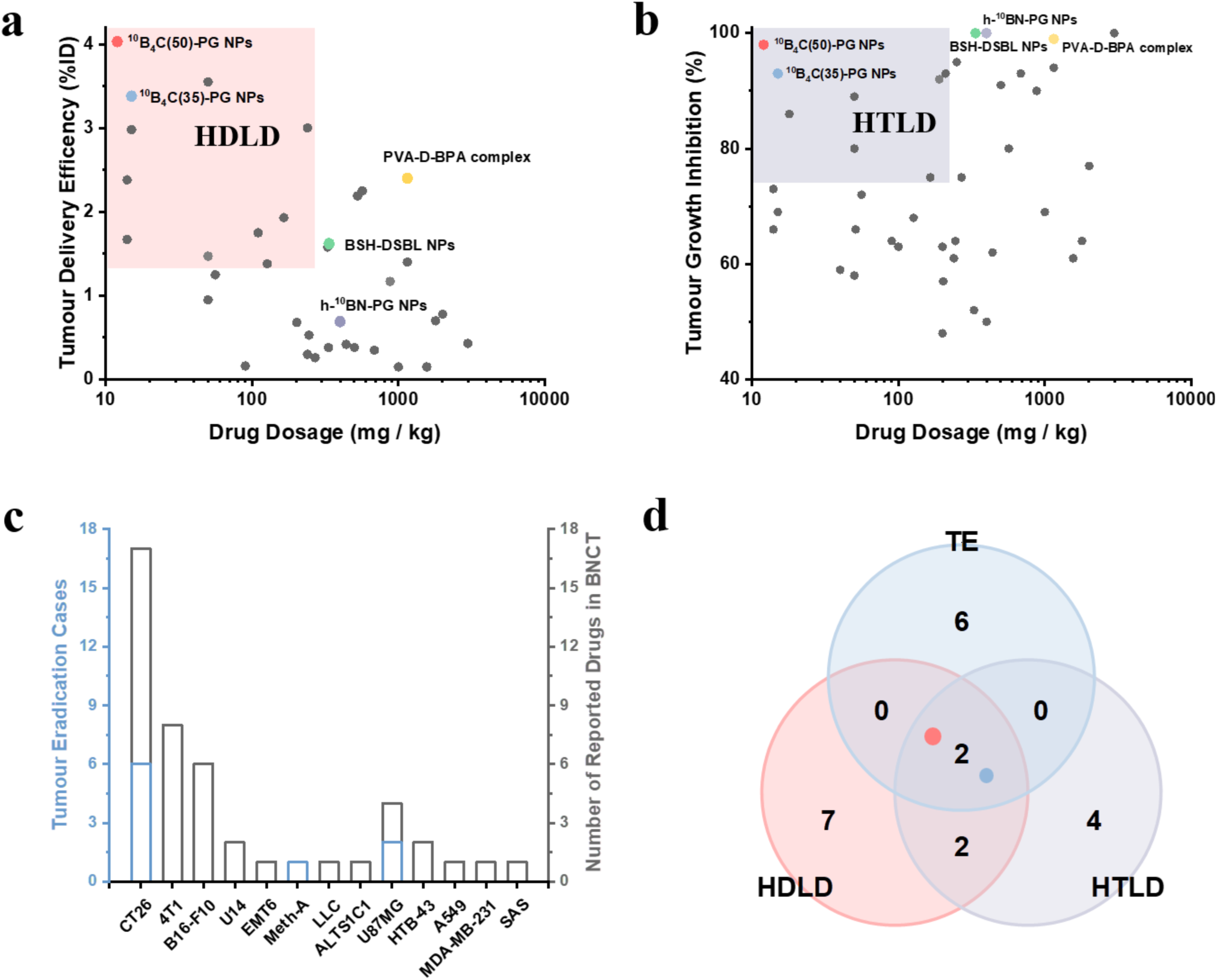
Statistical analysis of drugs used in preclinical BNCT. **a**, Delivery efficacy of drugs to tumour and drug dosage analysis (*n* = 34). Red area represents delivery efficacy of drug to tumour above median value and drug dosage below median value (HDLD). **b**, Neutron therapeutic efficacy to tumour and drug dosage analysis (*n* = 42). Purple area represents neutron therapeutic efficacy to tumour above median value and drug dosage below median value (HTLD). **c**, Tumour eradication cases (more than half of tumours) happened in different tumour models (*n* = 46). Tumour eradication (TE). **d,** drugs analysis from HDLD (*n* = 11), HTLD (*n* = 8) and TE (*n* = 8).

## References

1 Mitchell, M. J. et al. Engineering precision nanoparticles for drug delivery. Nat. Rev. Drug Discov. 20, 101–124 (2021).

2 Barenholz, Y. Doxil^®^--the first FDA-approved nano-drug: Lessons learned. J. Control. Release 160, 117–134 (2012).

3 Von Hoff, D. D. et al. Increased survival in pancreatic cancer with nab-paclitaxel plus gemcitabine. N. Engl. J. Med. 369, 1691–1703 (2013).

4 Anselmo, A. C. & Mitragotri, S. A review of clinical translation of inorganic nanoparticles. AAPS J 17, 1041–1054 (2015).

5 Anselmo, A. C. & Mitragotri, S. Nanoparticles in the clinic: An update. Bioeng. Transl. Med. 4, e10143 (2019).

6 Anu Mary Ealia, S. & Saravanakumar, M. P. A review on the classification, characterisation, synthesis of nanoparticles and their application. IOP Conf. Ser. Mater. Sci. Eng. 263, 032019 (2017).

7 Komatsu, N. Poly(glycerol)-based biomedical nanodevices constructed by functional programming on inorganic nanoparticles for cancer nanomedicine. Acc. Chem. Res. 56, 106–116 (2023).

8 Wilhelm, S. et al. Analysis of nanoparticle delivery to tumours. Nat. Rev. Mater. 1, 16014 (2016).

9 Elsaesser, A. & Howard, C. V. Toxicology of nanoparticles. Adv. Drug Deliv. Rev. 64, 129–137 (2012).

10 Locher G, L. Biological effects of therapeutic possibilities of neutrons. Am. J. Roentgenol. 36, 1 (1936).

11 Suzuki, M. Boron neutron capture therapy (BNCT): A unique role in radiotherapy with a view to entering the accelerator-based BNCT era. Int. J. Clin. Oncol. 25, 43–50 (2020).

12 Lamba, M., Goswami, A. & Bandyopadhyay, A. A periodic development of BPA and BSH based derivatives in boron neutron capture therapy (BNCT). Chem. Commun. (Camb.) 57, 827–839 (2021).

13 Nakashima, H. The new generation of particle therapy focused on boron element (boron neutron capture therapy; BNCT)-the world’s first approved BNCT drug. Yakugaku Zasshi 142, 155–164 (2022).

14 Mohammadpour, R., Yazdimamaghani, M., Cheney, D. L., Jedrzkiewicz, J. & Ghandehari, H. Subchronic toxicity of silica nanoparticles as a function of size and porosity. J. Control. Release 304, 216–232 (2019).

15 Daguenet, E. et al. Radiation-induced bystander and abscopal effects: Important lessons from preclinical models. Br. J. Cancer 123, 339–348 (2020).

16 Deng, S. et al. A PD-L1 siRNA-loaded boron nanoparticle for targeted cancer radiotherapy and immunotherapy. Adv. Mater., e2419418 (2025).

17 Fujimoto, T. et al. Overcoming immunotherapy resistance and inducing abscopal effects with boron neutron immunotherapy (B-NIT). Cancer Sci. 115, 3231–3247 (2024).

18 Zou, Y. et al. Polyglycerol grafting shields nanoparticles from protein corona formation to avoid macrophage uptake. ACS Nano 14, 7216–7226 (2020).

19 Mhadhbi, M. Modelling of the high-energy ball milling process. Adv. Mater. Phy. Chem. 11, 31–44 (2021).

20 Li, X. et al. The dispersion of boron carbide powder in aqueous media. J. Eur. Ceram. Soc. 33, 1655–1663 (2013).

21 Werheit, H., Kuhlmann, U., Rotter, H. W. & Shalamberidze, S. O. Isotopic effects on the phonon modes in boron carbide. J. Phys. Condens. Matter 22, 395401 (2010).

22 Zhao, L., et al. Chromatographic separation of highly soluble diamond nanoparticles prepared by polyglycerol grafting. Angew. Chem. Int. Ed. Engl. 50, 1388–1392 (2011).

23 Zou, Y., Ito, S., Fujiwara, M. & Komatsu, N. Probing the role of charged functional groups on nanoparticles grafted with polyglycerol in protein adsorption and cellular uptake. Adv. Funct. Mater. 32, 2111077 (2022).

24 Zhao, L. et al. Platinum on nanodiamond: A promising prodrug conjugated with stealth polyglycerol, targeting peptide and acid-responsive antitumor drug. Adv. Funct. Mater. 24, 5348–5357 (2014).

25 DeFrancesco, H., Dudley, J. & Coca, A. in Boron reagents in synthesis Vol. 1236 Acs symposium series Ch. 1, 1–25 (American Chemical Society, 2016).

26 Zhang, Y. et al. Tumor eradication by boron neutron capture therapy with ^10^B-enriched hexagonal boron nitride nanoparticles grafted with poly(glycerol). Adv. Mater. 35, e2301479 (2023).

27 Kobayashi, T. & Kanda, K. Microanalysis system of ppm-order ^10^B concentrations in tissue for neutron capture therapy by prompt gamma-ray spectrometry. Nucl. Instrum. Methods Phys. Res. 204, 525–531 (1983).

28 Domnich, V., Reynaud, S., Haber, R. A. & Chhowalla, M. Boron carbide: Structure, properties, and stability under stress. J. Am. Ceram. Soc. 94, 3605–3628 (2011).

29 Kar, S., Sanderson, H., Roy, K., Benfenati, E. & Leszczynski, J. Green chemistry in the synthesis of pharmaceuticals. Chem. Rev. 122, 3637–3710 (2022).

30 Ardila-Fierro, K. J. & Hernandez, J. G. Sustainability assessment of mechanochemistry by using the twelve principles of green chemistry. ChemSusChem 14, 2145–2162 (2021).

31 Nguyen, L. N. M. et al. The exit of nanoparticles from solid tumours. Nat. Mater. (2023).

32 Nomoto, T. et al. Poly(vinyl alcohol) boosting therapeutic potential of *p*-boronophenylalanine in neutron capture therapy by modulating metabolism. Sci. Adv. 6, eaaz1722 (2020).

33 Haraldsson, B., Nystrom, J. & Deen, W. M. Properties of the glomerular barrier and mechanisms of proteinuria. Physiol. Rev. 88, 451–487 (2008).

34 Elmore, S. A. et al. Recommendations from the inhand apoptosis/necrosis working group. Toxicol. Pathol. 44, 173–188 (2016).

35 Tang, L. et al. Investigating the optimal size of anticancer nanomedicine. PNAS 111, 15344–15349 (2014).

36 Janopaul-Naylor, J. R., Shen, Y., Qian, D. C. & Buchwald, Z. S. The abscopal effect: A review of pre-clinical and clinical advances. Int. J. Mol. Sci. 22 (2021).

37 Binnewies, M. et al. Understanding the tumor immune microenvironment (TIME) for effective therapy. Nat. Med. 24, 541–550 (2018).

38 Mahvi, D. A., Liu, R., Grinstaff, M. W., Colson, Y. L. & Raut, C. P. Local cancer recurrence: The realities, challenges, and opportunities for new therapies. CA Cancer J. Clin. 68, 488–505 (2018).

39 Netea, M. G. et al. Defining trained immunity and its role in health and disease. Nat. Rev. Immunol. 20, 375–388 (2020).

40 Moon, C. Y. et al. Dendritic cell maturation in cancer. Nat. Rev. Cancer (2025).

41 Lam, N., Lee, Y. & Farber, D. L. A guide to adaptive immune memory. Nat. Rev. Immunol. 24, 810–829 (2024).

42 Scrimieri, F. et al. Murine leukemia virus envelope GP70 is a shared biomarker for the high-sensitivity quantification of murine tumor burden. Oncoimmunology 2, e26889 (2013).

43 He, X. et al. A potent cancer vaccine adjuvant system for particleization of short, synthetic CD8^+^ T cell epitopes. ACS Nano 15, 4357–4371 (2021).

44 Poon, W. et al. Elimination pathways of nanoparticles. ACS Nano 13, 5785–5798 (2019).

45 Nishikawa, M. et al. Thorough elucidation of synthesis and structure of poly(glycerol) functionalized nanodiamonds. Carbon 205, 463–474 (2023).

46 Yoshino, F. et al. Preferential tumor accumulation of polyglycerol functionalized nanodiamond conjugated with cyanine dye leading to near-infrared fluorescence in vivo tumor imaging. Small 15, e1901930 (2019).

47 INTERNATIONAL ATOMIC ENERGY AGENCY, Advances in Boron Neutron Capture Therapy, Non-serial Publications, IAEA, Vienna (2023).

48 Balcı, S., Sezgi, N. A. & Eren, E. Boron oxide production kinetics using boric acid as raw material. Ind. Eng. Chem. Res. 51, 11091–11096 (2012).

