## supplementary file for "Boosting Antitumour Efficacy and Immunity by Boron Neutron Capture Therapy with Size-Controlled Nanoparticles"

**Supplementary Figures and Tables**

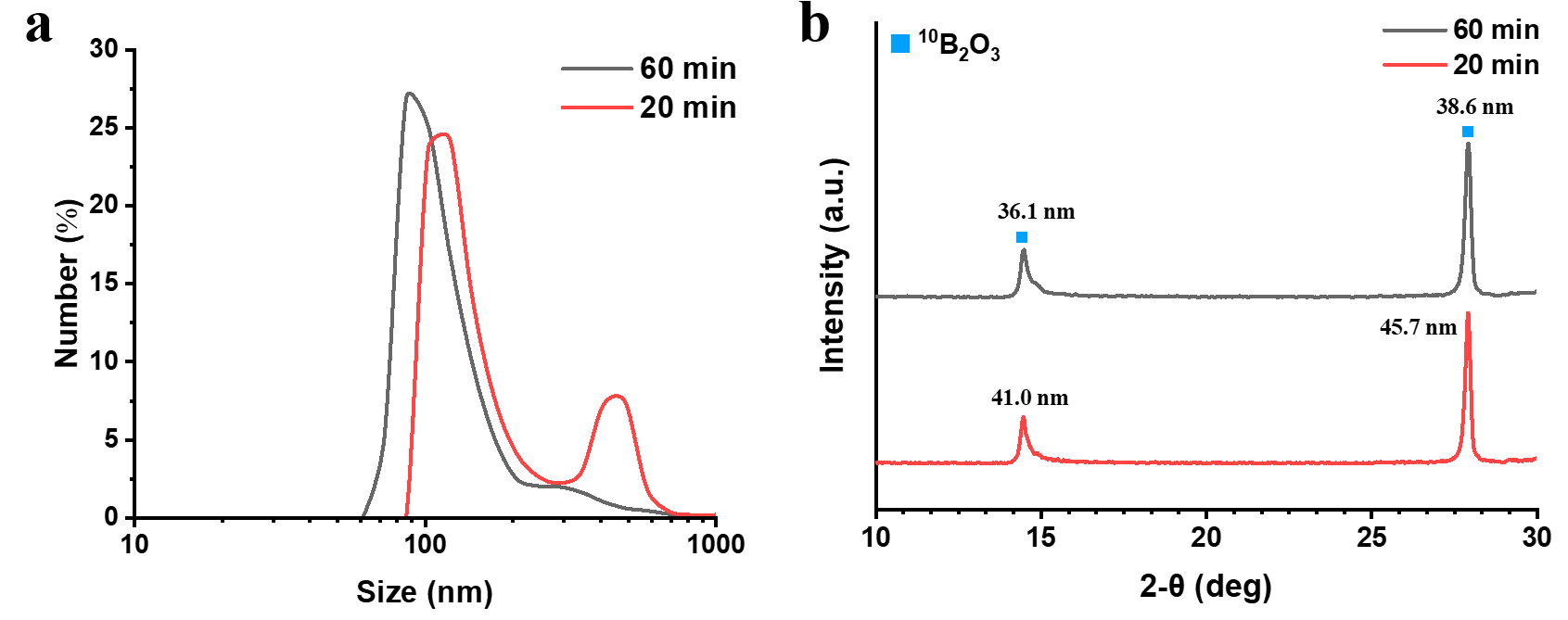

**Supplementary Fig. 1** **|** **Characterization of *p*-^10^B_2_O_3_ NPs. a**, Hydrodynamic size of *p*-^10^B_2_O_3_ NPs in DMSO by dynamic light scattering (DLS) after 20- and 60-min ball-milling (*n* = 5). **b**, Powder X-ray diffraction (XRD) profile of *p*-^10^B_2_O_3_ NPs after 20- and 60-min ball-milling.

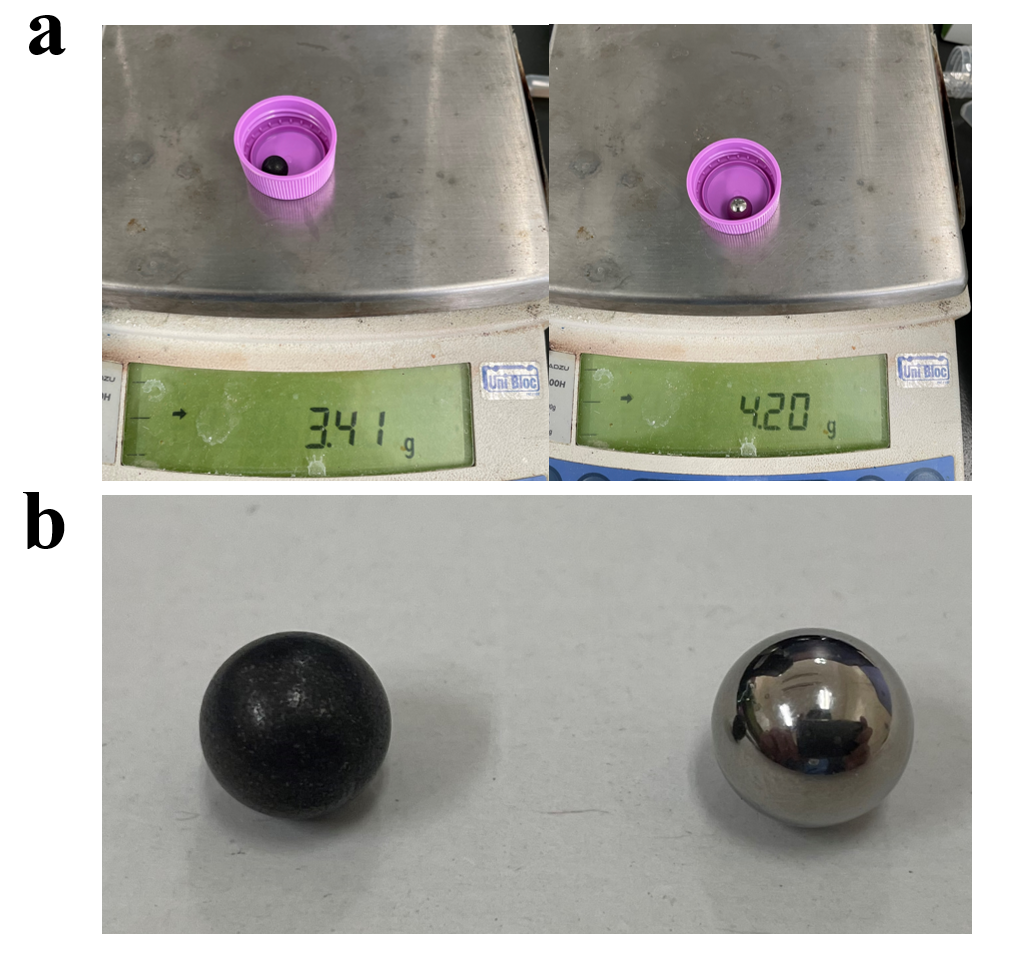

**Supplementary Fig. 2 |** The weight (**a**) and surface (**b**) differences between worn (left) and new (right) balls with 10 mm in diameters.

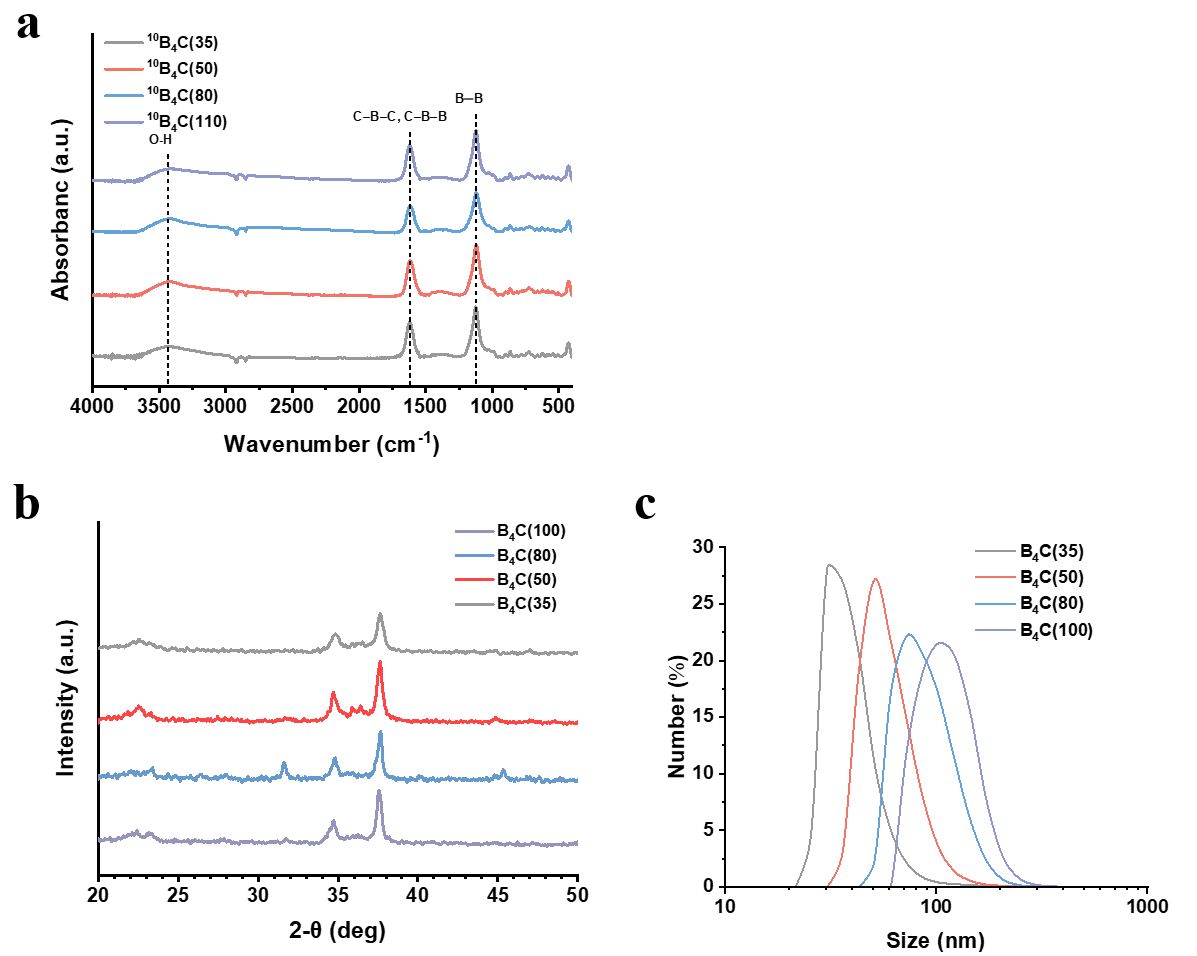

**Supplementary Fig. 3 | Characterization of ^10^B_4_C and B_4_C NPs. a,** FTIR spectra of ^10^B_4_C NPs. **b,** Powder XRD profiles of B_4_C NPs. **c,** Hydrodynamic size of B_4_C NPs dispersed in Milli-Q water, measured by DLS (*n* = 5).

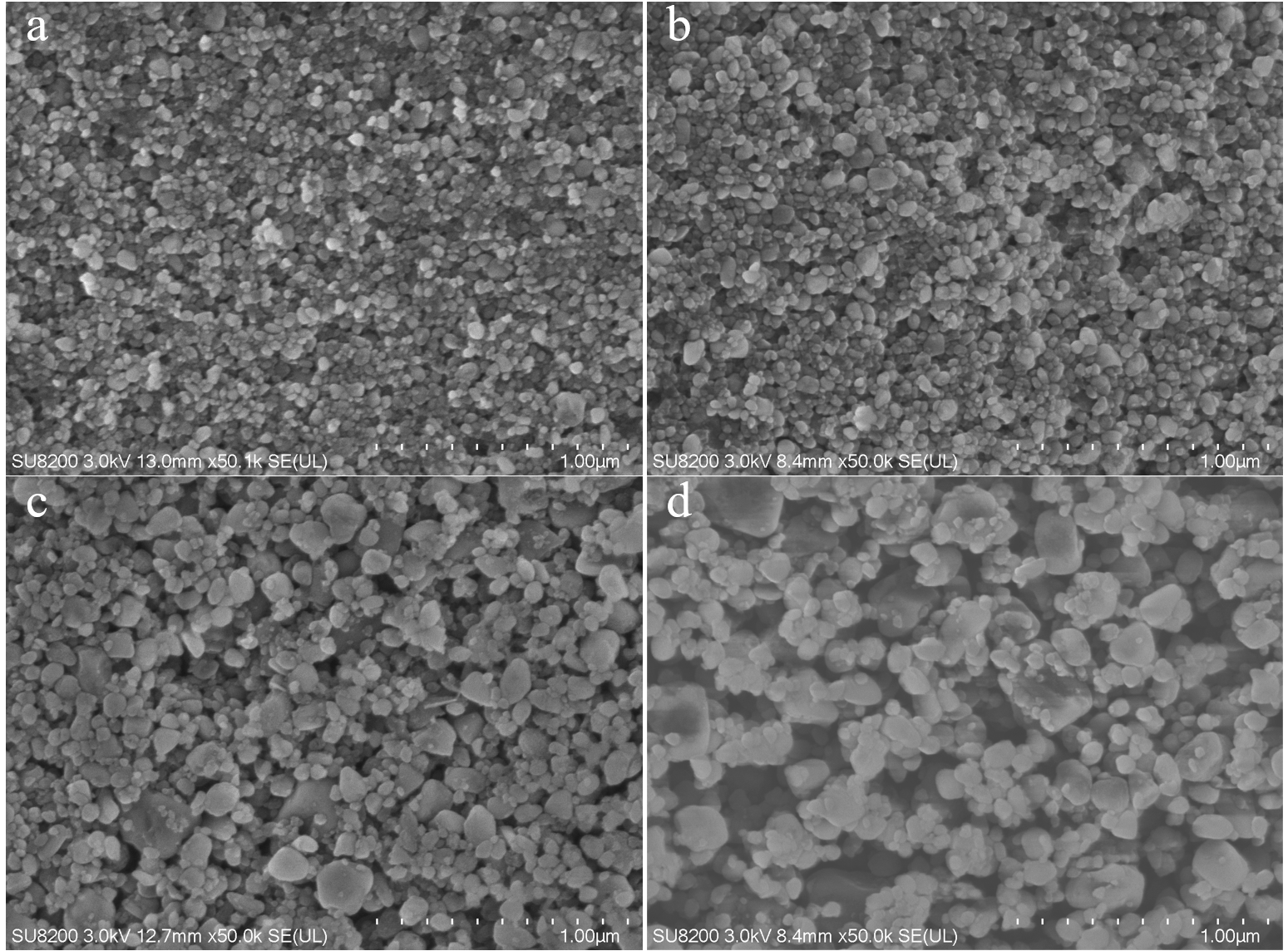

**Supplementary Fig. 4 | SEM images of ^10^B_4_C NPs with different sizes**. **a**, 35 nm. **b**, 50 nm. **c**, 80 nm. **d,** 110 nm. Scale bar = 1.00 μm.

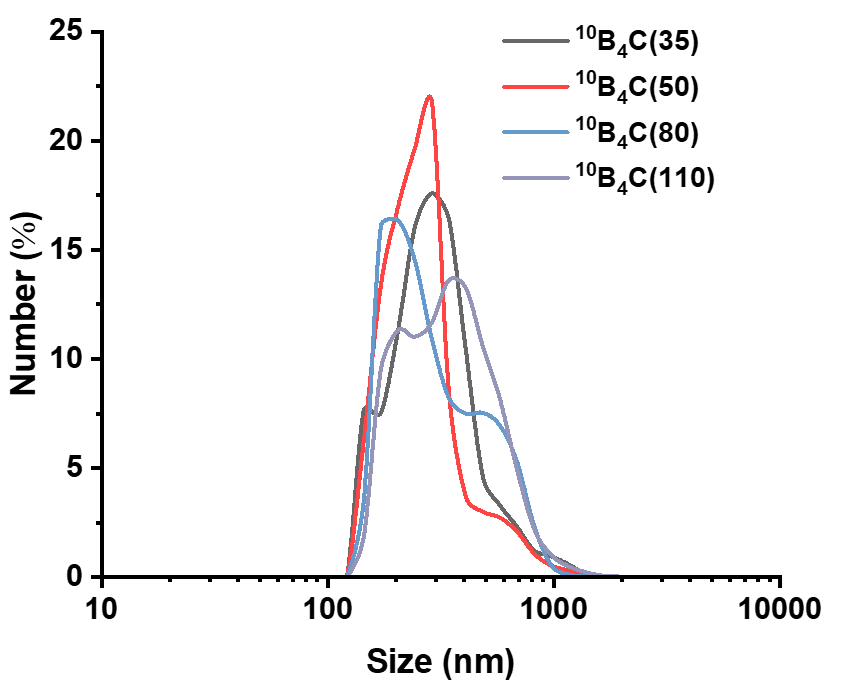

**Supplementary Fig. 5 |** Hydrodynamic size of ^10^B_4_C NPs with different sizes dispersed in PBS (*n* = 5).

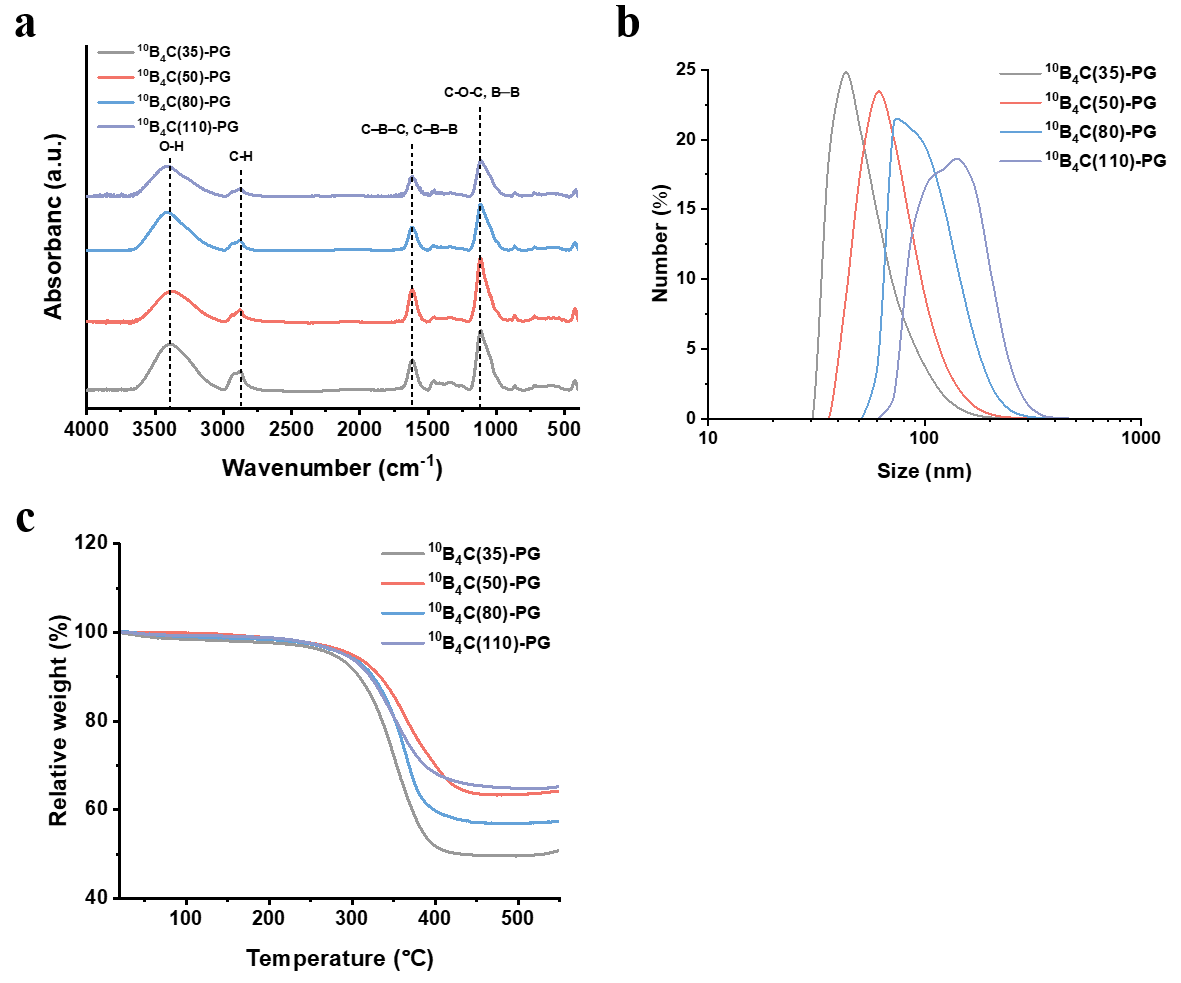

**Supplementary Fig. 6 | Characterization of ^10^B_4_C(*Y*)-PGs. a,** FTIR spectra of ^10^B_4_C(*Y*)-PGs. **b,** Hydrodynamic size distribution of ^10^B_4_C(*Y*)-PG with different sizes dispersed in PBS (*n* = 5). **c,** TGA curves of ^10^B_4_C(*Y*)-PGs with different sizes in nitrogen atmosphere. *Y* = 35, 50, 80 and 110 nm.

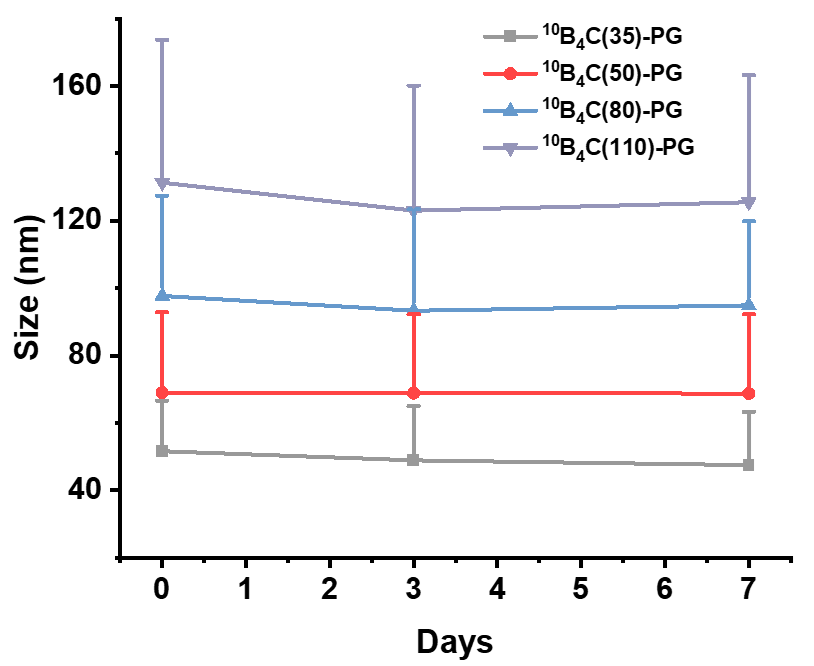

**Supplementary Fig. 7 |** Hydrodynamic size distribution of ^10^B_4_C(*Y*)-PGs (*Y* = 35, 50, 80 and 110) with different sizes dispersed in PBS on 0, 3 and 7 days (*n* = 5).

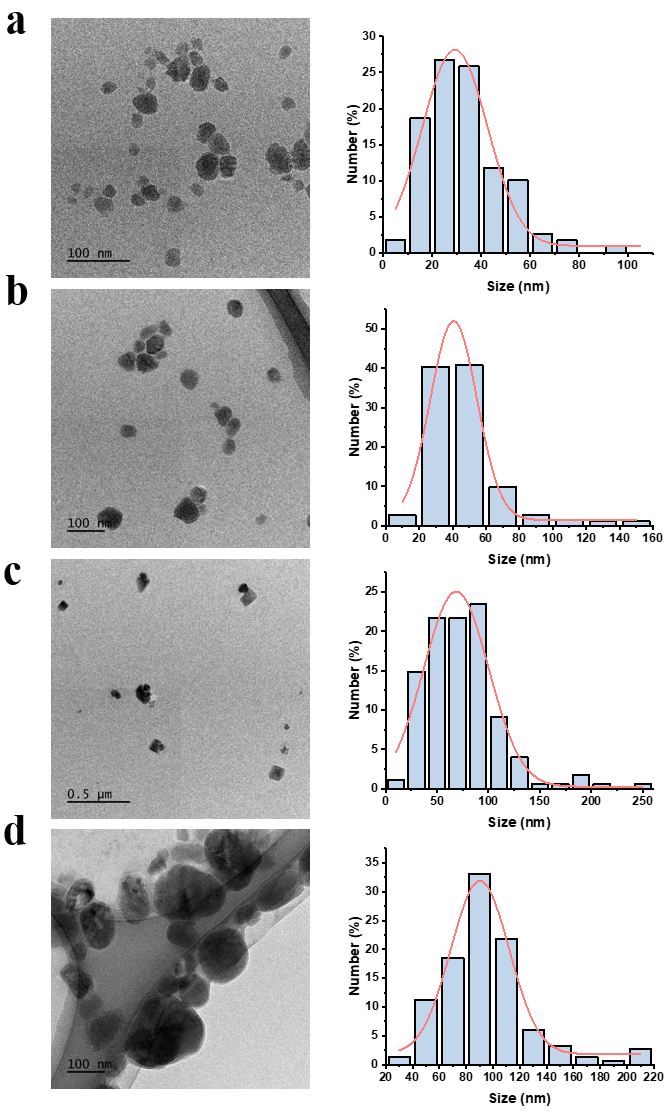

**Supplementary Fig. 8 |** **Core size distribution of ^10^B_4_C-PGs with different sizes observed in TEM.** **a**, ^10^B_4_C(35)-PG (*n* = 348). **b**, ^10^B_4_C(50)-PG (*n* = 267). **c**, ^10^B_4_C(80)-PG (*n* = 175). **d**, ^10^B_4_C(110)-PG (*n* = 151).

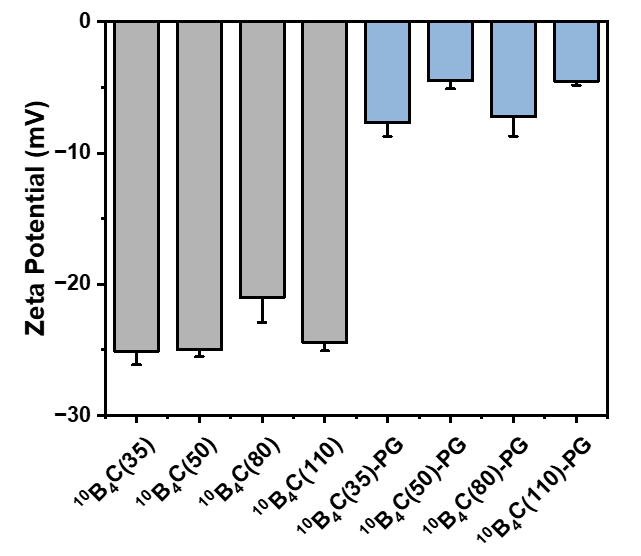

**Supplementary Fig. 9 |** Zeta potentials of ^10^B_4_C NPs and ^10^B_4_C-PGs with different sizes dispersed in PBS (*n* = 3).

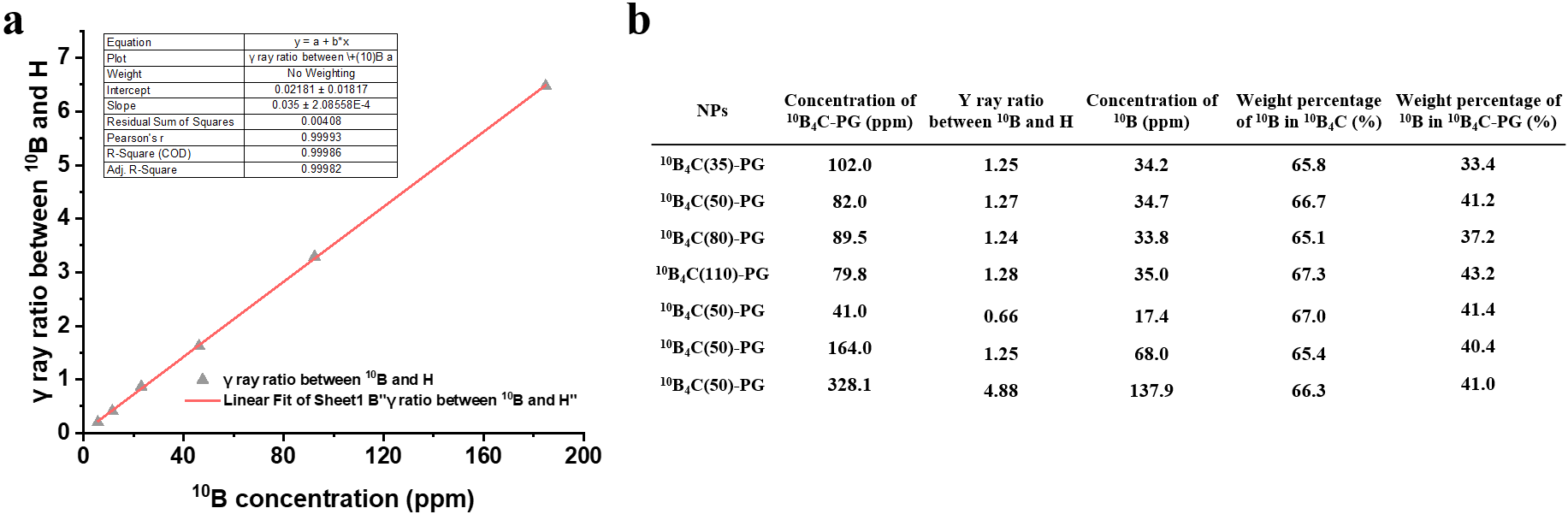

**Supplementary Fig. 10 | ^10^B quantification of ^10^B_4_C-PGs. a**, Calibration curve drawn by standard boric acid solutions for prompt γ-ray microanalysis (PGRA)^1^. **b**, ^10^B content in ^10^B_4_C-PG determined by PGRA. Same ^10^B_4_C(50)-PGs were measured at different concentrations.

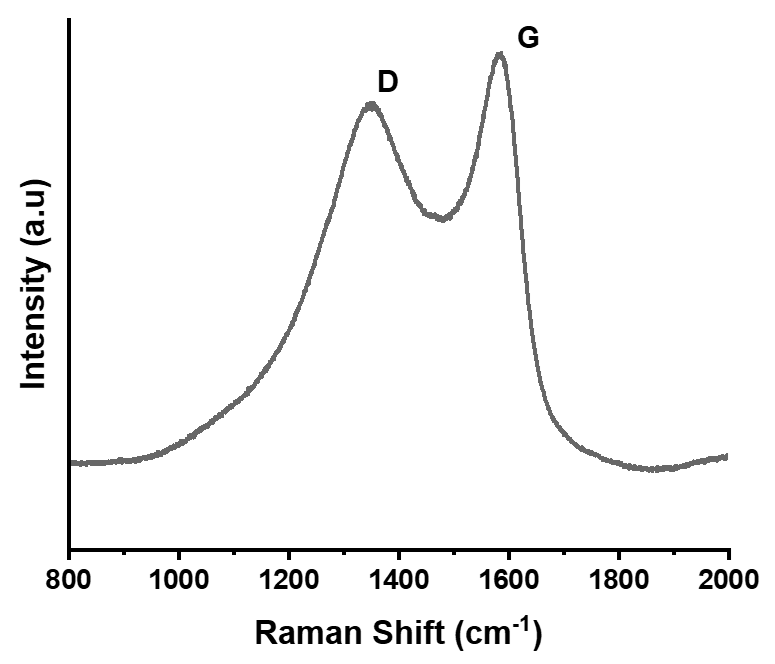

**Supplementary Fig. 11 |** Raman spectra of ^10^B_4_C(50)-PG (*n =* 4). **D** and **G** stand for D and G bands.

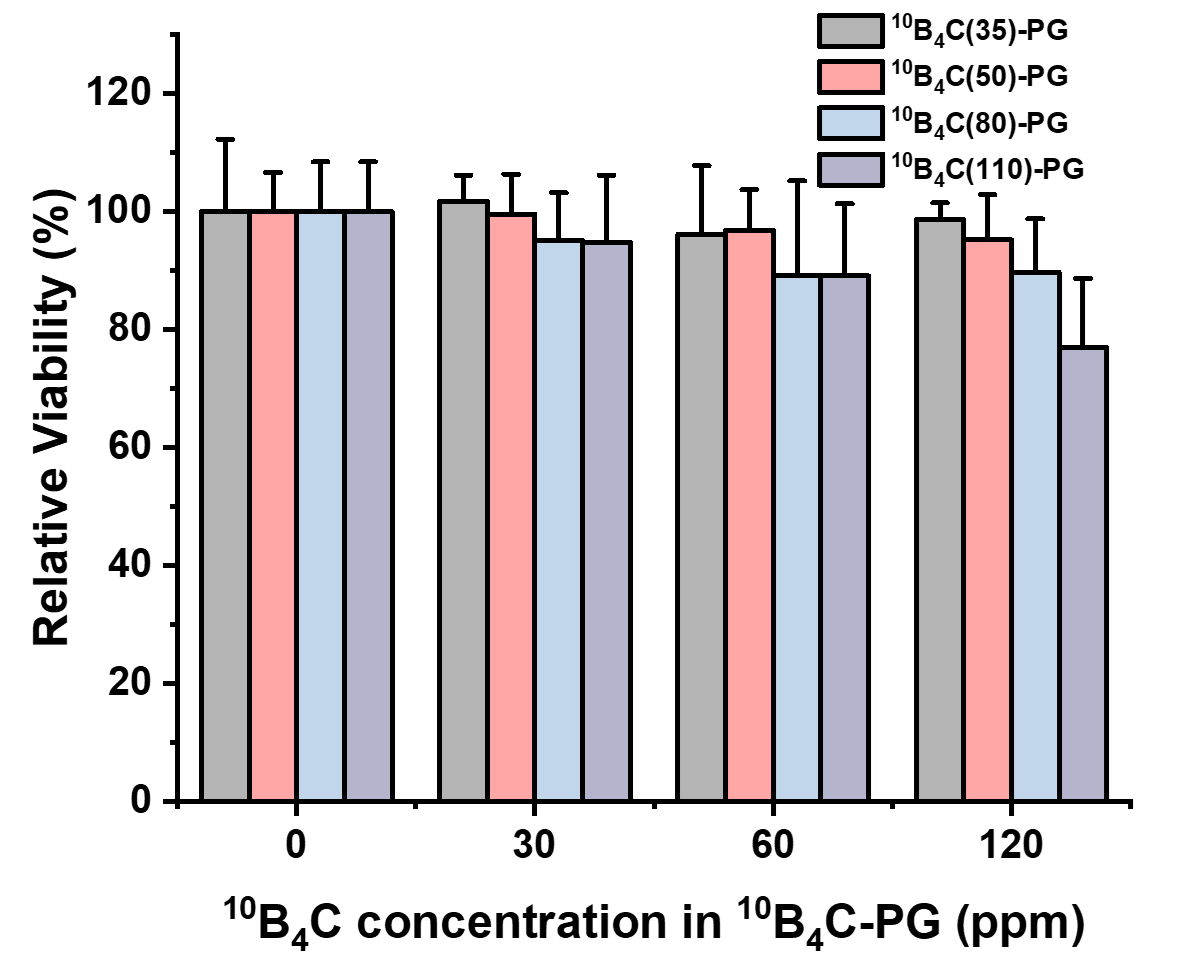

**Supplementary Fig. 1****2 |** Cell viability of CT26 after 24 h incubation of ^10^B_4_C-PGs with different core sizes at different concentrations. The experiments were carried out three times in quadruplicate.

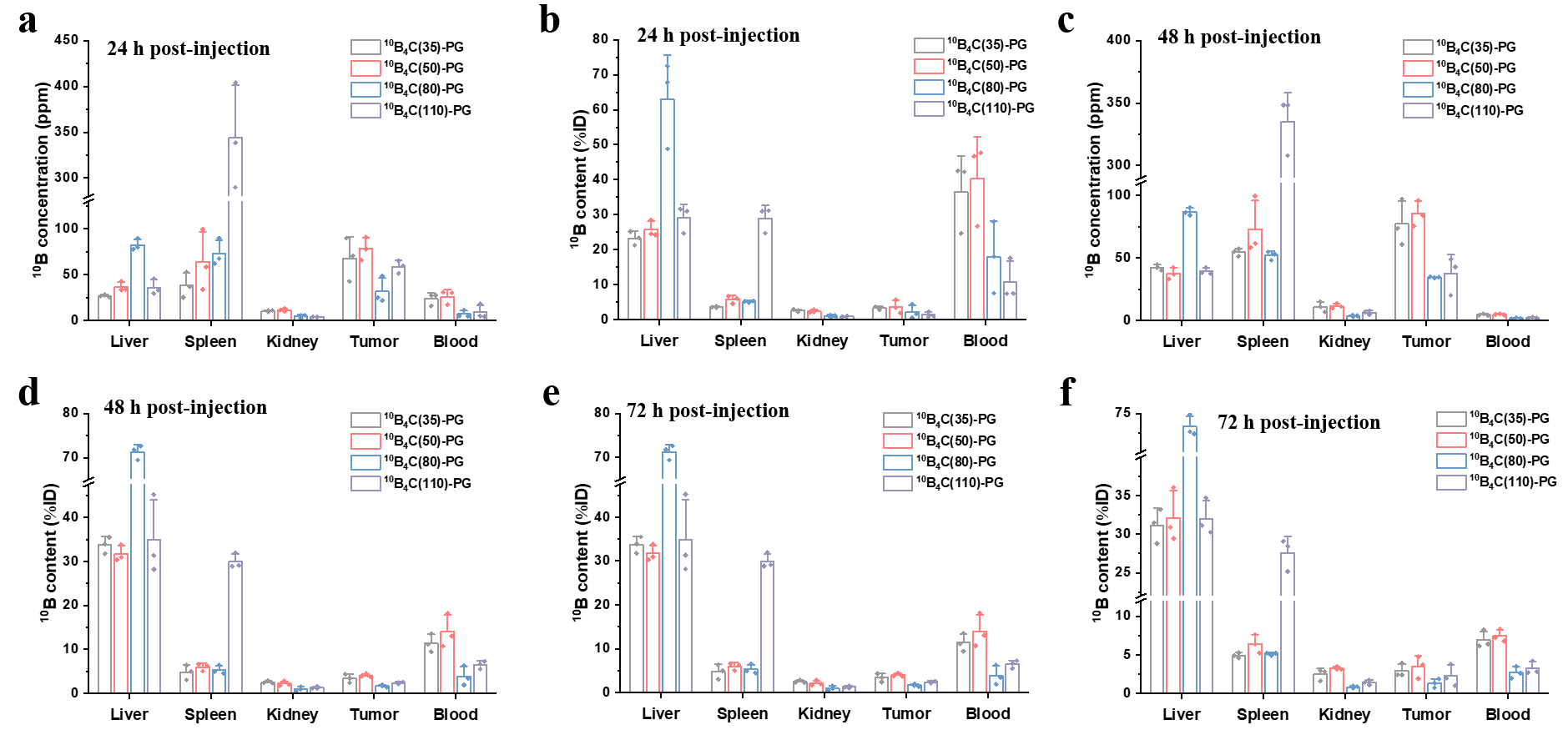

**Tumour**

**Tumour**

**Tumour**

**Tumour**

**Tumour**

**Tumour**

**Supplementary Fig. 13 |** Biodistribution of ^10^B_4_C-PGs with different sizes in liver, spleen, kidney, tumour and blood at a dosage of 5.1 mg [^10^B]/kg (mouse) *in vivo*. **a**, **b**, 24 h. **c, d**, 48 h. **e, f,** 72 h. Data are given as the mean ± SD (*n* = 3).

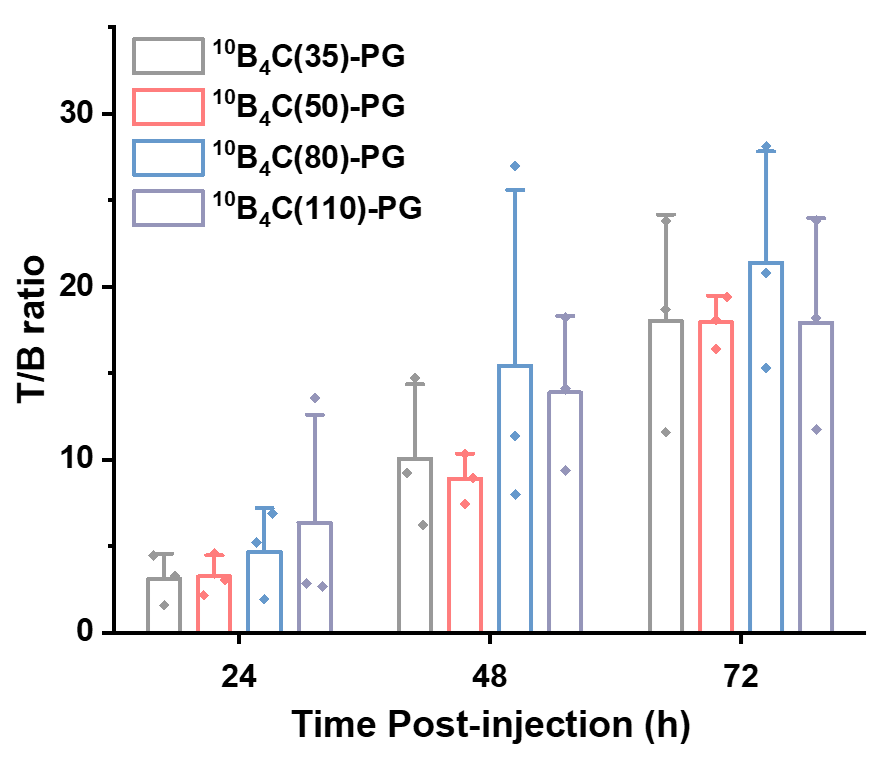

**Supplementary Fig. 14 |** The ^10^B concentration ratio between tumour and blood (T/B ratio) based on Supplementary Fig. 13. Data are given as the mean ± SD (*n* = 3).

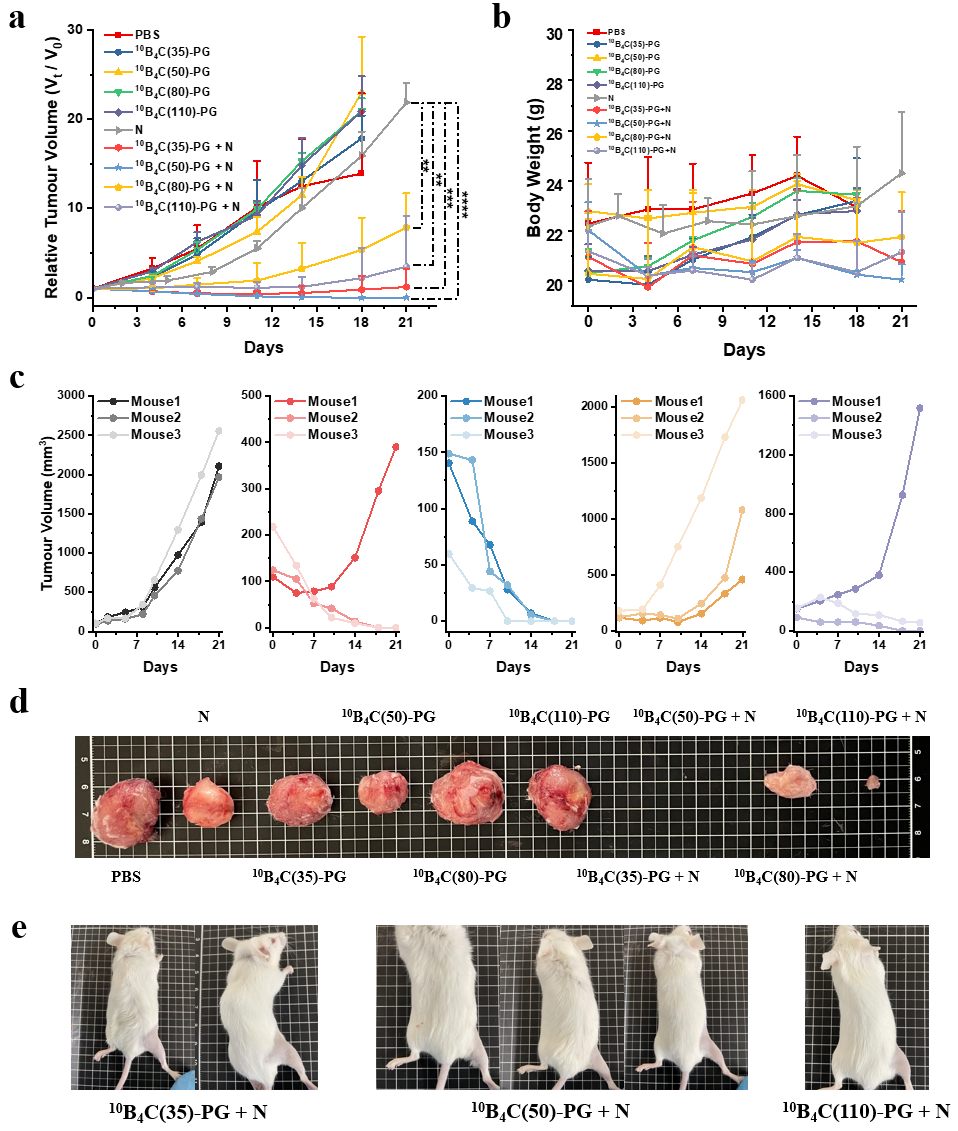

**Supplementary Fig. 15 | Therapeutic efficacy of ^10^B_4_C-PG with different sizes at dosage of 5.1 mg [^10^B]/kg (mouse) in *in vivo* BNCT. a**, Relative tumour volume monitored for 21 days. **b**, Time course of body weight for 21 days. **c**, Growth curves of individual in neutron control (N) and ^10^B_4_C(*Y*)-PG + N (*Y* = 35, 50, 80 and 110) groups (from left to right). **d**, Typical image of the explanted tumours in PBS, ^0^B_4_C(*Y*)-PG , N and ^10^B_4_C(*Y*)-PG + N (*Y* = 35, 50, 80 and 110) after 21 days (from left to right). **e**, Images of cured mice. Data are given as the mean ± SD (*n* = 3). Statistical significance: *p < 0.05, **p < 0.01, ***p < 0.001, ****p < 0.0001 and NS (no statistical difference).

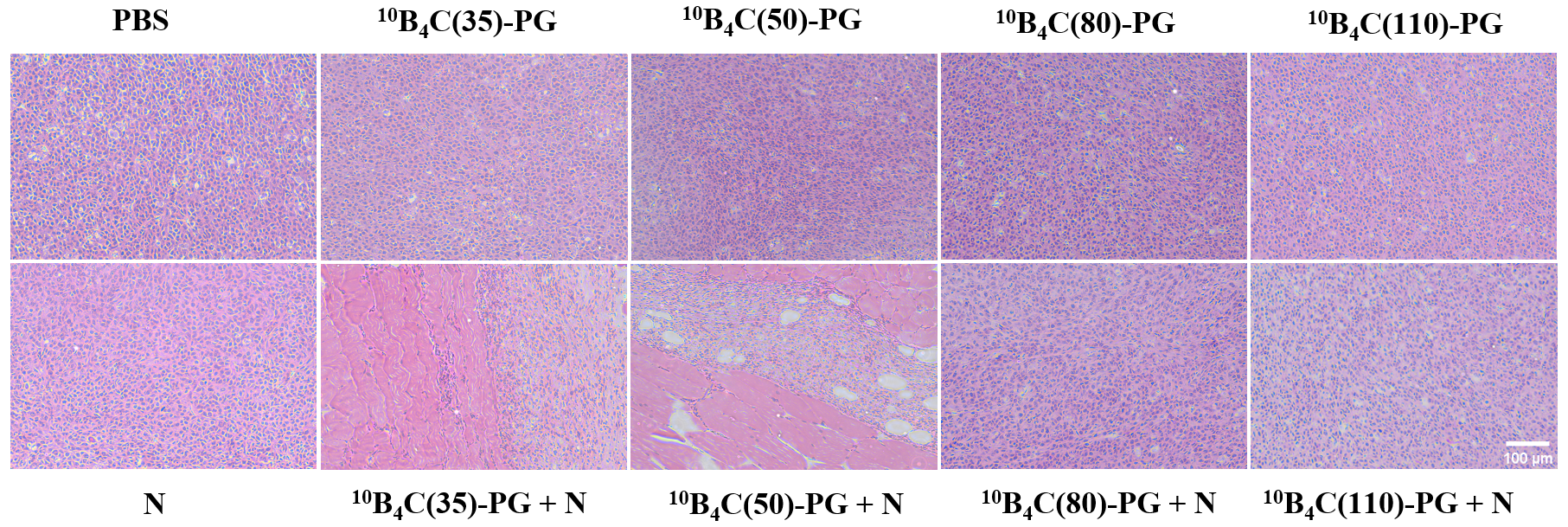

**Supplementary Fig. 16 |** H&E staining of tumour tissue in mouse from each group in Fig. 1f. Scale bar 100 µm.

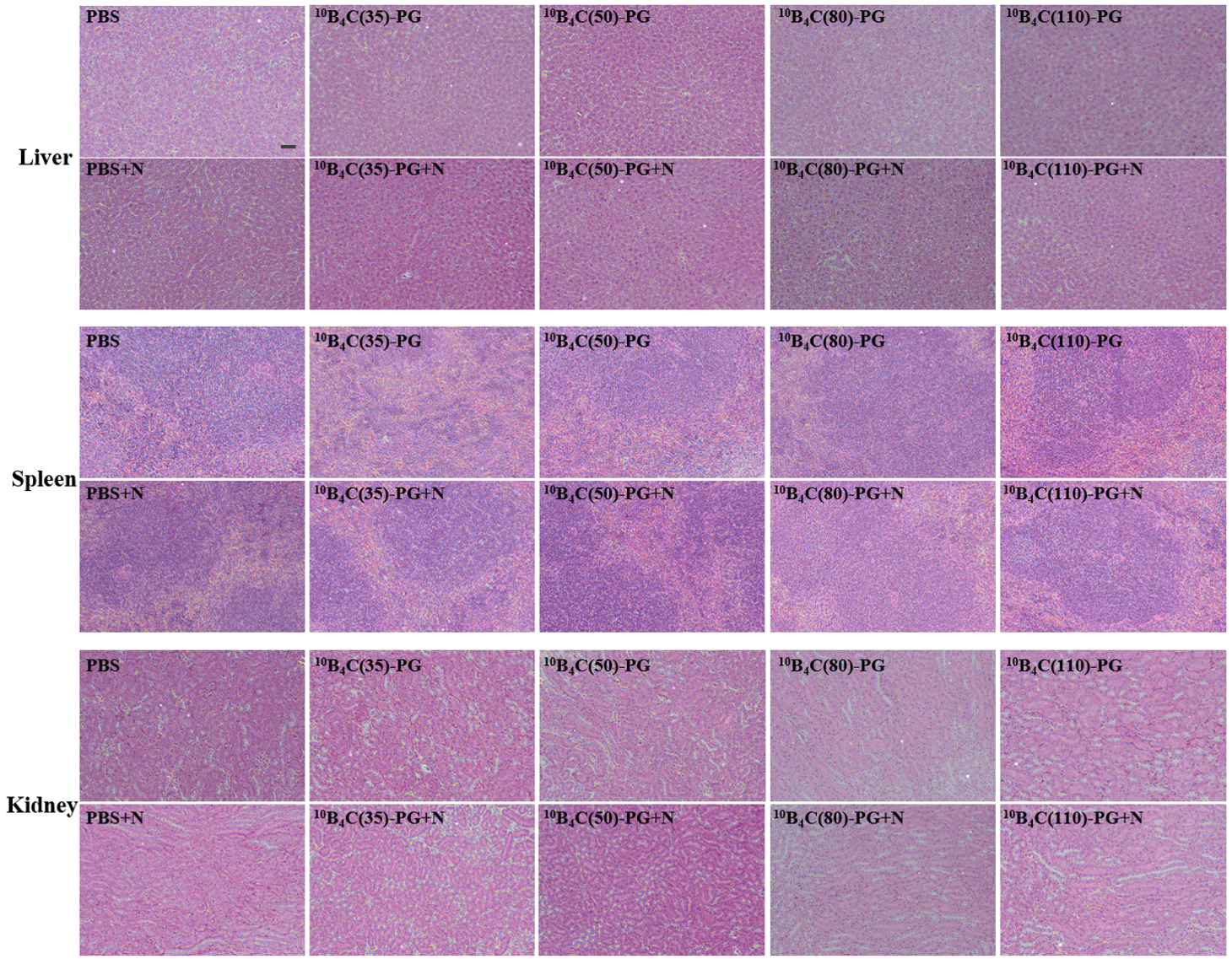

**Supplementary Fig. 17 |** H&E staining of liver, spleen and kidney in mouse from each group in Fig. 1f. Scale bar, 50 µm..

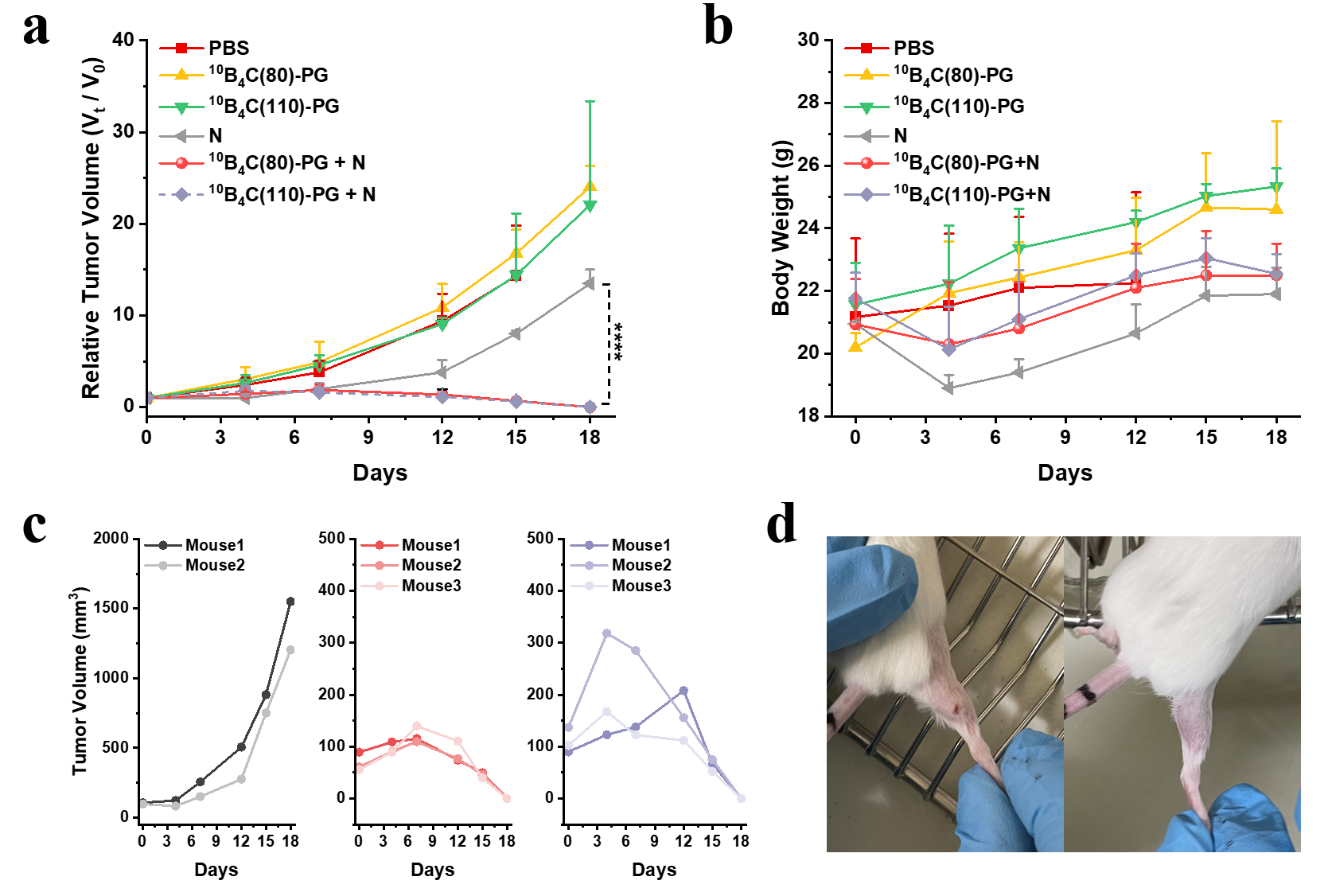

**Relative Tumour Volume (*V*_t_/*V*_0_)**

**Tumour Volume (mm^3^)**

**Supplementary Fig. 18 | CT26 tumour treatment with ^10^B_4_C(80)-PG and ^10^B_4_C(110)-PG at dosage of** **21.4 mg [^10^B]/kg (mouse) by *in vivo* BNCT.** **a**, Relative tumour volume monitored for 18 days. **b**, Time course of body weights for 18 days. **c**, Growth curves of individual tumour for N and ^10^B_4_C(*Y*)-PG + N groups (*Y* = 80 and 110) from left to right. **d**, Typical images of cured mice in ^10^B_4_C(80)-PG + N and ^10^B_4_C(110)-PG + N groups. Data are given as the mean ± SD (*n* = 2 in N group and *n* = 3 in other groups).

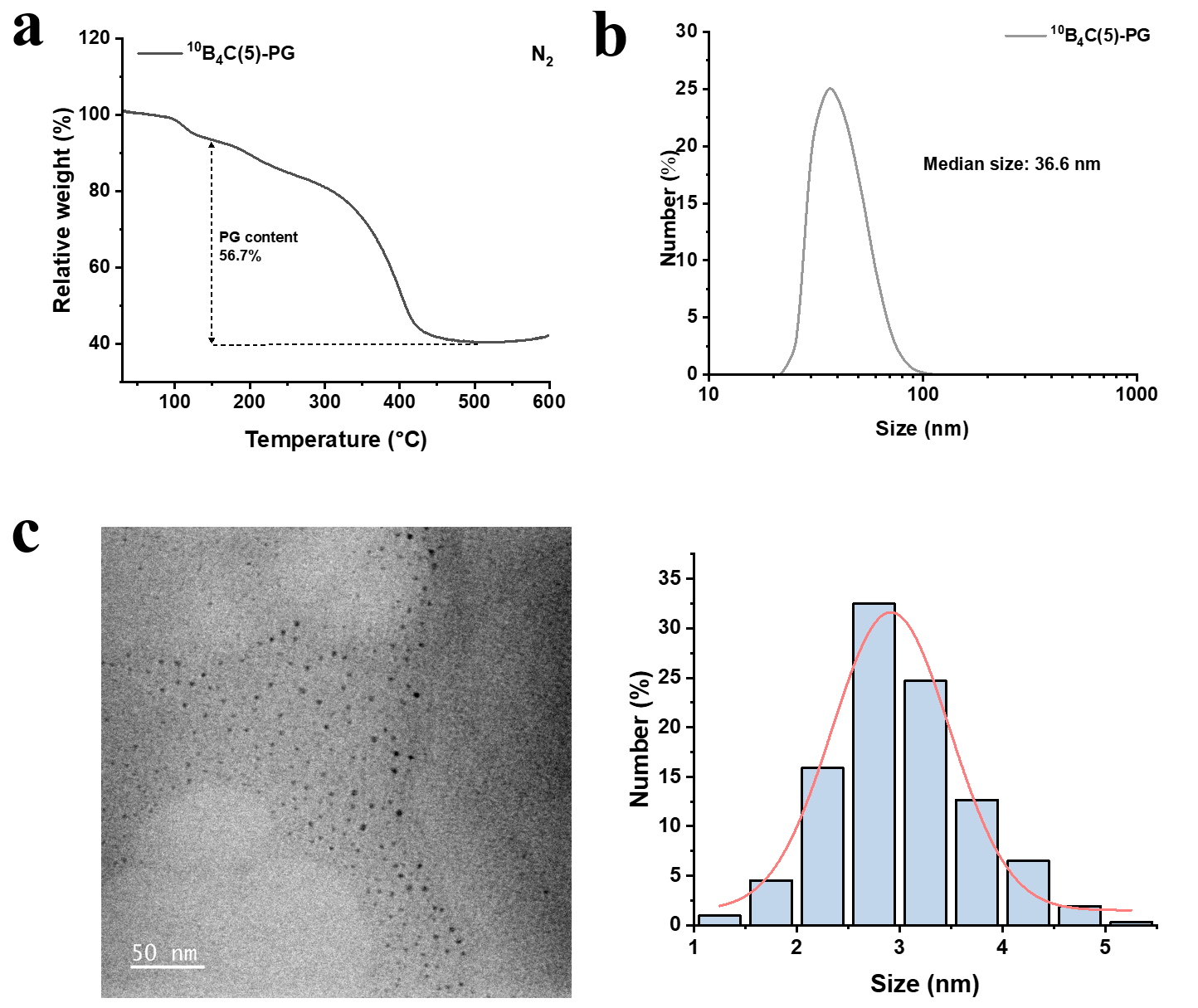

**Supplementary Fig. 19 | Characterization of ^10^B_4_C(5)-PG. a**, TGA curves of ^10^B_4_C(5)-PG in nitrogen atmosphere. **b**, Hydrodynamic size of ^10^B_4_C(5)-PG dispersed in PBS (*n* = 5). **c**, Core size distribution of ^10^B_4_C(5)-PG observed in TEM (*n* = 308).

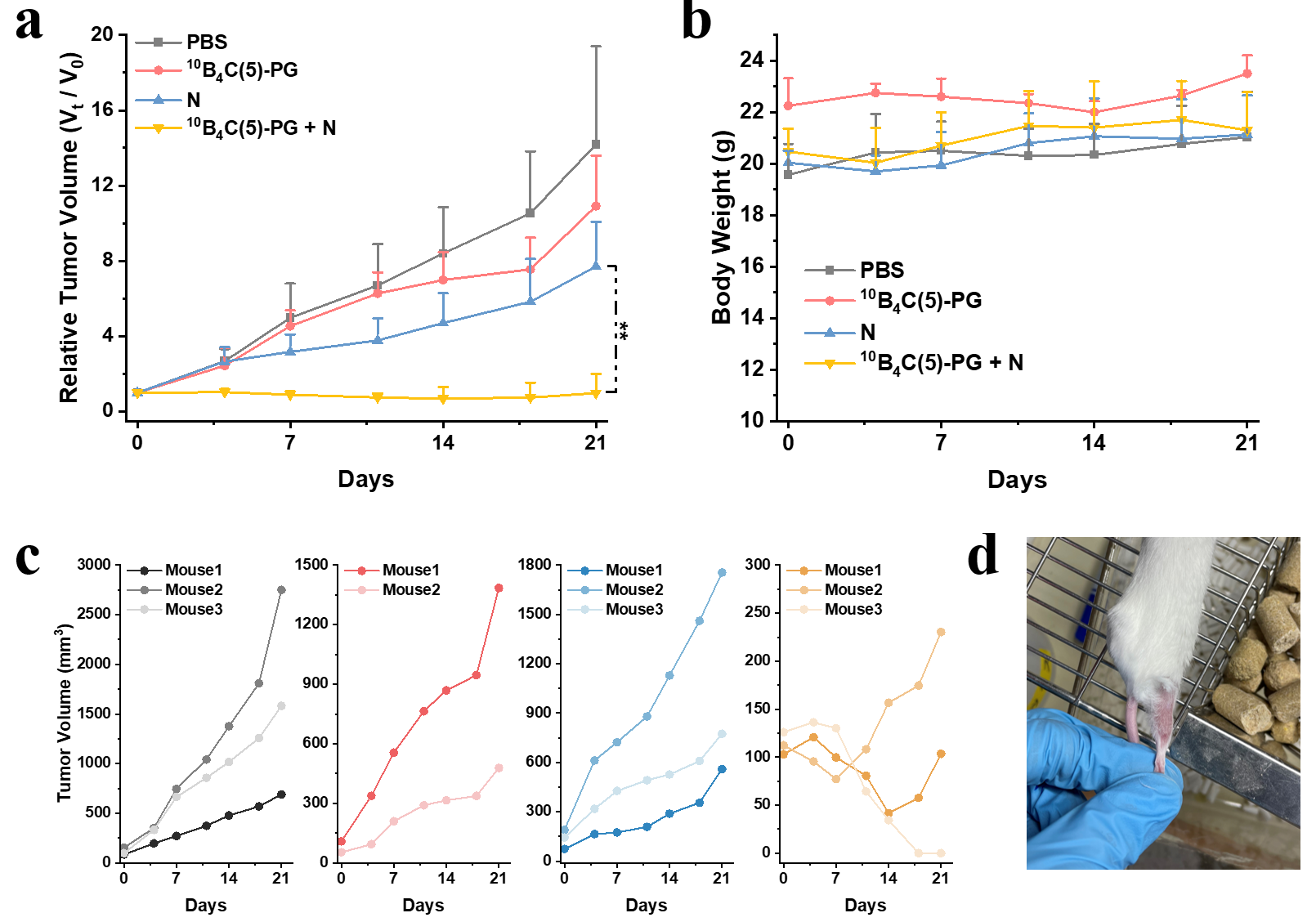

**Tumour Volume (mm^3^)**

**Relative Tumour Volume (*V*_t_/*V*_0_)**

**Supplementary Fig. 20 | CT26 tumour treatment with^10^B_4_C(5)-PG at dosage of 5.2 mg [^10^B]/kg (mouse) by *in vivo* BNCT.** **a**, Relative tumour volume monitored for 21 days. **b**, Time course of body weights for 21 days. **c**, Growth curves of individual tumour in PBS, ^10^B_4_C(5)-PG, N and ^10^B_4_C(5)-PG + N groups. **d**, Image of cured mouse in ^10^B_4_C(5)-PG + N group after 187 days post irradiation. Data are given as the mean ± SD (*n* = 2 in ^10^B_4_C(5)-PG group and *n* = 3 in other groups).

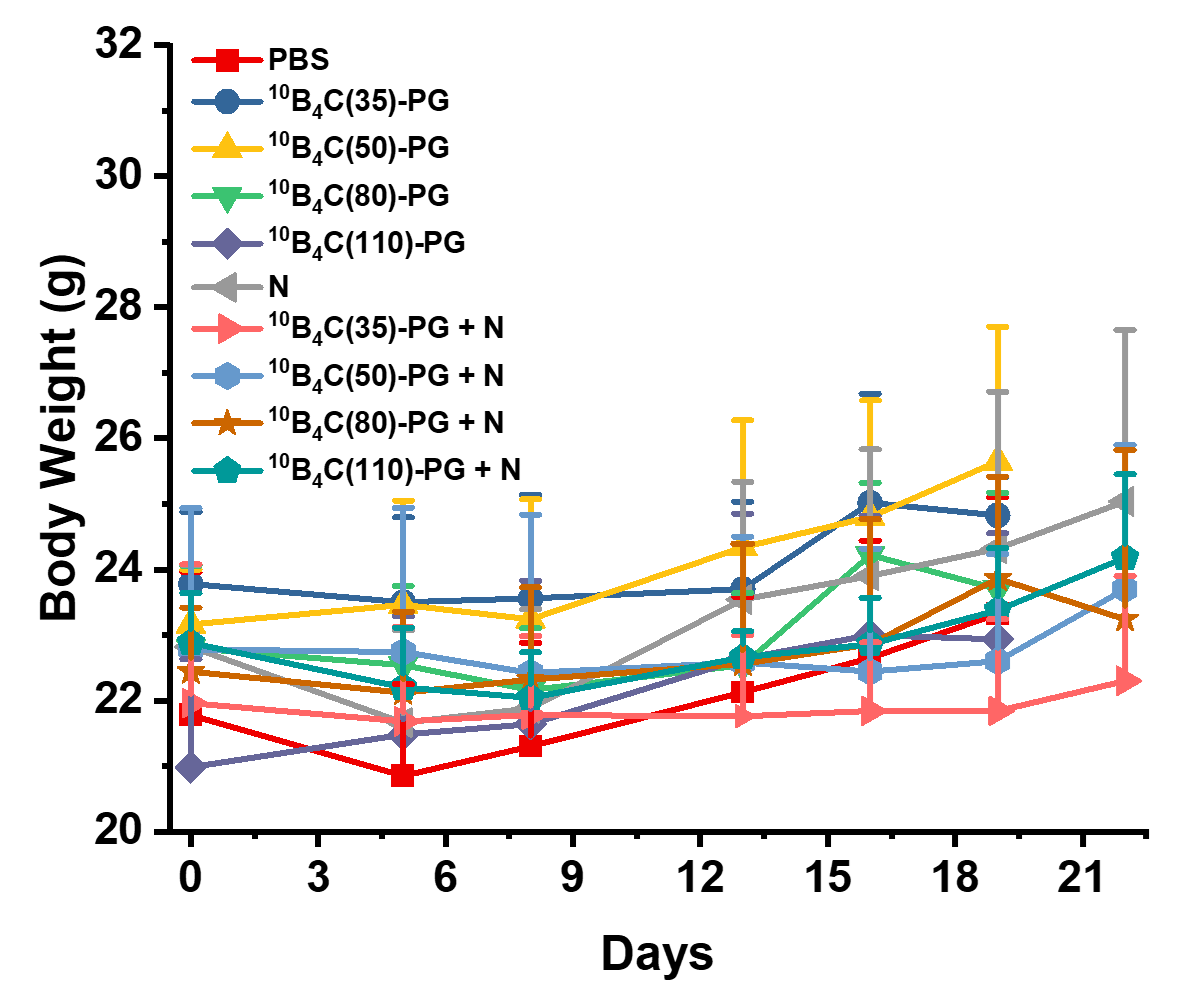

**Supplementary Fig. 21 |** Time course of body weights of mice in Fig. 1f.

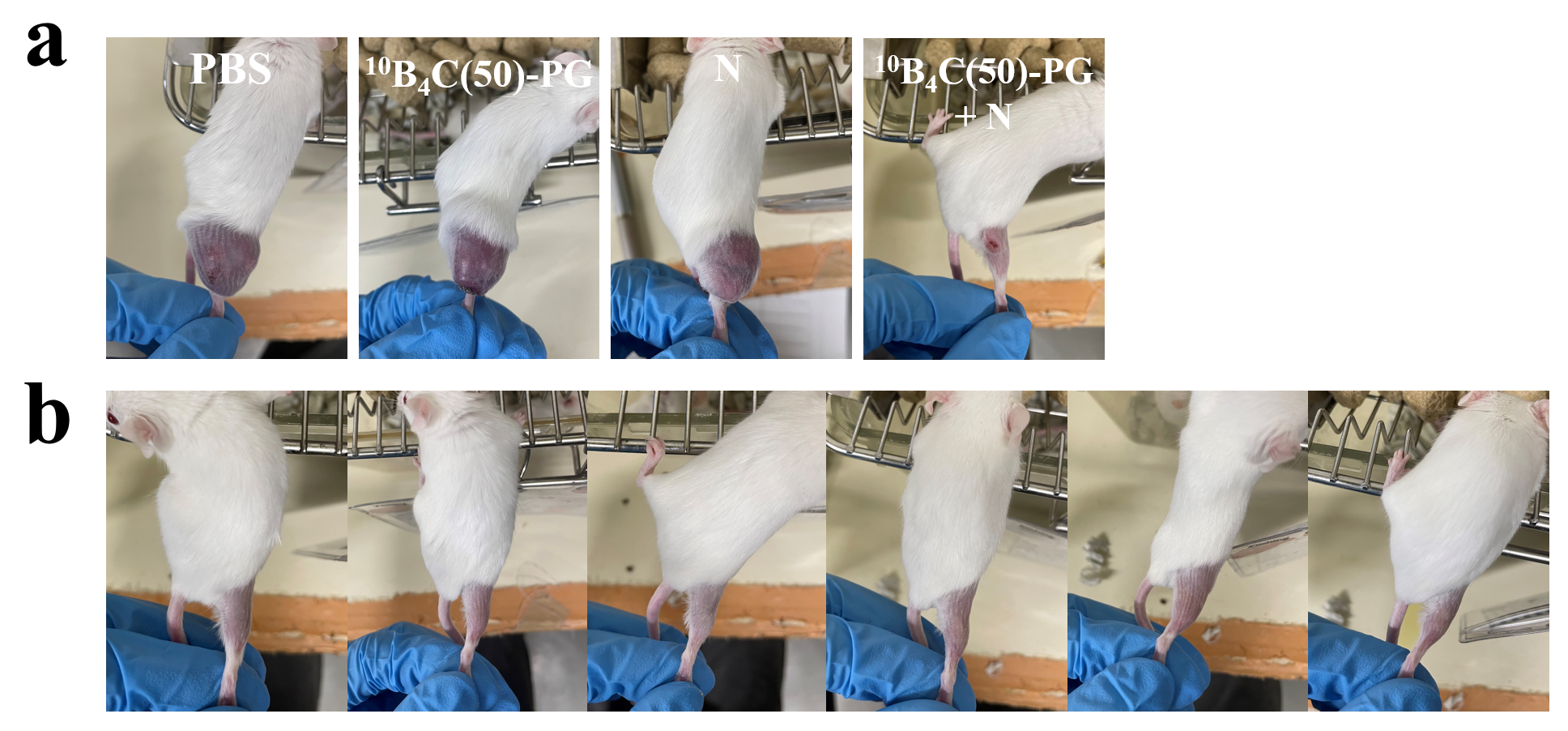

**Supplementary Fig. 22 | a**, Images of typical mice from four groups with average initial volumes of 171 mm^3^ after 18 days post neutron irradiation as shown in Fig. 2b. **b**, Images of cured mice at ^10^B_4_C(50)-PG + N group after 18 days post neutron irradiation as shown in Fig. 2b.

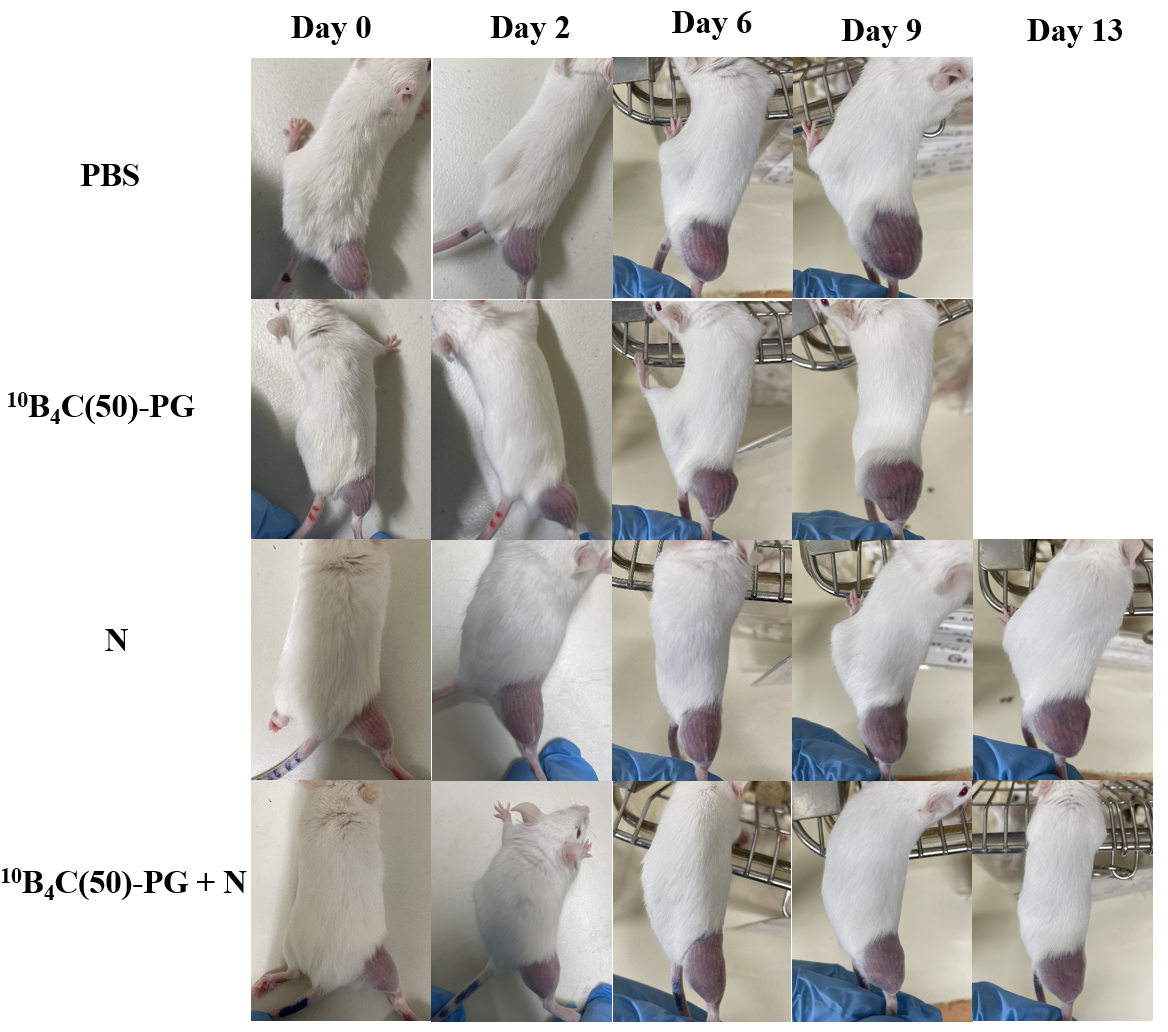

**Supplementary Fig. 23 |** Images of typical mice from four groups with average initial volumes of 645 mm^3^ after 0, 2, 6, 9 and 13 days post neutron irradiation as shown in Fig. 2c.

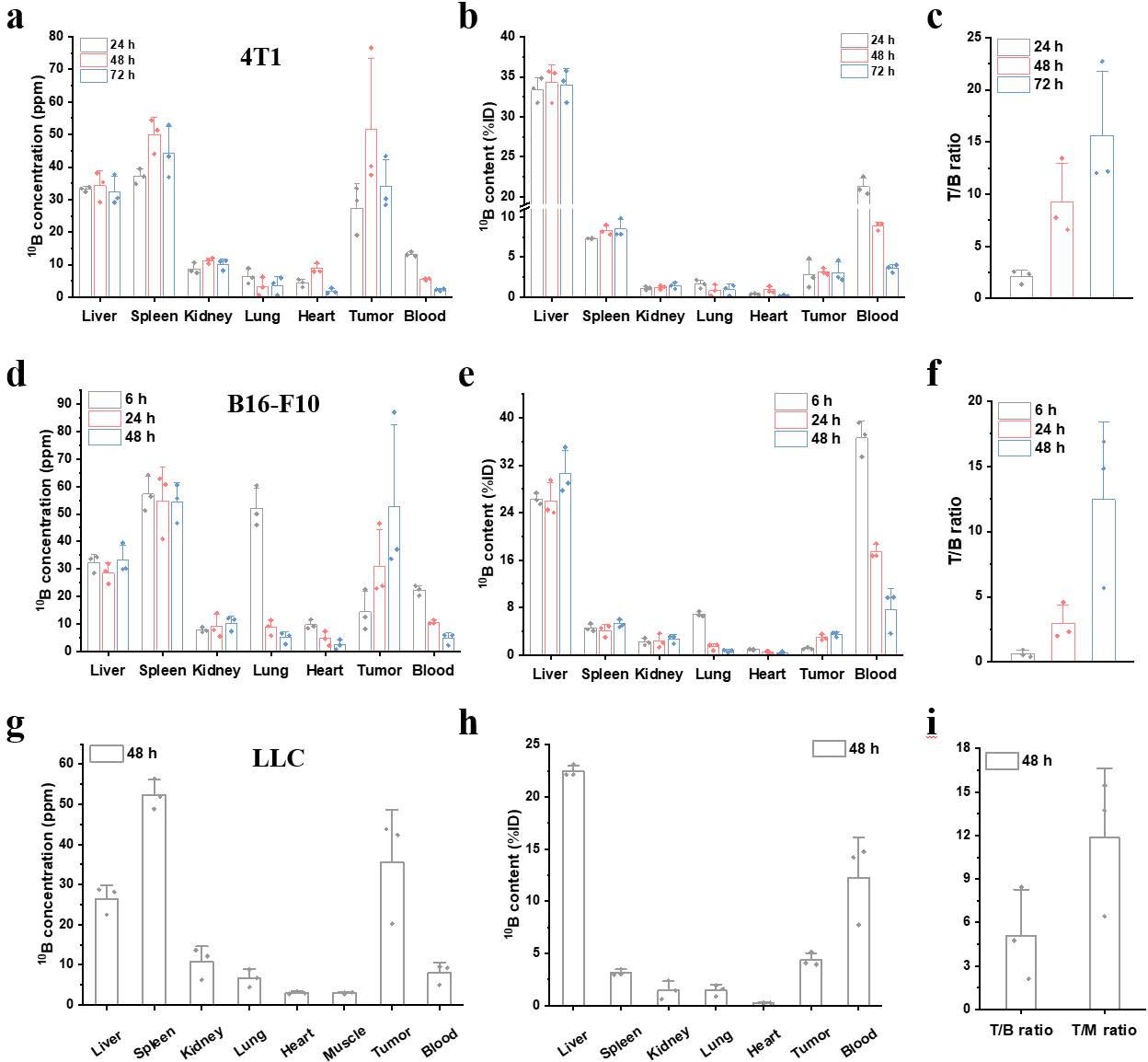

**Tumour**

**Tumour**

**Supplementary Fig. 24 |** Biodistribution of ^10^B_4_C(50)-PG in major organs, muscle, tumour and blood, and ^10^B concentration ratios between tumour and blood (T/B) and tumour and muscle (T/M) at different timepoints after injection at a dosage of 5.2 mg [^10^B]/kg (mouse) in 4T1 (**a** - **c**), and 5.1 mg [^10^B]/kg (mouse) in B16-F10 (**d** - **f**) and LLC (**g** - **i**) model mice. Data are given as the mean ± SD (*n* = 3).

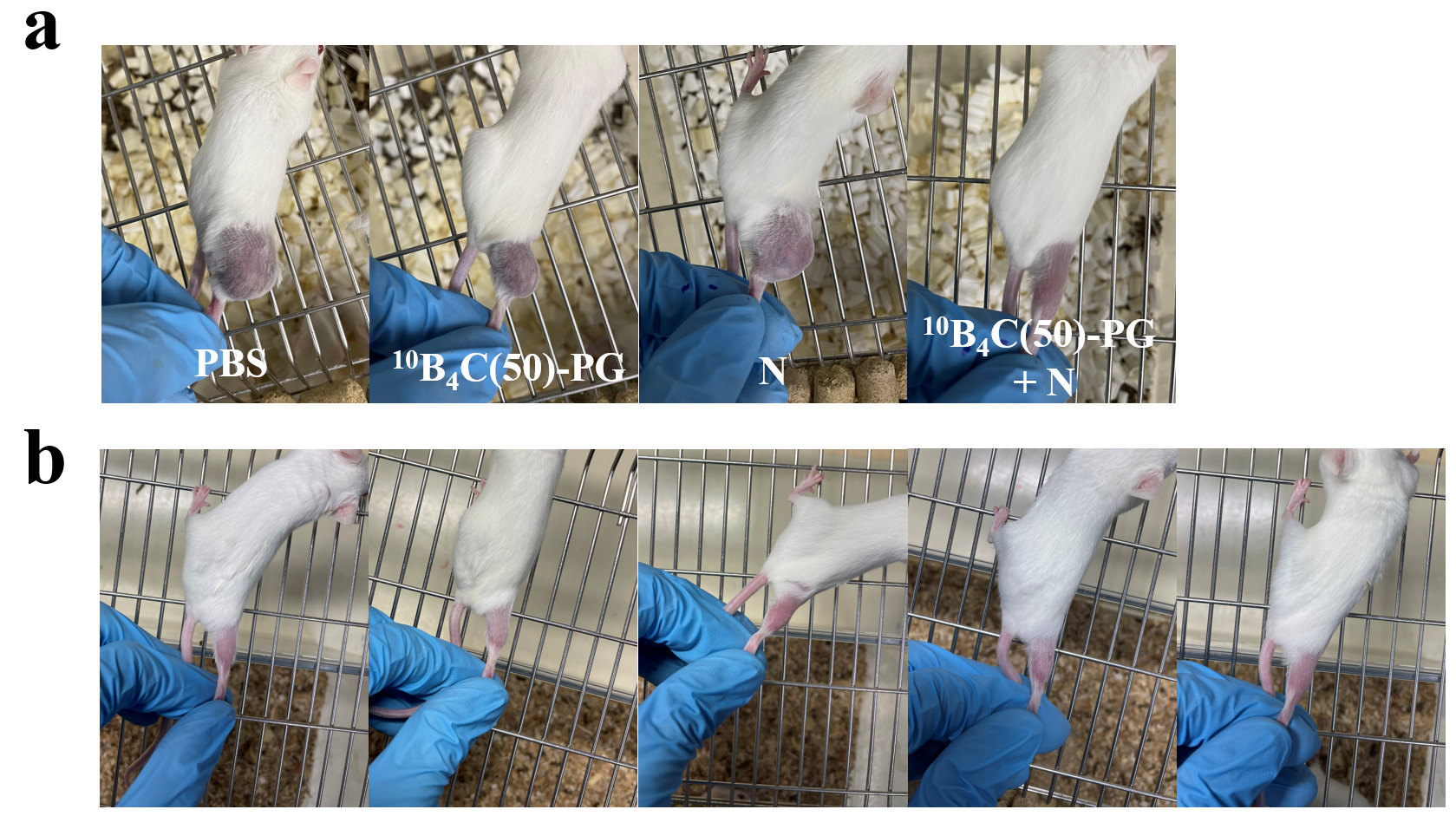

**Supplementary Fig. 25 | a**, Typical tumour image after 18 days post-irradiation at different groups in Meth-A tumour models (Fig. 2e). **b**, Image of cured mice after 163 days post-injection (Fig. 2f).

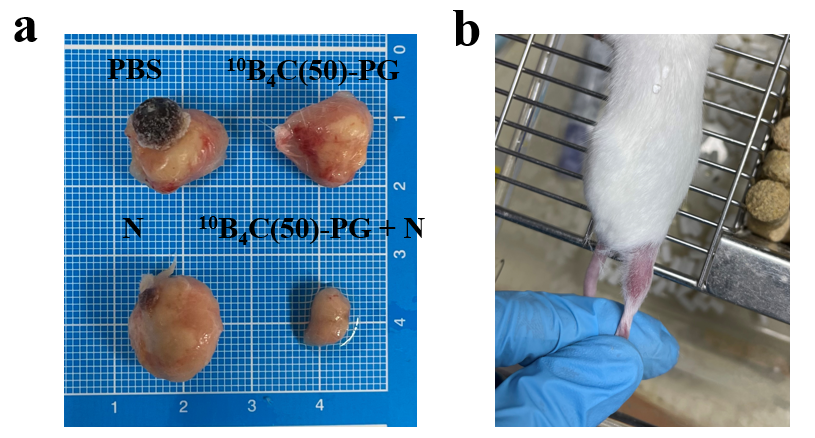

**Supplementary Fig. 26 | a**, Typical tumour image after 33 days post-injection at different groups in 4T1 tumour models (Fig. 2g). **b**, Image of cured mouse after 250 days post-injection.

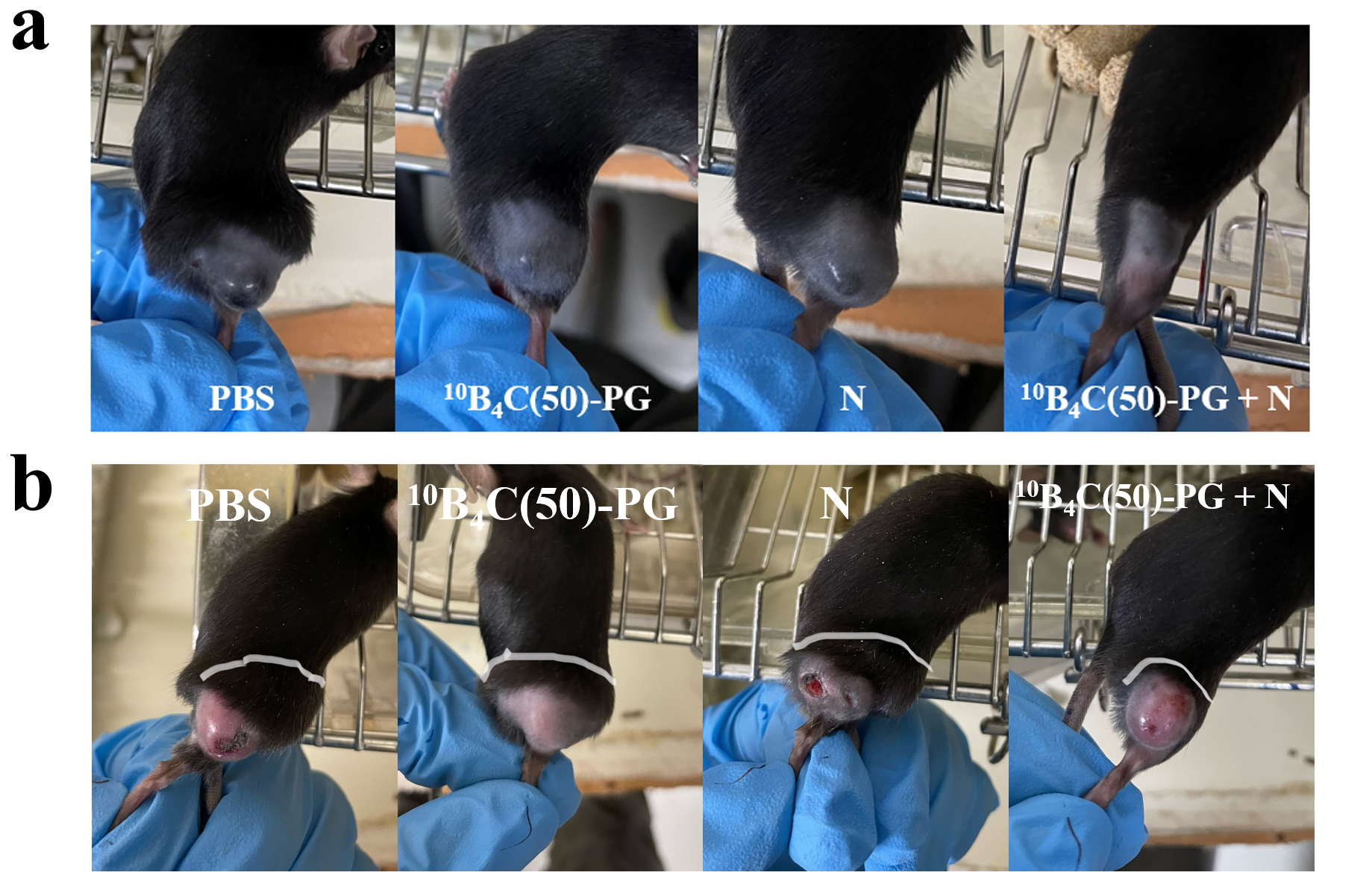

**Supplementary Fig. 27 | a**, Typical tumour images after 10 days post-irradiation at different groups in B16-F10 tumour models (Fig. 2h). **b**, Images of LLC tumours at the day 11 in different groups in Fig. 2i.

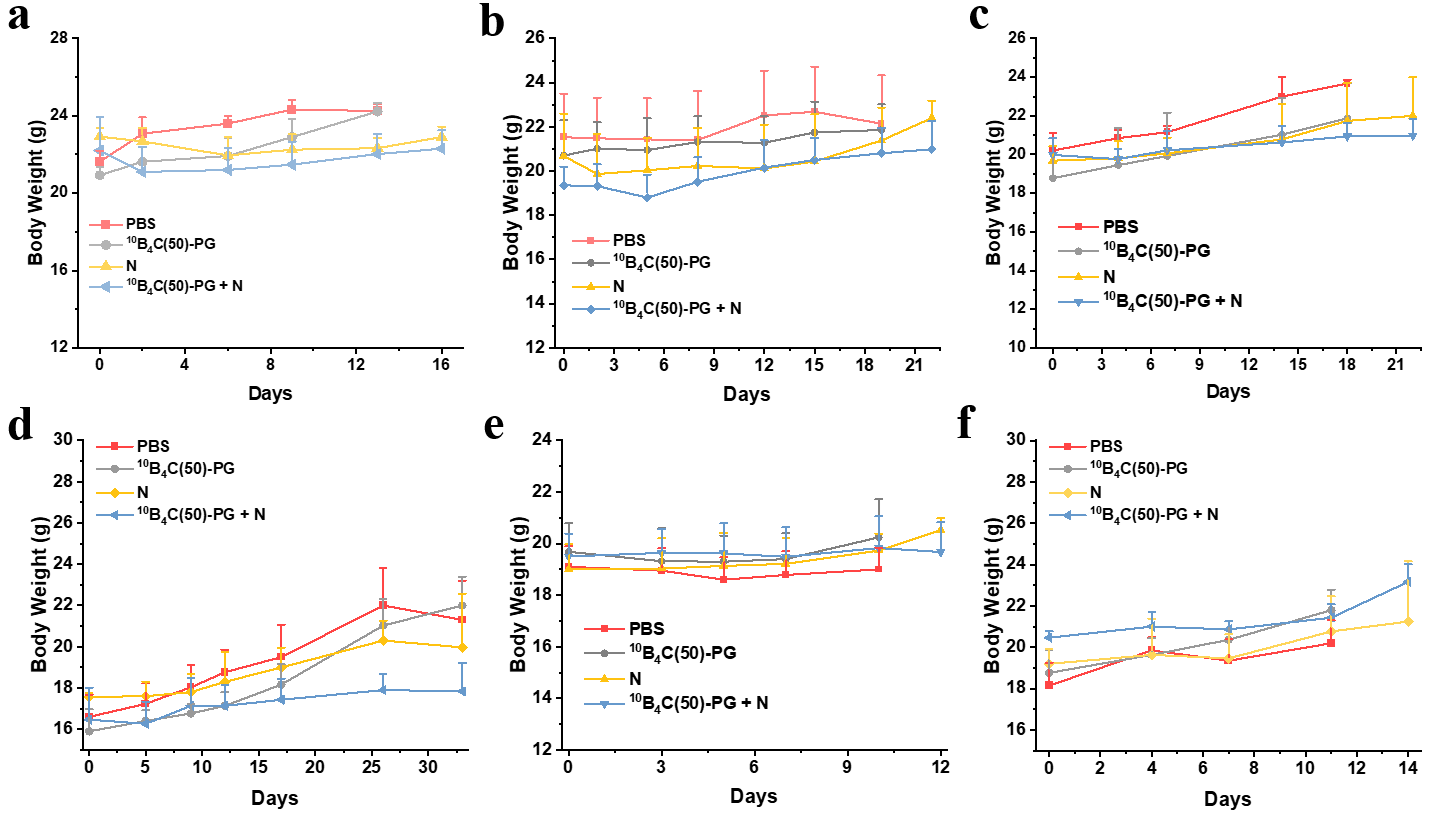

**Supplementary Fig. 28 | a,** Time course of body weights of mice in CT26 at 171 mm^3^ in Fig. 2b. **b,** CT26 at 645 mm^3^ in Fig. 2c. **c,** Meth-A in Fig. 2e. **d,** 4T1 in Fig. 2g. **e,** B16-F10 in Fig. 2h**. f,** LLC in Fig. 2i.

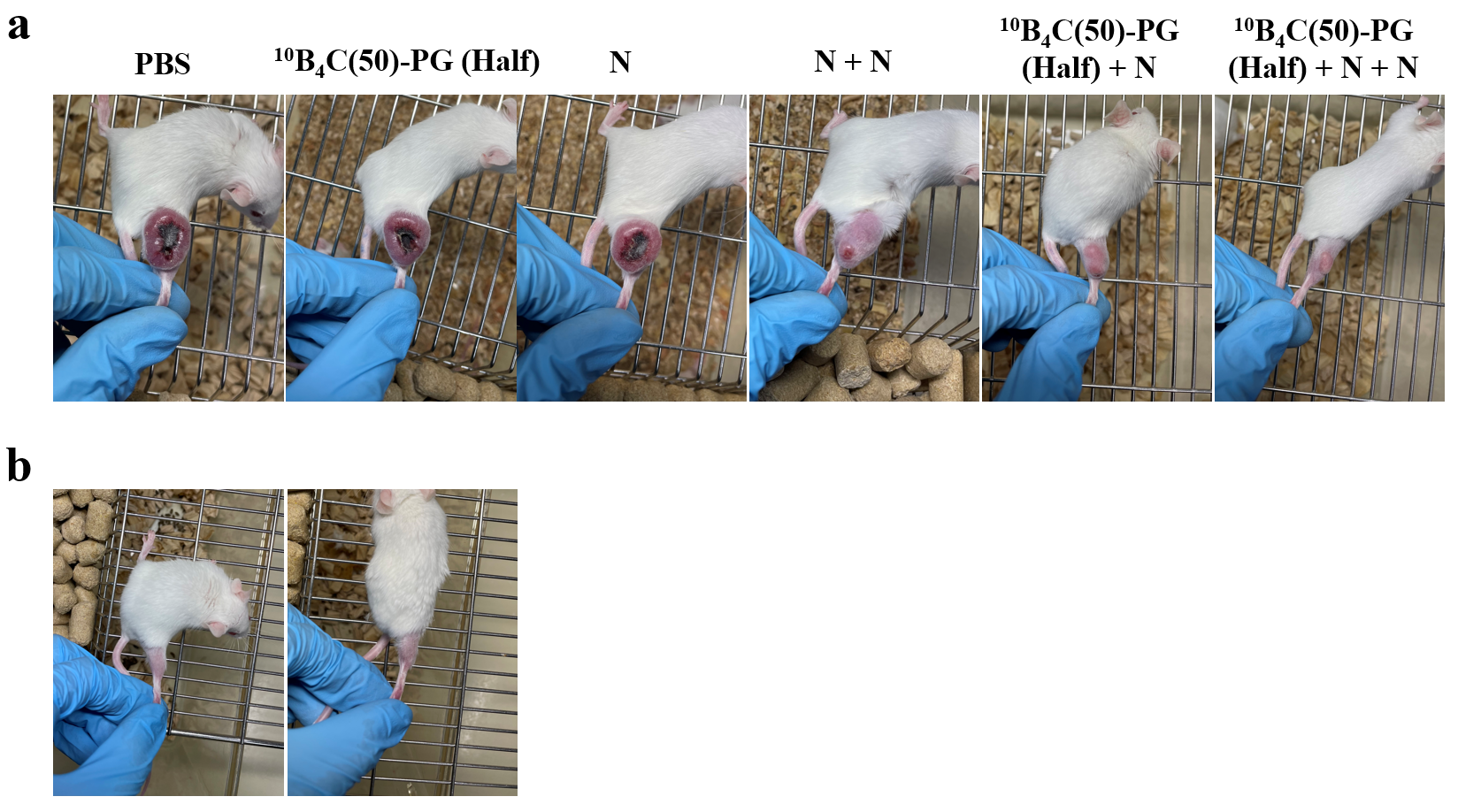

**Supplementary Fig. 29 | a**, Typical tumour images of mice at six groups in Fig. 2l after 20 days post irradiation. **b**, Tumour images of cured mice after 63 days post irradiation.

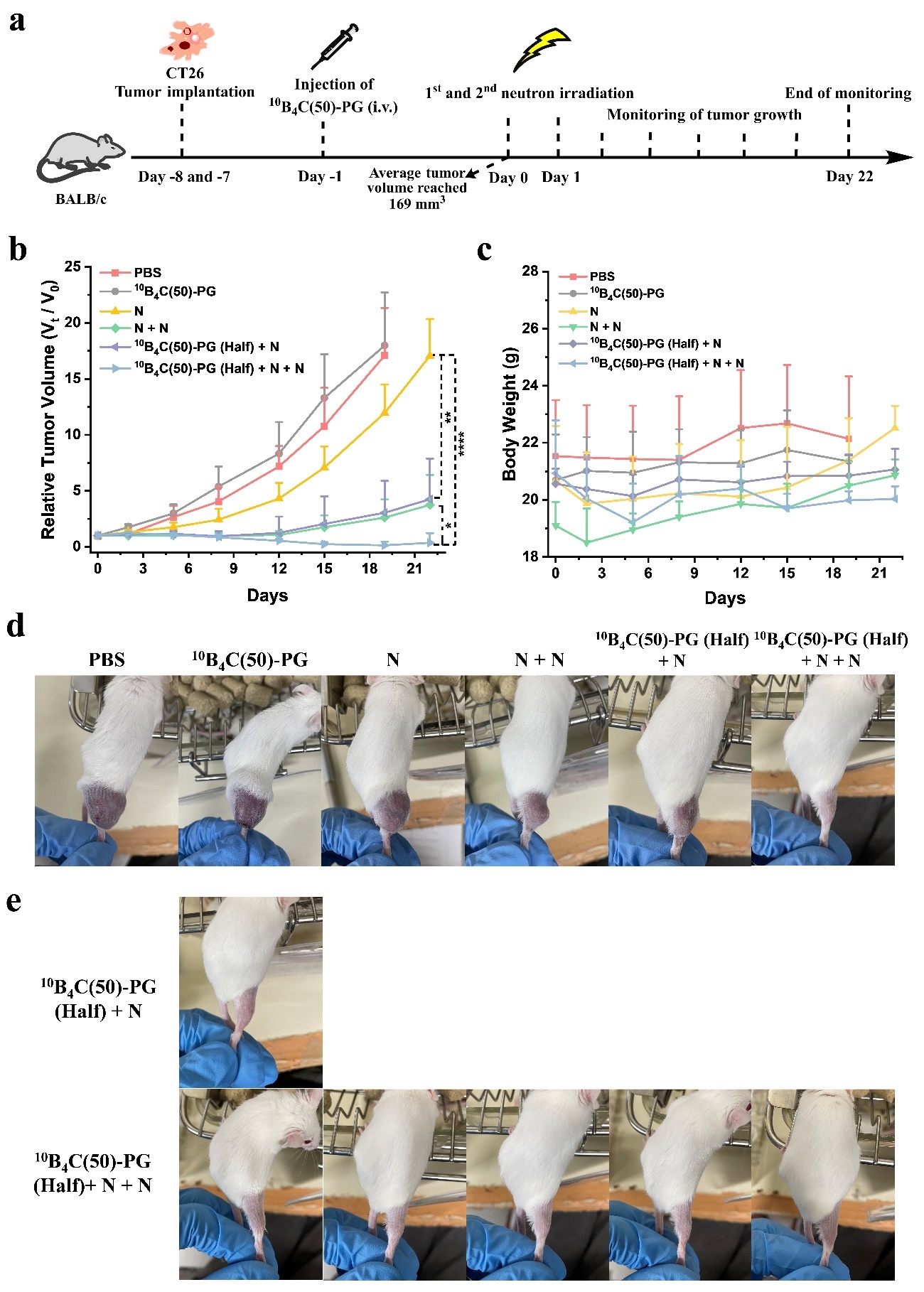

**Monitoring tumour growth**

**Relative Tumour Volume (*V*_t_/*V*_0_)**

**CT26 Tumour implantation**

**Supplementary Fig. 30 | CT26 tumour treatment process by BNCT at a dosage of 2.5 mg [^10^B_4_C(50)-PG]/kg (mouse) in double neutron irradiation.** **a**, Schematic of CT26 tumour treatment process. **b**, Relative tumour volume monitored for 21 days post-irradiation (*n* = 5 in N + N group and *n* = 6 in others). **c**, Time course of body weights of mice for 21 days. **d**, Tumour image of mouse in each group after 19 days post-irradiation. **e**, Images of cured mice. Data are given as the mean ± SD. The mice in PBS and N groups were shared from Fig. 2b since these experiments were performed at the same time.

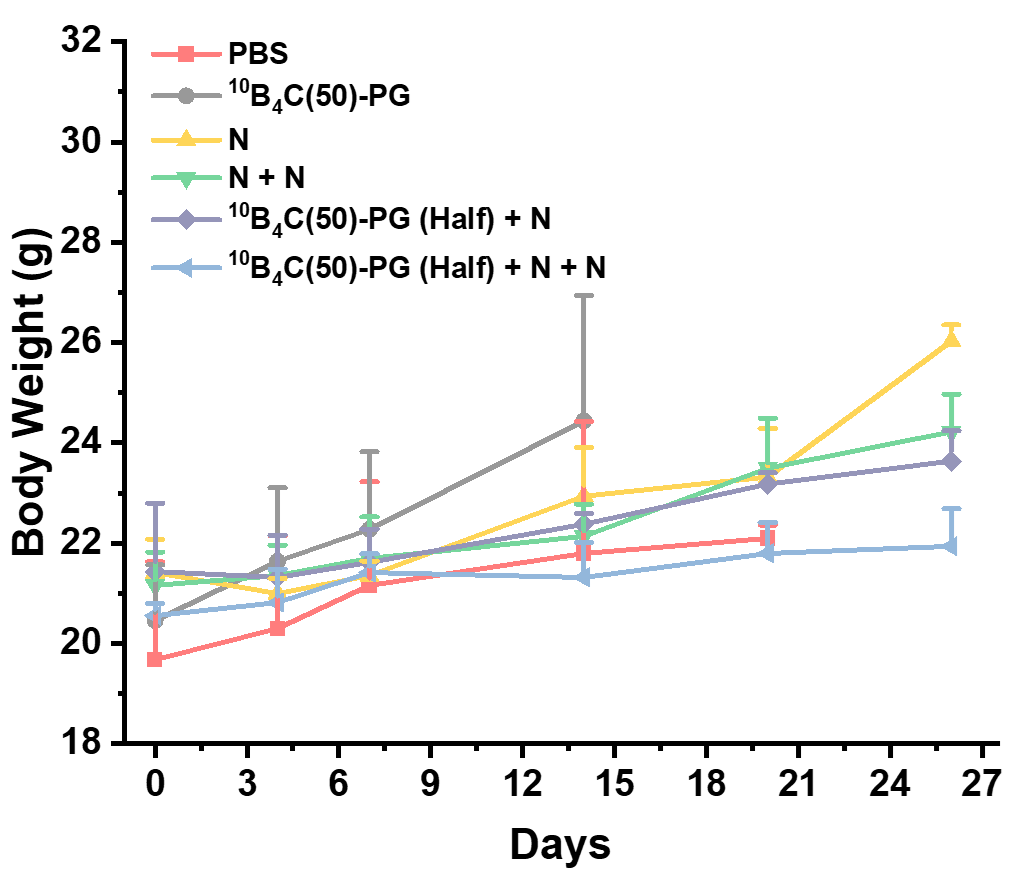

**Supplementary Fig. 31 |** Time course of body weights of mice in Fig. 2l.

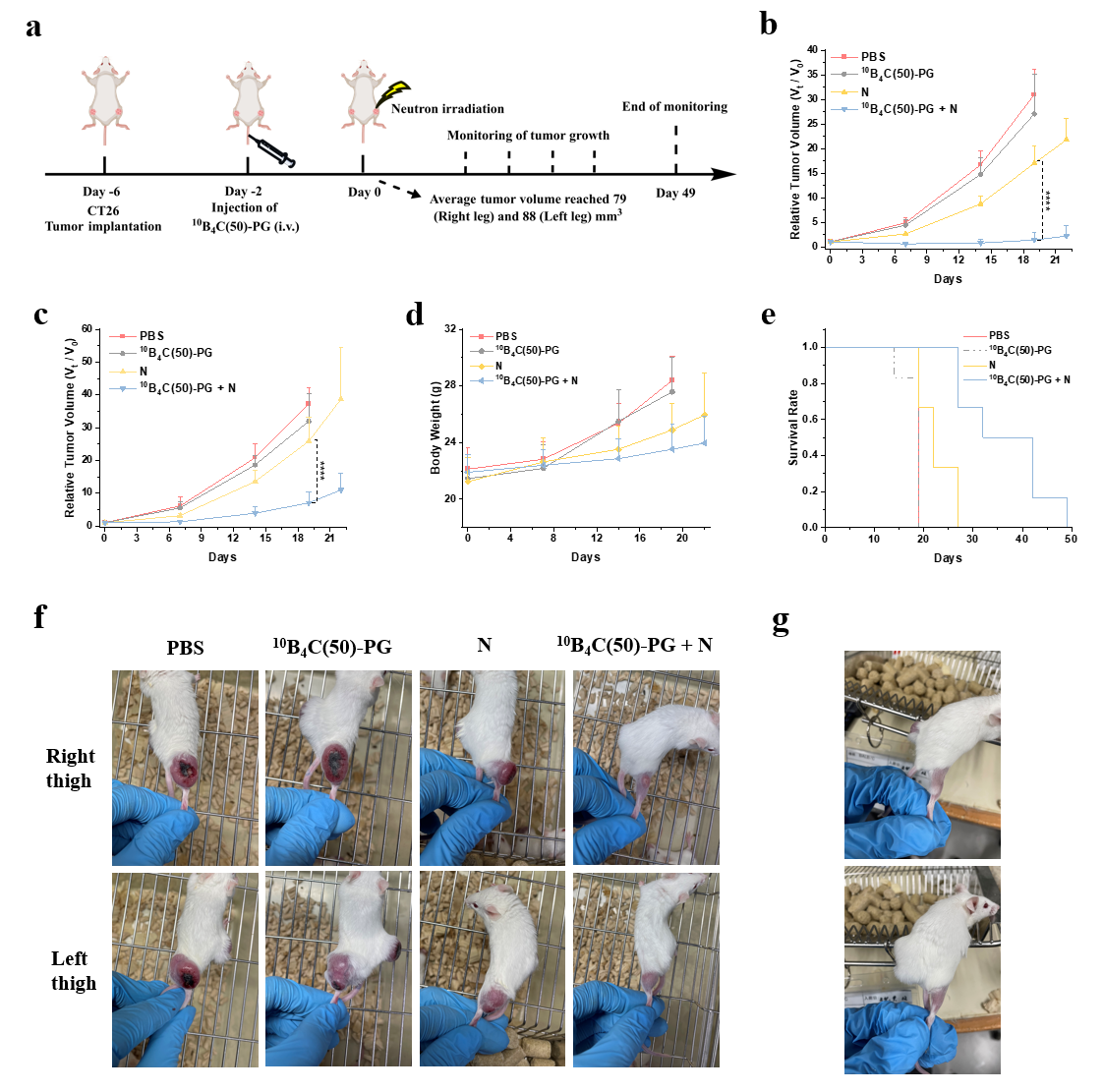

**Relative Tumour Volume (*V*_t_/*V*_0_)**

**Relative Tumour Volume (*V*_t_/*V*_0_)**

**Monitoring tumour growth**

**CT26 Tumour implantation**

**Supplementary Fig. 32 | Dual CT26 tumour treatment by *in vivo* BNCT at a dosage of 5.1 mg [^10^B]/kg (mouse).** **a**, Schematic of dual CT26 tumour treatment process with neutron irradiation only at right thigh in *in vivo* BNCT (*n* = 7) **b**, Relative primary tumour volumes in right leg, with direct neutron irradiation in N and ^10^B_4_C(50)-PG + N groups, monitored for 22 days. **c**, Relative distant tumour volumes in left leg, without direct neutron irradiation in N and ^10^B_4_C(50)-PG + N groups, monitored for 22 days. **d**, Time course of body weights for 22 days. **e**, Survival rates of mice in different groups of dual tumour models (*n* = 6). **e**, Tumour images at right and left legs of mice in each group after 22 days post irradiation. **f**, Images of mice with tumour eradicated at right leg after 27 days post irradiation. Statistical analysis of relative tumour volume with one-way ANOVA post Bonferroni test; *p < 0.05, **p < 0.01, ***p < 0.001, ****p < 0.0001.

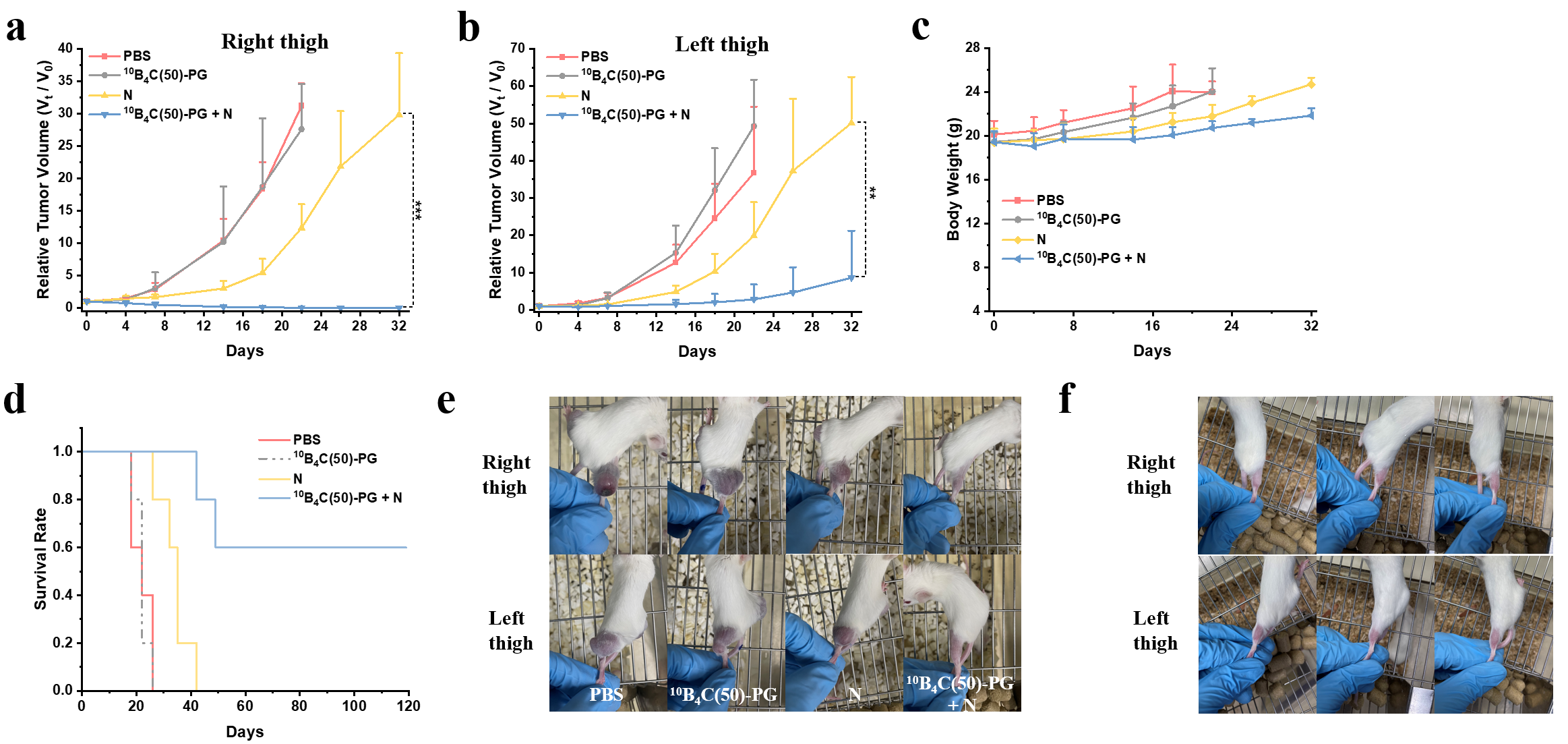

**Relative Tumour Volume (*V*_t_/*V*_0_)**

**Relative Tumour Volume (*V*_t_/*V*_0_)**

**Supplementary Fig. 33 | Dual CT26 tumour treatment by *in vivo* BNCT at a dosage of 5.1 mg [^10^B]/kg (mouse).** **a**, Relative primary tumour volumes in right thigh (average initial volume: 58 mm^3^), with direct neutron irradiation in N and ^10^B_4_C(50)-PG + N groups, monitored for 32 days. **b**, Relative distant tumour volumes in left thigh (average initial volume: 46 mm^3^), without direct neutron irradiation in N and ^10^B_4_C(50)-PG + N groups, monitored for 32 days. **c**, Time course of body weights for 32 days. **d**, Survival rates of mice at different groups in dual tumour model. **e**, Tumour images at right and left thighs of mice in each group after 22 days post irradiation. **f**, Images at right and left legs of cured mice after 77 days post irradiation. Data are given as the mean ± SD (*n* = 5). Statistical analysis of relative tumour volume with one-way ANOVA post Bonferroni test; *p < 0.05, **p < 0.01, ***p < 0.001, ****p < 0.0001.

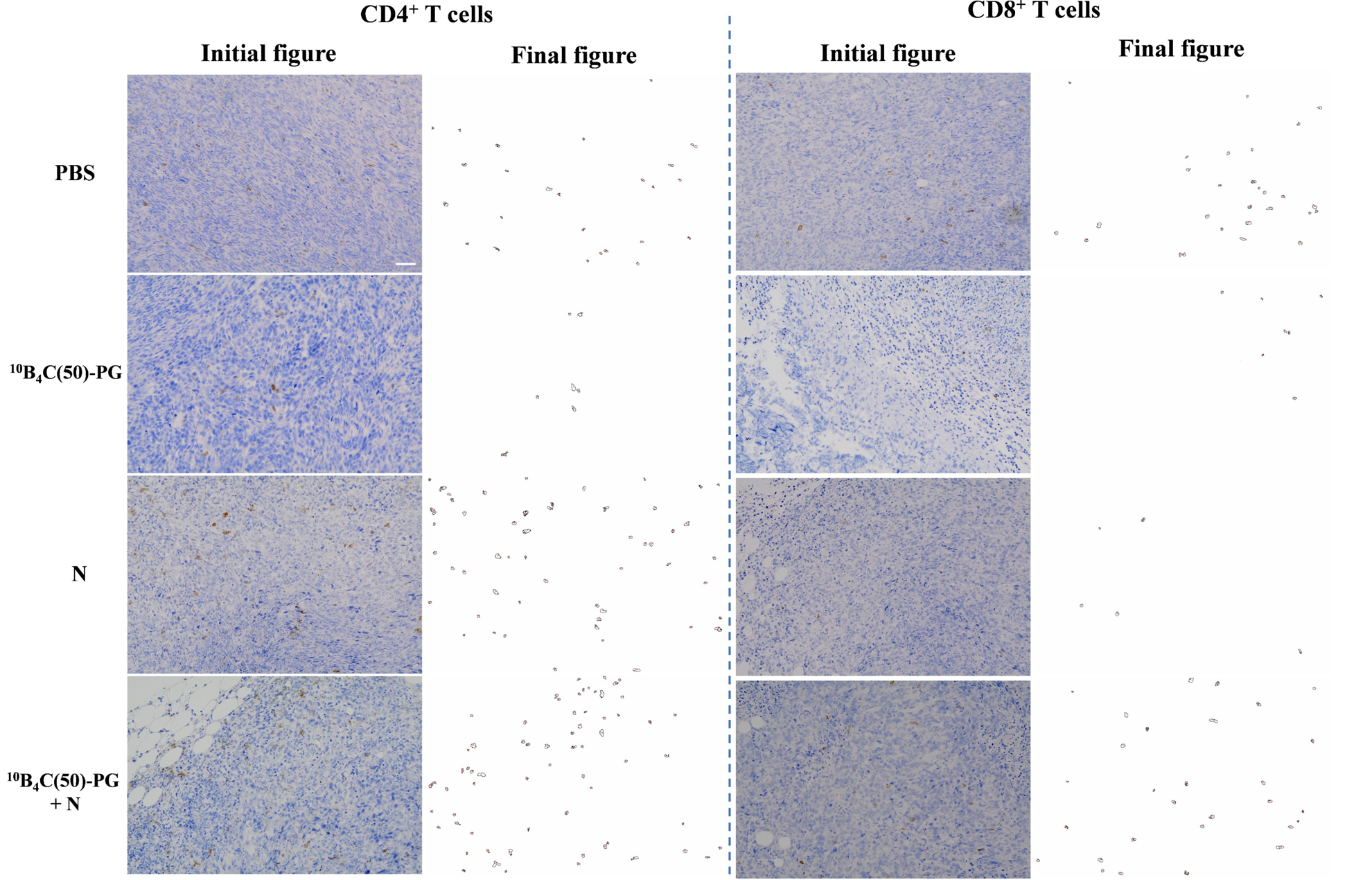

**Supplementary Fig. 34 |** Quantification of CD4^+^ and CD8^+^ T cell number from immunohistochemistry images after recognizing T cells using machine learning tool FIJI. Examples to recognize CD4^+^ and CD8^+^ T cells in immunohistostaining by using machine learning tool (Trainable Weka Segmentation) in FIJI. Scale bar, 50 µm.

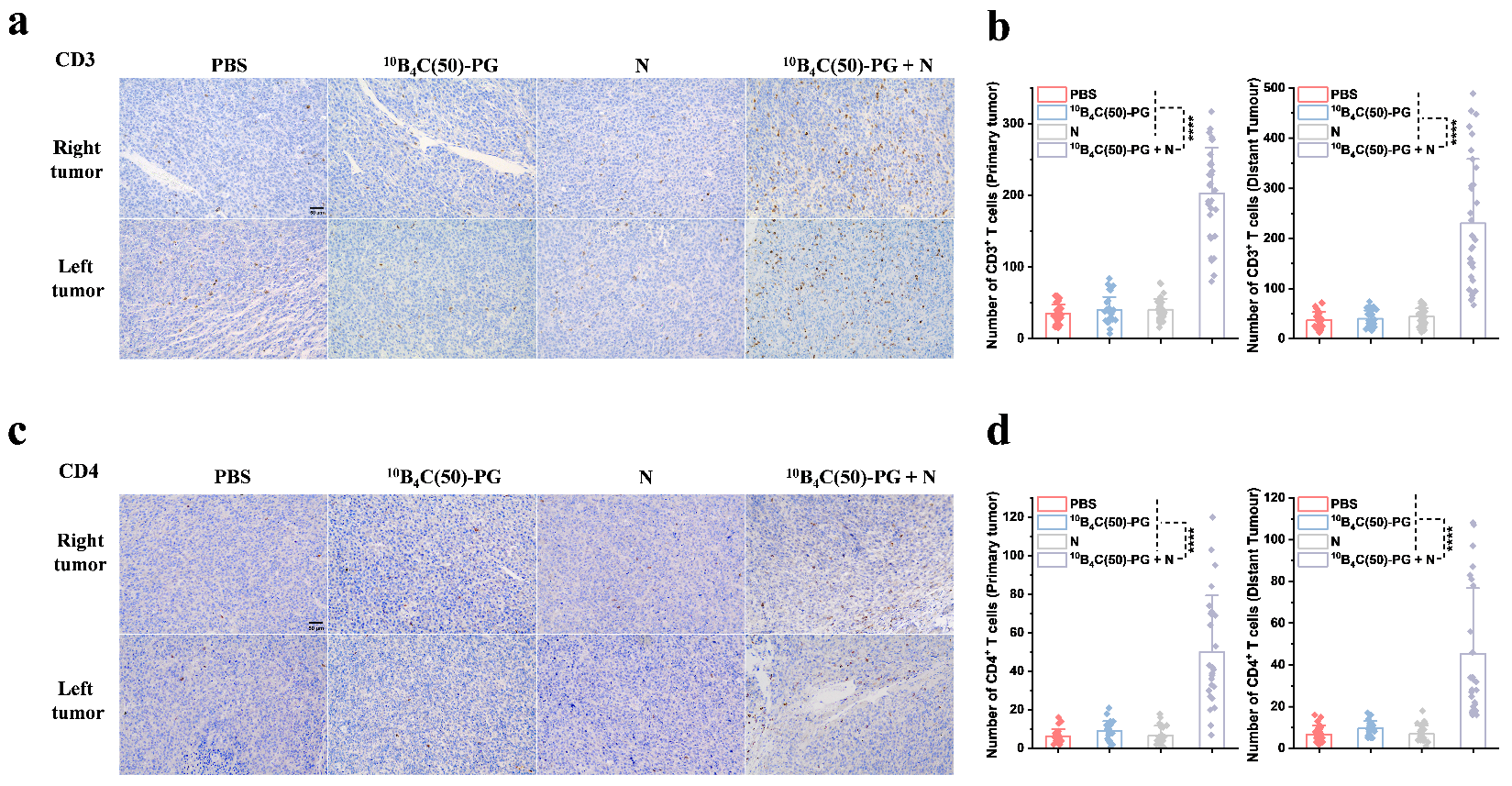

**Number of CD4^+^ T cells (Primary Tumour)**

**Number of CD3^+^ T cells (Primary Tumour)**

**Left tumour**

**Right tumour**

**Right tumour**

**Left tumour**

**Supplementary Fig. 35 | a, b,** Quantification of CD3^+^ cell number from immunohistochemistry images after recognizing T cells using machine learning tool FIJI in tumour on day 22 in Supplementary Fig. 32b and c (*n* = 32). Scale bar, 50 um **c, d,** Quantification of CD4^+^ cell number (*n* = 25).

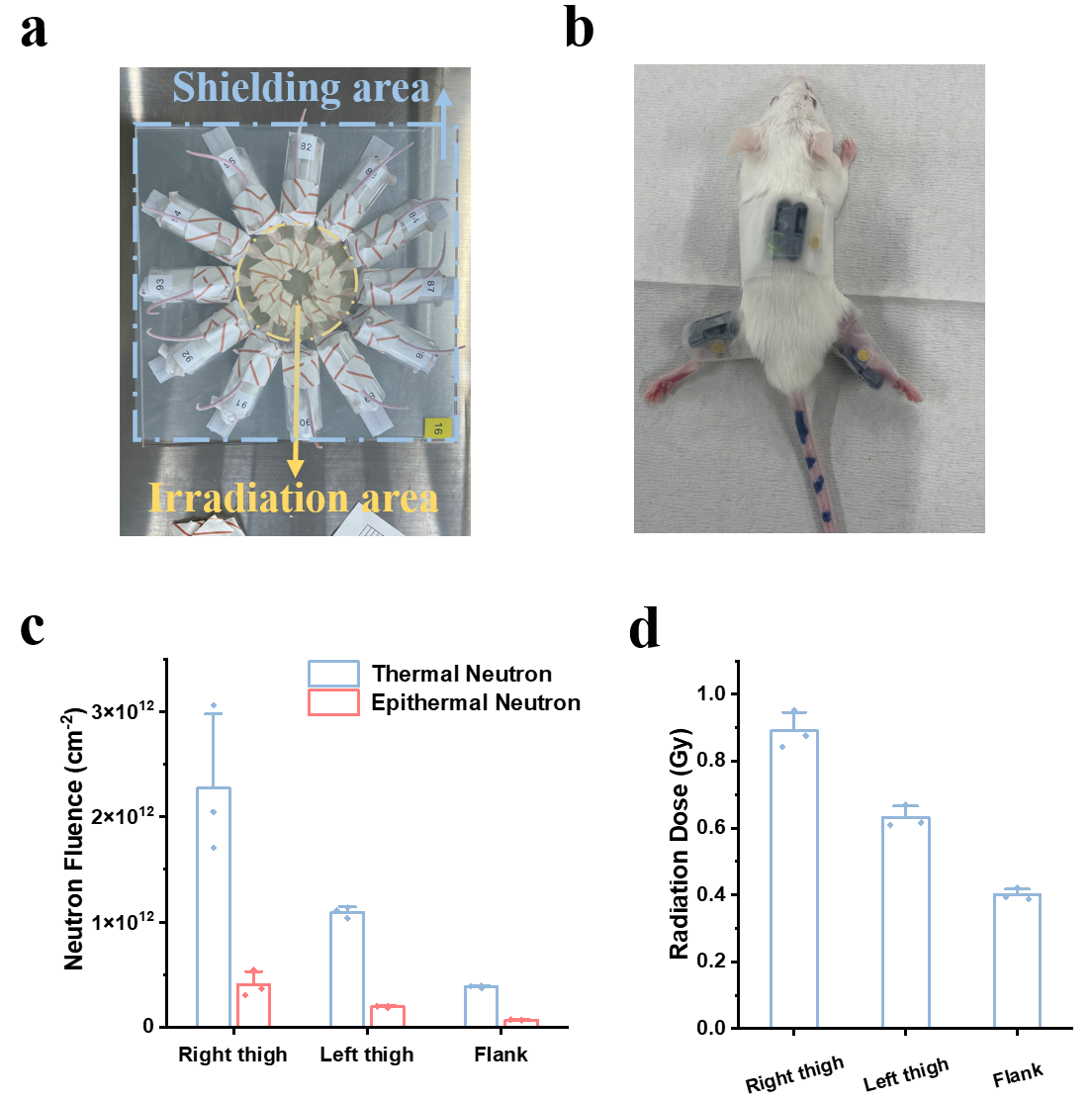

**Supplementary Fig. 36 | Quantification of neutron fluence and radiation dose on right thigh, left thigh and flank after standard neutron irradiation. a**, LiF shielding area and irradiation area for tumour on right thigh when irradiating. **b**, TLD (Thermoluminescent dosimeters, black one) and gold were attached on the right thigh, left thigh and flank of mice to detect the neutron fluence and radiation dose. **c**, Thermal and Epithermal neutron fluence on three places when irradiating (*n* = 3). **d**, Radiation dose when irradiating (*n* = 3).

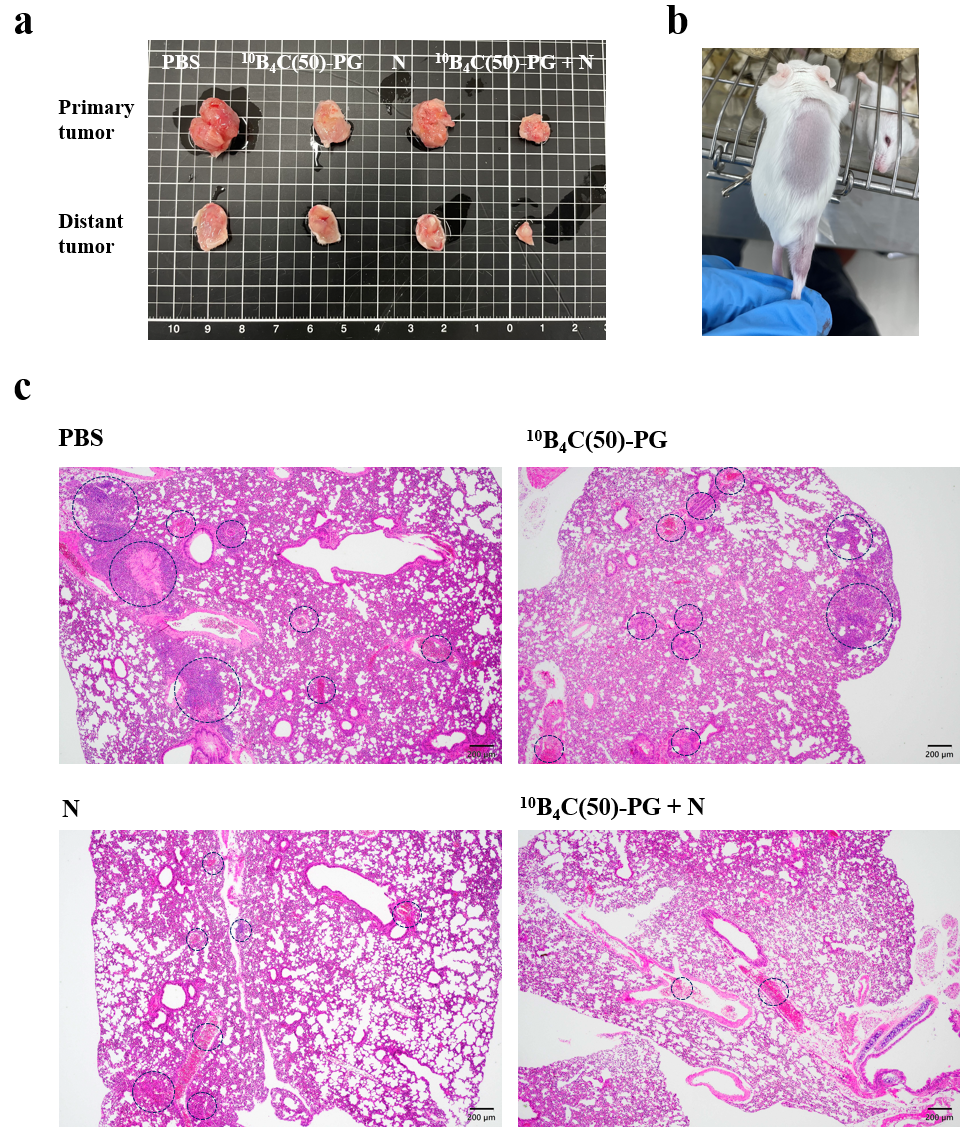

**Primary tumour**

**Distant tumour**

**Supplementary Fig. 37 | a**, Dissected tumour on right thigh and flank of mice in each group after 21 days post irradiation in Fig. 3d and 3e. **b,** Image of cured mice after 21 days post irradiation. **c,** Hematoxylin and eosin staining of histology sections from lung. Scale bar, 200 µm.

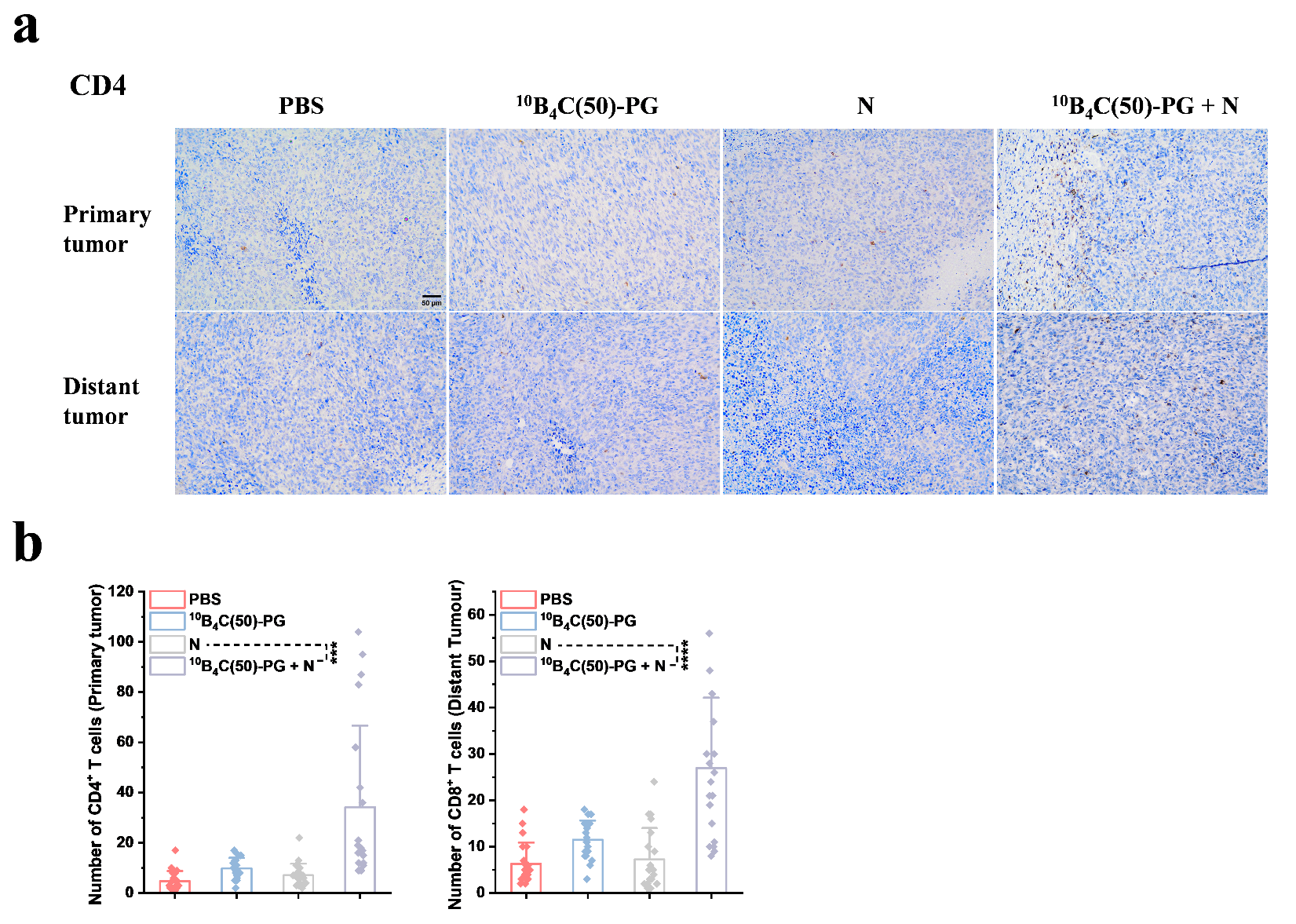

**Number of CD4^+^ T cells (Distant Tumour)**

**Number of CD4^+^ T cells (Primary Tumour)**

**Distant tumour**

**Primary tumour**

**Supplementary Fig. 38 | a**, **b**, Quantification of CD4^+^ cell number from immunohistochemistry images after recognizing T cells using machine learning tool FIJI in tumour on day 22 in Fig. 3d and 3e (*n* = 20). Scale bar, 50 um Statistical analysis with one-way ANOVA post Bonferroni test; *p < 0.05, **p < 0.01, ***p < 0.001, ****p < 0.0001.

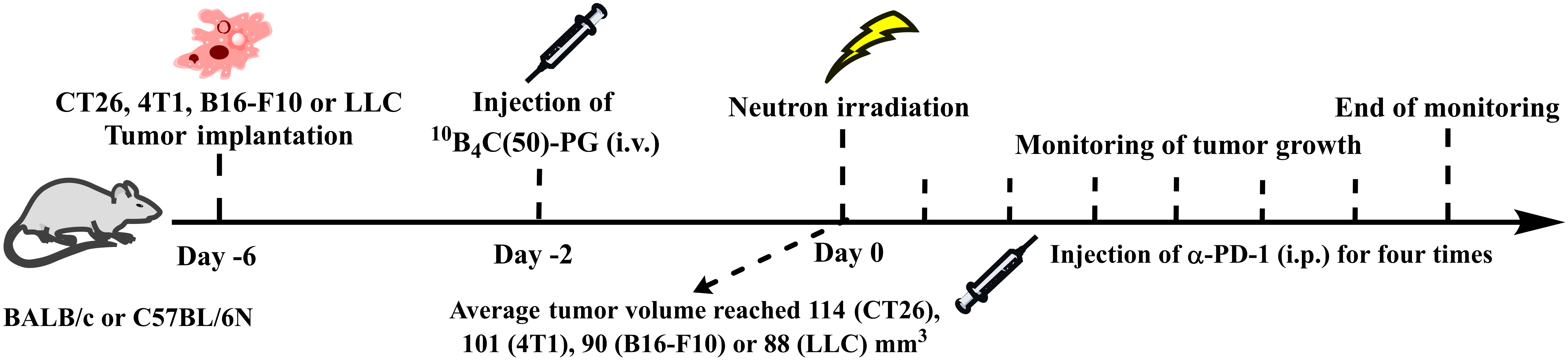

**Monitoring tumour growth**

**CT26, 4T1, B16-F10 and LLC Tumour implantation**

**Supplementary Fig. 39 |** Schematic of CT26, 4T1, B16-F10 and LLC tumour treatment process by *in vivo* BNCT with a dosage of 2.5 mg [^10^B]/kg (mouse) followed by immunotherapy with αPD-1 at a dose of 200 μg / mouse for four times (total dose: 800 μg / mouse).

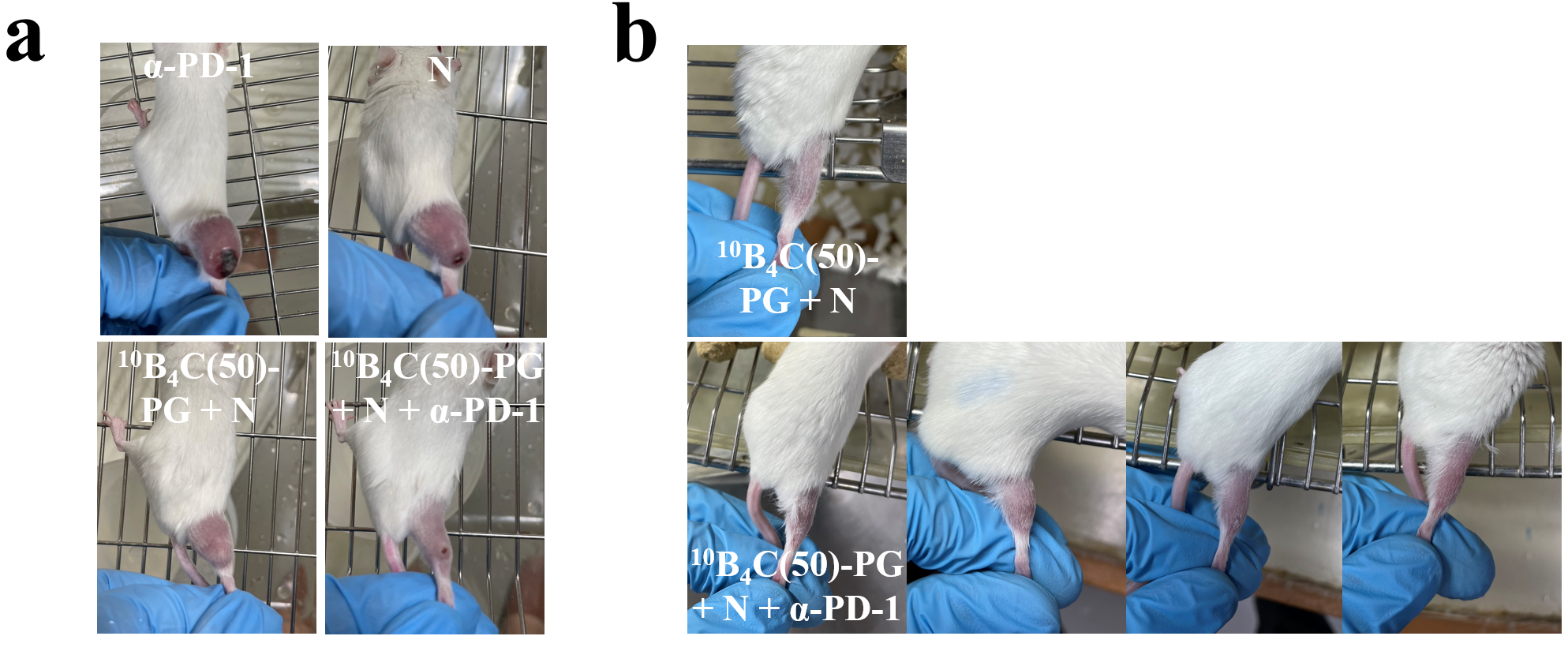

**Supplementary Fig. 40 | a**, Typical tumour images of mice at four groups after 18 days post irradiation in Fig. 3h. **b**, Images of all the survived mic on day 327 in Fig. 3i.

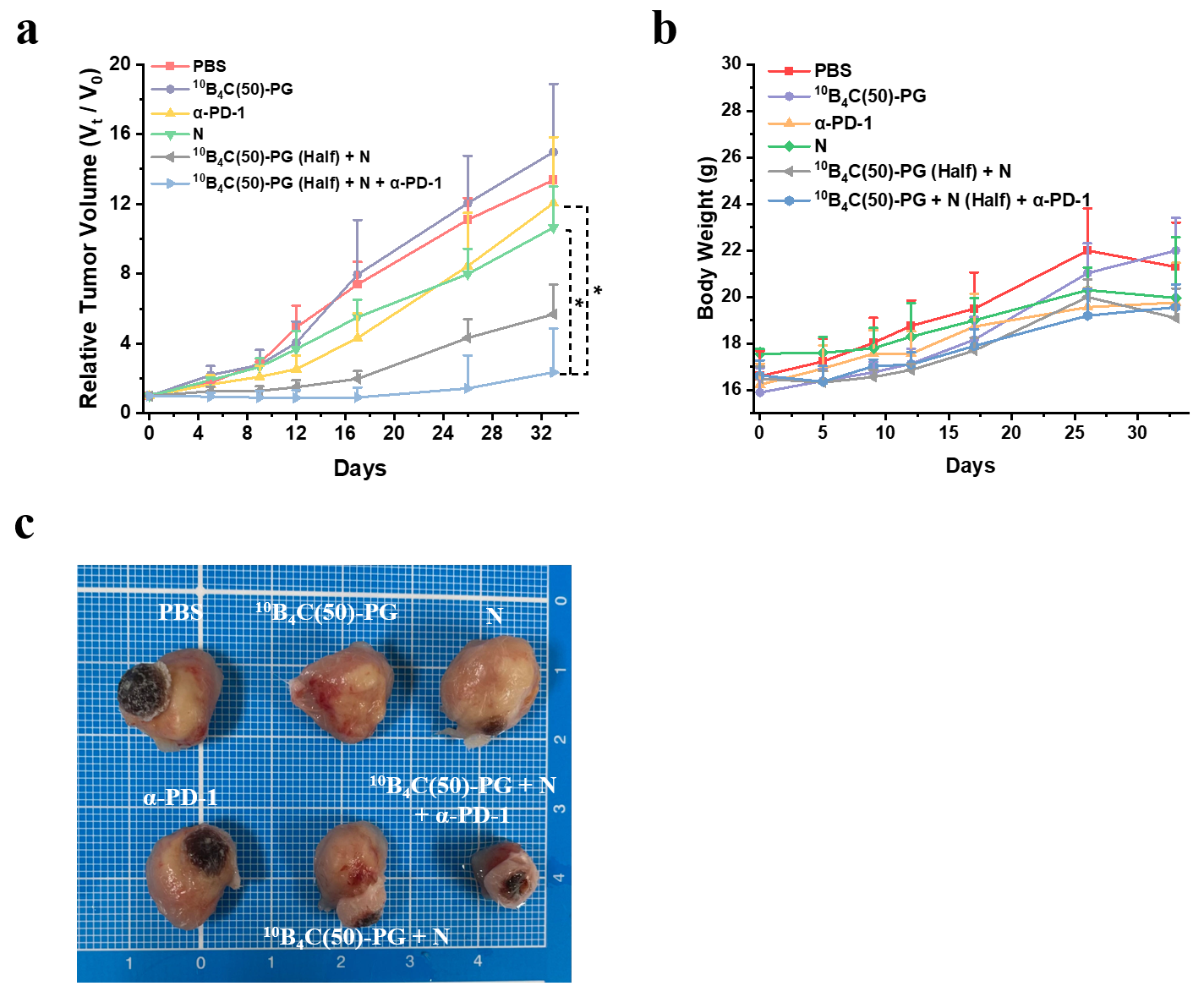

**Relative Tumour Volume (*V*_t_/*V*_0_)**

**Supplementary Fig. 41 |** **4T1 tumour treatment process by BNCT at a dosage of 2.5 mg [^10^B_4_C(50)-PG]/kg (mouse) with αPD-1. a**, Relative 4T1 tumour volume monitored for 32 days. **b**, Time course of body weights of mice. **c,** Typical tumour images of mice in six groups at day 33. The mice in PBS, ^10^B_4_C(50)-PG and N groups were shared from Fig. 2g since these experiments were performed at the same time. Data are given as the mean ± SD (*n* = 3). Statistical analysis of relative tumour volume with one-way ANOVA post Bonferroni test; *p < 0.05, **p < 0.01, ***p < 0.001, ****p < 0.0001.

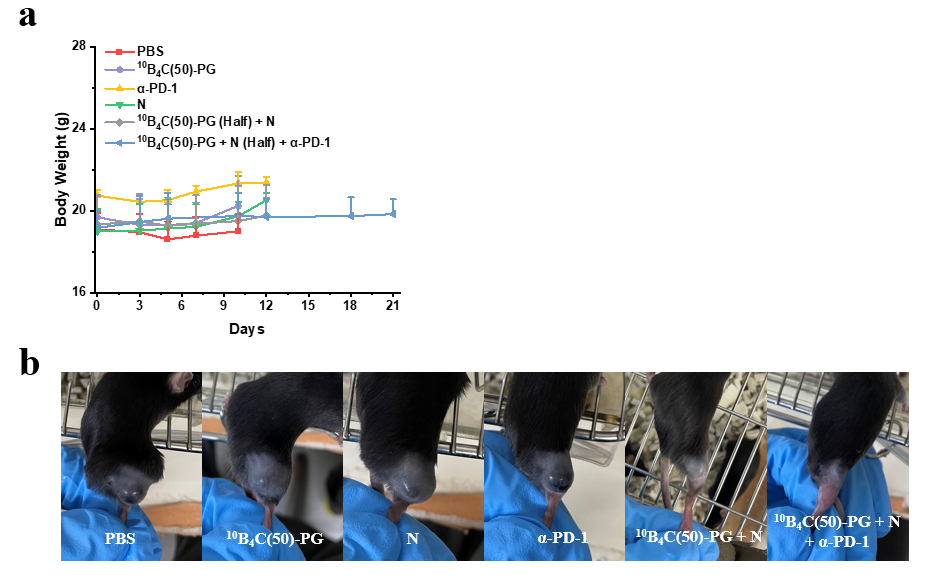

**Supplementary Fig. 42 | B16-F10 tumour treatment process by BNCT at a dosage of 2.5 mg [^10^B_4_C(50)-PG]/kg (mouse) with α-PD-1. a**, Time course of body weights of mice in Fig. 3j. **b,** Typical tumour images of mice in six groups at day 10. The mice in PBS, ^10^B_4_C(50)-PG and N groups were shared from Fig. 2h since these experiments were performed at the same time. Data were given as the mean ± SD (*n* = 5 in αPD-1 group and *n* = 6 in others).

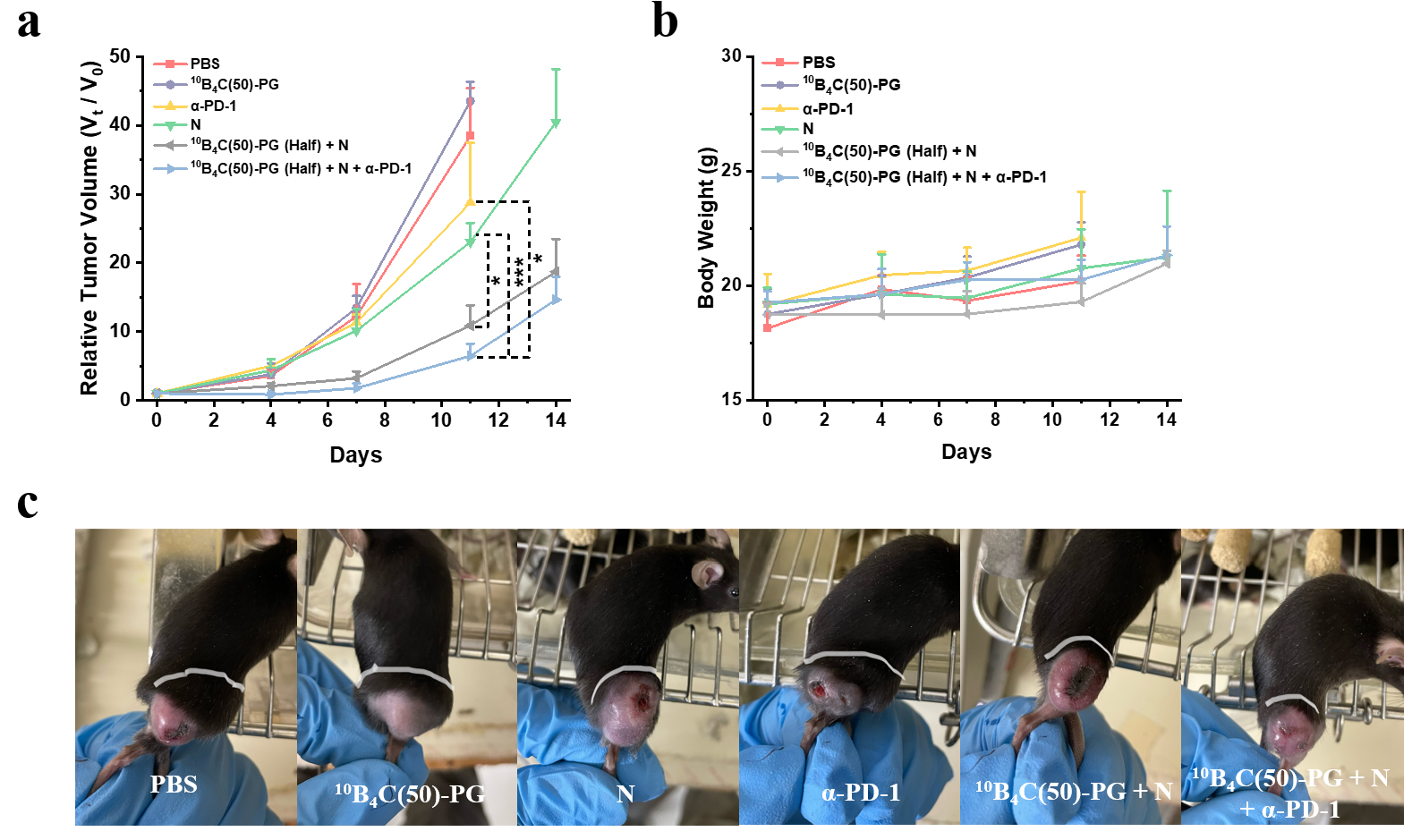

**Relative Tumour Volume (*V*_t_/*V*_0_)**

**Supplementary Fig. 43 | LLC tumour treatment process by BNCT at a dosage of 2.5 mg [^10^B_4_C(50)-PG]/kg (mouse) with α-PD-1. a**, Relative LLC tumour volume monitored for 14 days. **b**, Time course of body weights of mice. **c,** Typical tumour images of mice in six groups at day 11. The mice in PBS, ^10^B_4_C(50)-PG and N groups were shared from Fig. 2i since these experiments were performed at the same time. Data are given as the mean ± SD (*n* = 3). Statistical analysis of relative tumour volume with one-way ANOVA post Bonferroni test; *p < 0.05, **p < 0.01, ***p < 0.001, ****p < 0.0001.

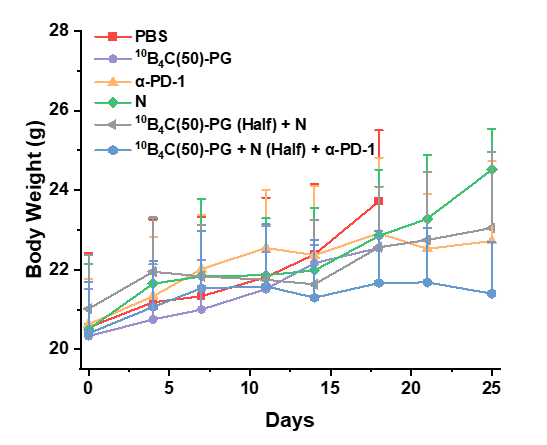

**Supplementary Fig. 44 |** Time course of body weights of mice in Fig. 3h.

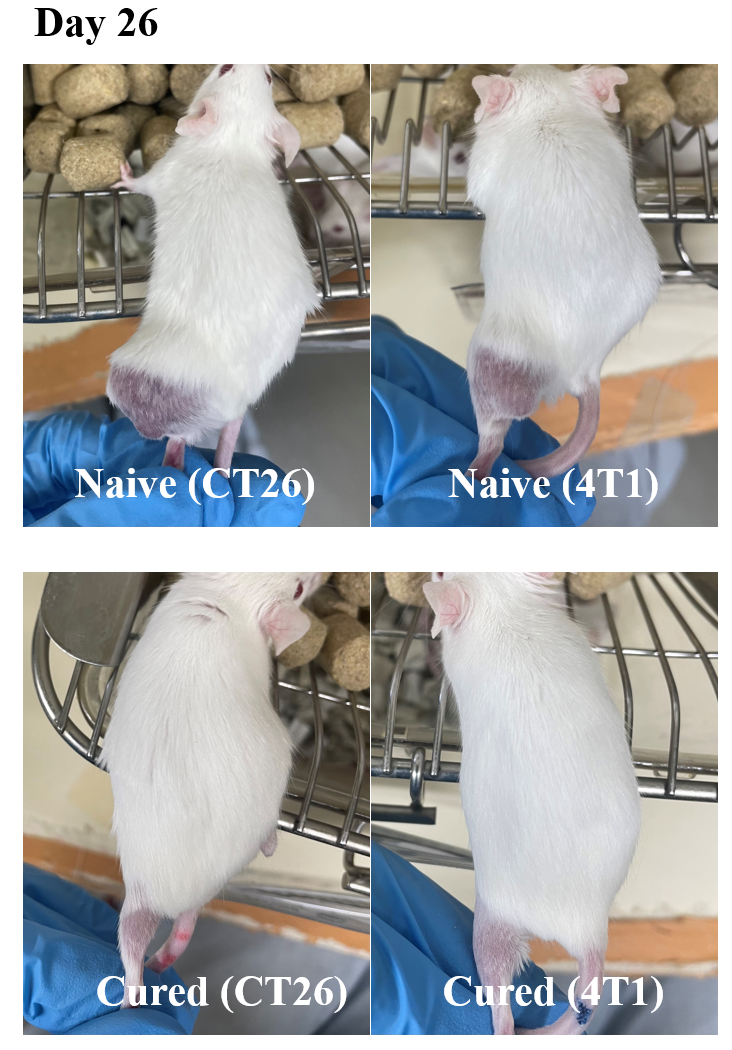

**Supplementary Fig. 45 |** Typical images of tumour mice in naive and cured groups after 26 days post implantation of CT26 and 4T1 cells in Fig. 4a.

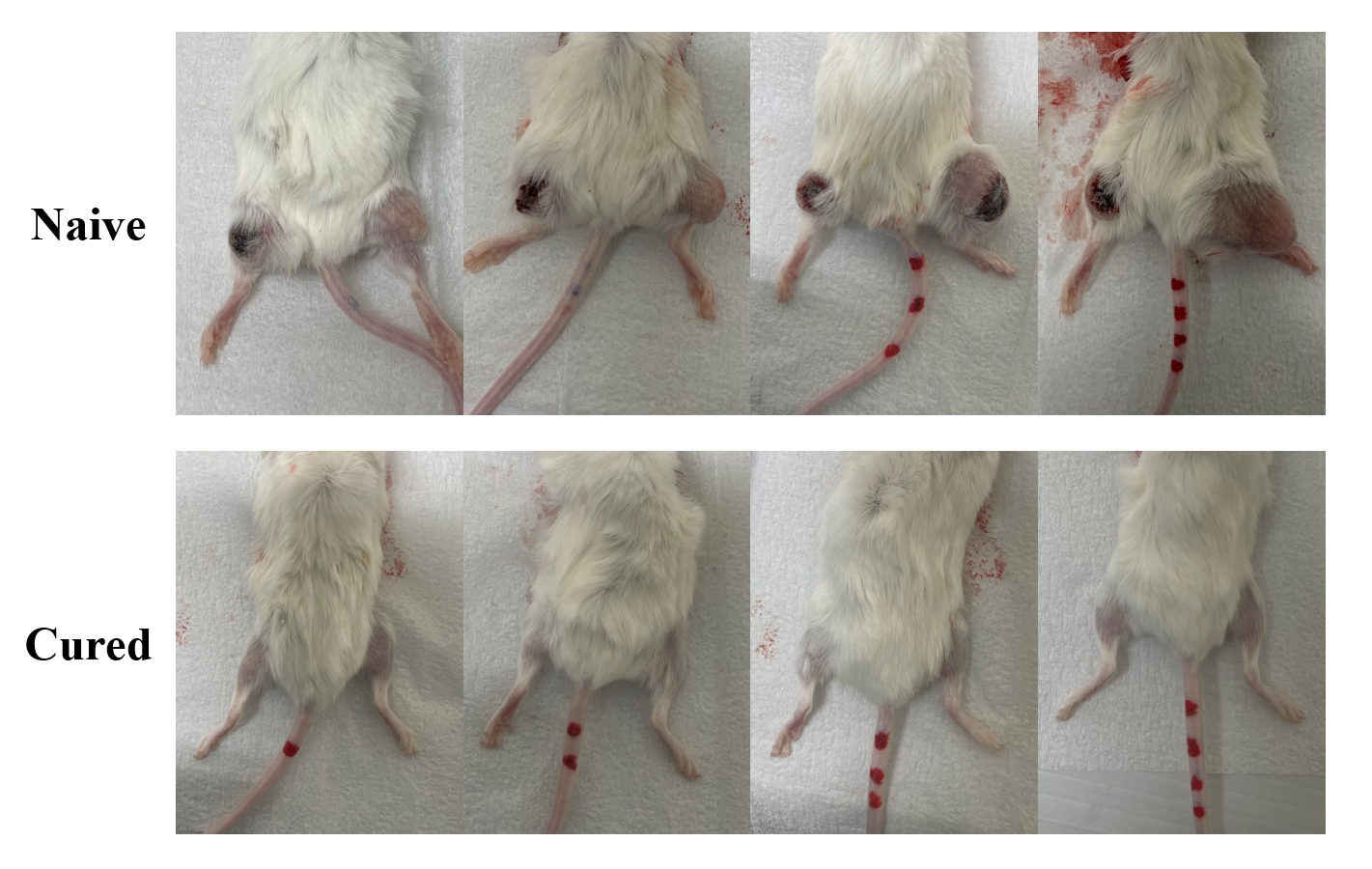

**Supplementary Fig. 46 |** Typical images of tumour mice, Naive (CT26 + 4T1) and Cured (CT26 + 4T1), with CT26 and 4T1 cells implanted simultaneously in the right and left thighs, respectively, after 33 days post implantation in Fig. 4d.

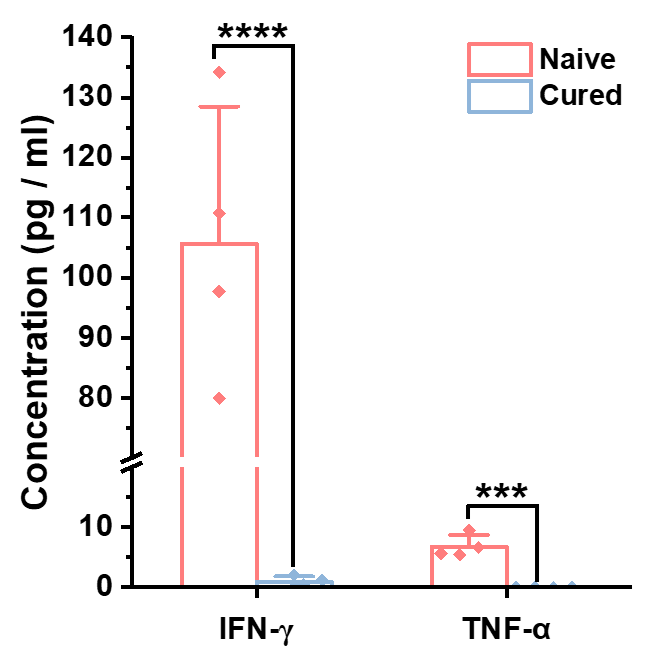

**Supplementary Fig. 47 |** IFN-γ and TNF-α concentrations in serum from cured and naive mice after 33 days post CT26 and 4T1 implantations in Fig. 4d (*n* = 4).

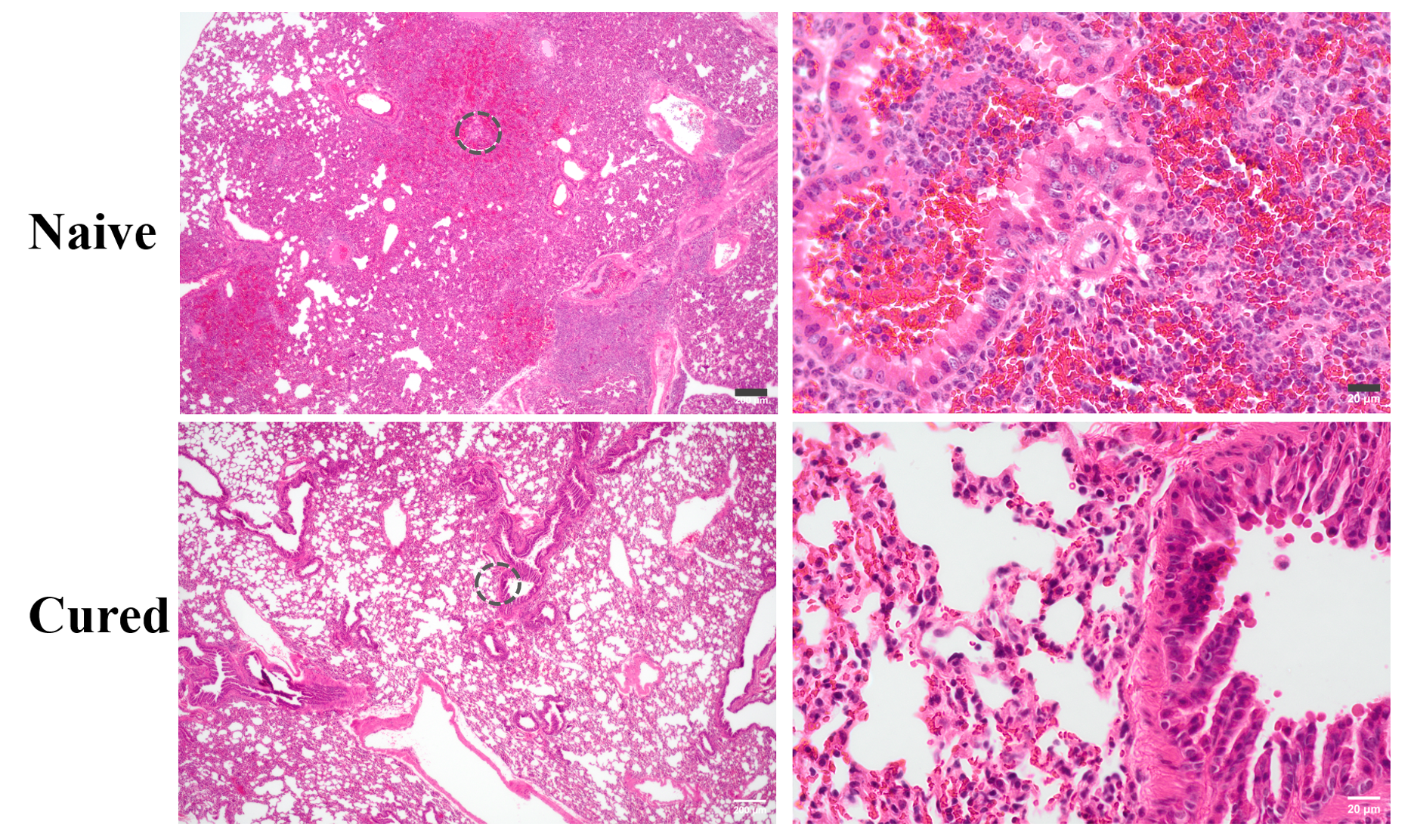

**Supplementary Fig. 48 |** Hematoxylin and eosin staining of histology sections from lung after 33 days post both CT26 and 4T1 implantation in Fig. 4d. (Scale bar, 200 µm in left images and 20 µm in right magnified images).

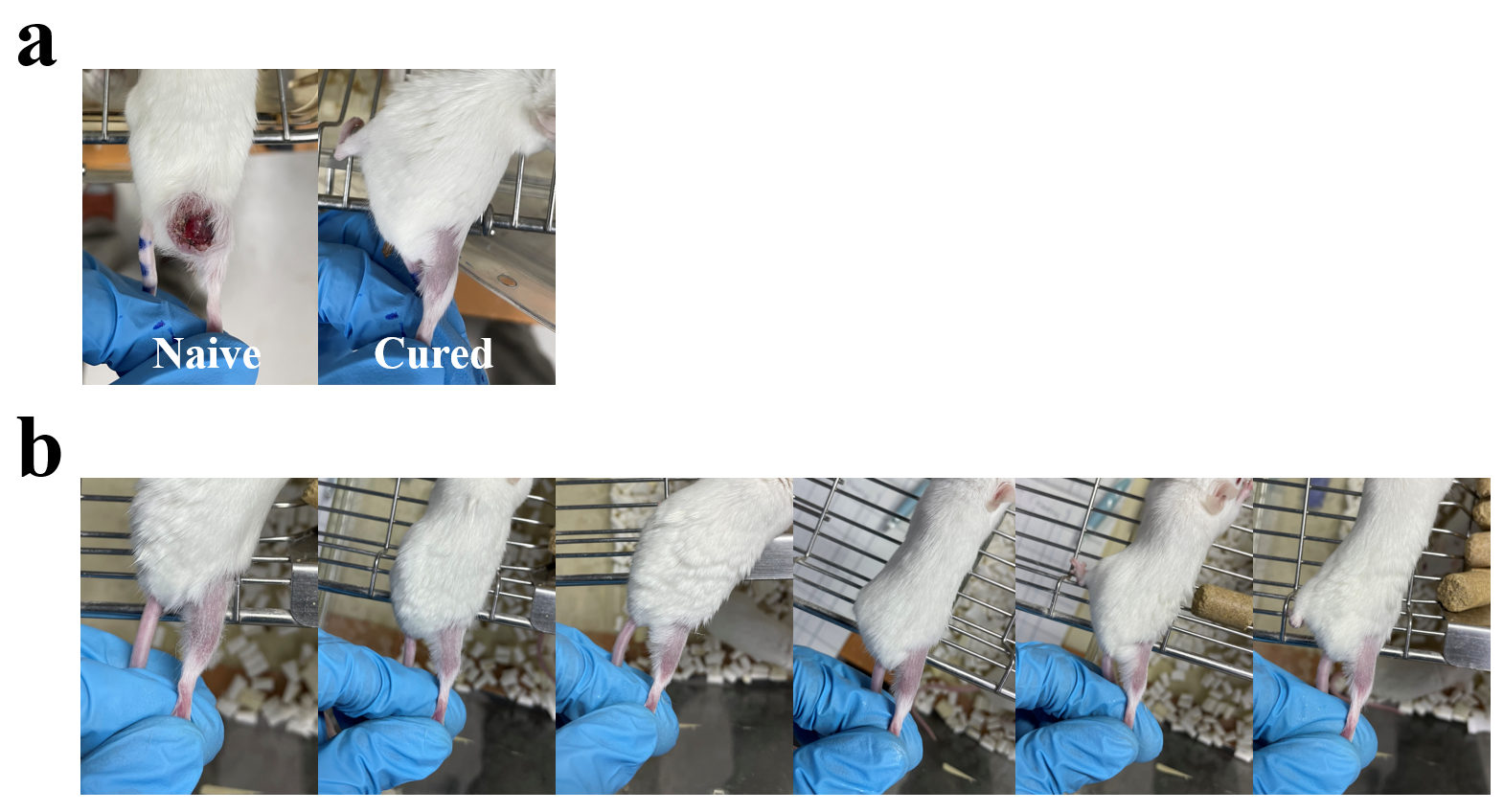

**Supplementary Fig. 49 | a**, Typical tumour images of Naive (Meth-A) and Cured (Meth-A) mice after 34 days post Meth-A implantation in the right leg in Fig. 4e. **b**, Typical tumour images of Cured (Meth-A) mice after 149 days post Math-A implantation in Fig. 4f.

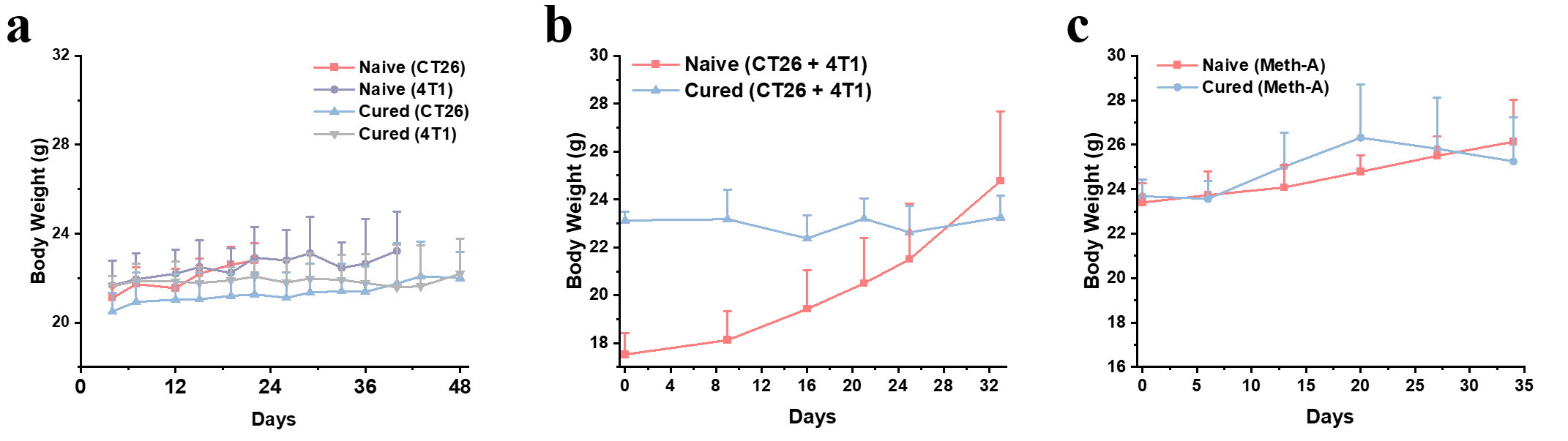

**Supplementary Fig. 50 |** Time course of body weights of mice in **a** (Fig. 4b), **b** (Fig. 4d) and **c** (Fig. 4f).

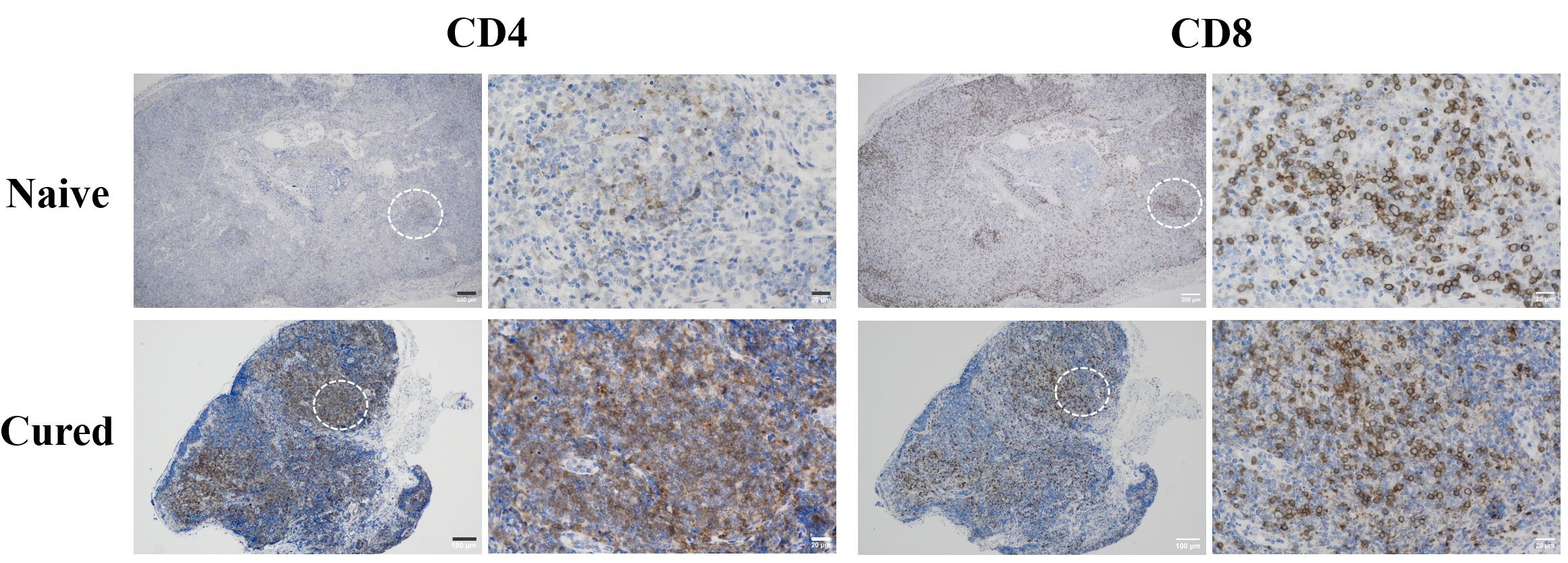

**Supplementary Fig. 51 |** CD4 and CD8 expression in lymph nodes in Naive (CT26 + 4T1) and Cured (CT26 + 4T1). (Scale bar, 200 µm in top left images, 100 µm in bottom left images and 20 µm in right magnified images, respectively).

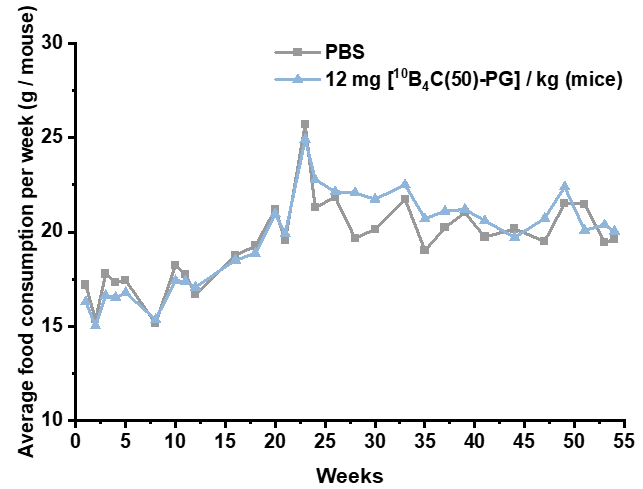

**Supplementary Fig. 52 |** Average food consumption of PBS and ^10^B_4_C(50)-PG administered mice (*n* = 4 and 5, respectively) monitored for 53 weeks.

**Supplementary Table 1** Comparison of various boron-10 drugs injected intravenously to cancered mice for BNCT.

| Boron drug | Hydrody-namic Size (nm) | **Tumour model and animal** | Tumour stage | Drug dosage  (mg / kg) | ^10^B dosage  (mg / kg) | ^10^B in tumour (ppm) | Tumour / blood ^10^B ratio | Delivery  efficiency  (%ID / g) | Delivery  efficiency (%ID) | Thermal neutron fluence  (cm^-2^) | Tumour growth inhibition (%) | Tumour eradication rate | Ref |
| --- | --- | --- | --- | --- | --- | --- | --- | --- | --- | --- | --- | --- | --- |
| Molecular drug | | | | | | | | | | | | | |
| cRGD- MID-BSA | **NA** | **Human glioblastoma adenocarcinoma (****U87MG) subcutaneously inoculated to BALB/c nude mice** | **Volume:**  **~100 mm³**  **Mass:**  **~0.10 g** | **239** | **7.5** | **4.0** | **0.8** | **3.00** | **0.30** | **1.6**  **× 10^12^** | **61** | **0** | **^2^** |
| Fructose-  BPA complex | **NA** | **Murine colon adenocarcinoma (CT26) subcutaneously inoculated to BALB/c mice** | **Volume:**  **~200 mm³**  **Mass:**  **~0.20 g** | **1793** | **24.0** | **16.8** | **2.5** | **3.50** | **0.70** | **3.8**  **× 10^12^** | **64**  **(Initial tumour volume**  **35 mm^3^)** | **0** | **^3^** |
| Poly(vinyl alcohol)-BPA complex | **NA** | **Murine colon adenocarcinoma (CT26) subcutaneously inoculated to BALB/c mice** | **Volume:**  **~200 mm³**  **Mass:**  **~0.20 g** | **1150** | **24.0** | **33.6** | **1.8** | **7.00** | **1.40** | **3.8**  **× 10^12^** | **94**  **(Initial tumour volume**  **35 mm^3^)** | **0** | **^3^** |
| PVA-D-BPA | **NA** | **Murine colon adenocarcinoma (CT26) subcutaneously inoculated to BALB/c mice** | **Volume:**  **~200 mm³**  **Mass:**  **~0.20 g** | **1150** | **24.0** | **60.0** | **2.1** | **12.00** | **2.40** | **2.6**  **× 10^12^** | **99**  **(Initial tumour volume**  **NA)** | **5/6** | **^4^** |
| BPA-Tyr | **NA** | **Human pancreas adenocarcinoma (AsPC-1) subcutaneously inoculated to BALB/c nude mice** | **NA** | **370** | **10.3** | **1.4** | **2.0** | **NA** | **NA** | **NA**  **(No BNCT)** | **NA**  **(No BNCT)** | **NA**  **(No BNCT)** | **^5^** |
| BSH-polymer conjugates | **NA** | **Murine colon adenocarcinoma (CT26) subcutaneously inoculated to BALB/c mice** | **Volume:**  **~50 mm³**  **Mass:**  **~0.05 g** | **685** | **60.3** | **128.0** | **21.0** | **7.00** | **0.35** | **1.9**  **× 10^12^** | **93** | **0** | **^6^** |
| PHPMA-PL-BSH | **NA** | **Murine colon adenocarcinoma (CT26) subcutaneously inoculated to BALB/c mice** | **Volume:**  **~200 mm³**  **Mass:**  **~0.20 g** | **328** | **10.0** | **15.8** | **3.0** | **7.92** | **1.58** | **3.7**  **× 10^12^** | **52**  **(Initial tumour volume**  **50 mm^3^)** | **0** | **^7^** |
| Fluoroboron-otyrosine | **NA** | **Murine melanoma (B16-F10) subcutaneously inoculated to C57BL/6 mice** | **Volume:**  **~50 mm³**  **Mass:**  **~0.05 g** | **1000** | **10.0** | **3.6** | **3.2** | **3.04** | **0.15** | **1.8**  **× 10^14^** | **69** | **0** | **^8^** |
| BSH-based PBC-IP | **NA** | **Human glioblastoma adenocarcinoma (U87MG) subcutaneously inoculated to BALB/c nude mice** | **Volume:**  **~75 mm³**  **Mass:**  **~0.75 g** | **245** | **25.0** | **30.0** | **0.9** | **7.06** | **0.53** | **3.6**  **× 10^12^** | **64** | **0** | **^9^** |
| ^64^Cu-DOTA-cRGD(_D_-BPA)K | **NA** | **Human glioblastoma adenocarcinoma (U87MG) subcutaneously inoculated to BALB/c nude mice** | **NA** | **2093** | **3.5** | **1.3** | **2.1** | **3.00** | **NA** | **NA**  **(No BNCT)** | **NA**  **(No BNCT)** | **NA**  **(No BNCT)** | **^10^** |
| [^18^F]BBPA-PET | **NA** | **Murine melanoma (B16-F10) subcutaneously inoculated to C57BL/6 mice** | **NA** | **250** | **4.3** | **3.6** | **5.2** | **5.23** | **NA** | **3.4**  **× 10^12^** | **95**  **(Initial tumour volume**  **100 mm^3^)** | **1/6** | **^11^** |
| S-BPA | **NA** | **Human glioblastoma adenocarcinoma (U87MG) intracranially inoculated to BALB/c nude mice** | **NA** | **1415** | **29.9** | **28.0** | **3.8** | **4.68** | **NA** | **NA**  **(No BNCT)** | **NA**  **(No BNCT)** | **NA**  **(No BNCT)** | **^12^** |
| Organic (Soft) nanoparticles | | | | | | | | | | | | | |
| BSH-DSBL liposomes | **127** | **Murine colon adenocarcinoma (CT26) subcutaneously inoculated to BALB/c mice** | **Volume:**  **~150 mm³**  **Mass:**  **~0.15 g** | **336** | **15.0** | **29** | **0.7** | **10.78** | **1.62** | **1.7**  **× 10^12^** | **100** | **5/5** | **^13^** |
| Carborane liposomes | **115** | **Murine colon adenocarcinoma (CT26) subcutaneously inoculated to BALB/c mice** | **Volume:**  **~50 mm³**  **Mass:**  **~0.05 g** | **190** | **21.0** | **NA** | **NA** | **NA** | **NA** | **NA** | **92** | **0** | **^14^** |
| TAC/MAC liposomes | **122** | **Murine mammary carcinoma (EMT6) subcutaneously inoculated to BALB/c mice** | **Volume:**  **~115 mm³**  **Mass:**  **~0.12 g** | **878** | **37.1** | **67.8** | **1.9** | **10.00** | **1.17** | **3.4**  **× 10^12^** | **90** | **0** | **^15^** |
| Boronated liposomes | **100** | **Murine mammary carcinoma (4T1) subcutaneously inoculated to BALB/c mice** | **Volume:**  **~70 mm³**  **Mass:**  **~0.07 g** | **500** | **NA** | **17.4** | **4.9** | **5.49** | **0.38** | **3.4**  **× 10^12^** | **91** | **2/9** | **^16^** |
| BSH-based immunoliposomes | **130** | **Human glioblastoma adenocarcinoma (U87MG) intracranially inoculated to BALB/c nude mice** | **NA** | **NA** | **35.0** | **28.4** | **2.8** | **NA** | **NA** | **NA**  **(No BNCT)** | **NA**  **(No BNCT)** | **NA**  **(No BNCT)** | **^17^** |
| BSH-based TF-PEG liposomes | **123** | **Murine colon adenocarcinoma (CT26) subcutaneously inoculated to BALB/c mice** | **Volume:**  **~88 mm³**  **Mass:**  **~0.09 g** | **2976** | **35.0** | **35.5** | **2.5** | **4.73** | **0.43** | **2.0**  **× 10^12^** | **100** | **5/5** | **^18^** |
| BPN-based  PLGA−mPEG micelles | **100** | **Murine melanoma (B16-F10) subcutaneously inoculated to C57BL/6 mice** | **Volume:**  **55 mm³**  **Mass:**  **~0.06 g** | **1562** | **47.7** | **23.3** | **33.9** | **2.57** | **0.15** | **1.0**  **× 10^12^** | **61** | **0** | **^19^** |
| DSPE-BCOP-5T COP micelles | **110** | Murine mammary carcinoma (4T1) subcutaneously inoculated to BALB/c mice | **Volume:**  **~50 mm³**  **Mass:**  **~0.05 g** | **439** | **9.3** | **15.8** | **7.5** | **8.49** | **0.42** | **1.0**  **× 10^12^** | **62** | **0** | **^20^** |
| mPEG-b-PMPCB  micelles | **47** | Murine cervical (U14) subcutaneously inoculated to KM mice | **Volume:**  **~110 mm³**  **Mass:**  **~0.11 g** | **127** | **3.7** | **10.7** | **NA** | **12.77** | **1.38** | **NA** | **68** | **0** | **^21^** |
| Ｃarborane-based ^10^B-NapE4 cluster | **103** | Human glioblastoma adenocarcinoma (A549) subcutaneously inoculated to BALB/c nude mice | **Volume:**  **~50 mm³**  **Mass:**  **~0.05 g** | **15** | **1.9** | **22.7** | **28.9** | **59.67** | **2.98** | **NA** | **69** | **0** | **^22^** |
| BPA-containing polydopamine NPs | **189** | Human glioblastoma adenocarcinoma (U87MG) intracranially inoculated to NCG mice | **NA** | **200** | **1.8** | **29.7** | **6.8** | **71.74** | **NA** | **NA** | **48** | **5/8** | **^23^** |
| BPA-containing polydopamine NPs | **195** | Murine melanoma (B16-F10) subcutaneously inoculated to C57BL/6 mice | **NA** | **200** | **1.8** | **31.5** | **4.9** | **79.55** | **NA** | **NA** | **63**  **(Initial tumour volume**  **100 mm^3^)** | **0** | **^24^** |
| iRGD-PEG-PCCL-B micelles | **NA** | Human glioblastoma adenocarcinoma (A549) subcutaneously inoculated to BALB/c nude mice | **Volume:**  **~175 mm³**  **Mass:**  **~0.18 g** | **525** | **20.0** | **40.0** | **14.1** | **12.50** | **2.19** | **NA**  **(No BNCT)** | **NA**  **(No BNCT)** | **NA**  **(No BNCT)** | **^25^** |
| PVA/BA NPs | **134** | Human squamous cell carcinoma (SAS) subcutaneously inoculated to C.B17/Icr-Prkdcscid/CrlNarl male mice | **Volume:**  **32 mm³**  **Mass:**  **~0.03 g** | **51** | **5.0** | **NA** | **NA** | **NA** | **NA** | **1.2**  **× 10^12^** | **66** | **0** | **^26^** |
| ^10^B/siNC NPs | **97** | Murine mammary carcinoma (4T1) subcutaneously inoculated to BALB/c mice | **Volume:**  **~100 mm³**  **Mass:**  **~0.10 g** | **567** | **10.0** | **42.7** | **5.2** | **22.47** | **2.25** | **NA**  **(Two times in one week)** | **80**  **(20 mg / kg of ^10^B dosage)** | **0** | **^27^** |
| Inorganic (hard) nanoparticles | | | | | | | | | | | | | |
| Carborane-appended carbon nanotubes | **NA** | Murine mammary carcinoma (EMT6) subcutaneously inoculated to BALB/c mice | **NA** | **230** | **NA** | **4.4** | **1.8** | **NA** | **NA** | **NA**  **(No BNCT)** | **NA**  **(No BNCT)** | **NA**  **(No BNCT)** | **^28^** |
| BSH-LDH NPs | **100** | Human glioblastoma adenocarcinoma (U87MG) subcutaneously inoculated to BALB/c nude mice | **Volume:**  **~100 mm³**  **Mass:**  **~0.10 g** | **110** | **NA** | **7.7** | **3.5** | **17.50** | **1.75** | **NA**  **(No BNCT)** | **NA**  **(No BNCT)** | **NA**  **(No BNCT)** | **^29^** |
| ^10^BSGRF NPs | **90** | Murine glioblastoma (ALTS1C1) intracranially inoculated to C57BL/6J mice | **Volume:**  **~8 mm³** | **100** | **NA** | **50.5** | **2.8** | **NA** | **NA** | **3.6**  **× 10^12^** | **63** | **0** | **^30^** |
| anti-EGFR-^10^BPO_4_ NPs | **110** | Human hypopharyngeal (HTB-43) subcutaneously inoculated to BALB/C nude mice | **Volume:**  **~50 mm³**  **Mass:**  **~0.05 g** | **50** | **16.6** | **63.0** | **4.3** | **19.00** | **0.95** | **3.6**  **× 10^12^** | **89**  **(Initial tumour volume**  **14 mm^3^)** | **0** | **^31^** |
| Phasetransitione lysozyme@Boron nitride NPs | **100** | Murine mammary carcinoma (4T1) subcutaneously inoculated to BALB/c mice | **Volume:**  **~38 mm³**  **Mass:**  **~0.04 g** | **270** | **NA** | **19.6** | **2.7** | **6.20** | **0.26** | **2.3**  **× 10^12^** | **75** | **0** | **^32^** |
| Boron cluster-containing redox NPs | **36** | Murine colon adenocarcinoma (CT26) subcutaneously inoculated to BALB/c mice | **Volume:**  **~140 mm³**  **Mass:**  **~0.14 g** | **2006** | **15.0** | **20.0** | **6.0** | **5.53** | **0.78** | **1.5**  **× 10^12^** | **77** | **0** | **^33^** |
| PLMB NPs | **108** | Murine cervical (U14) subcutaneously inoculated to KM mice | **Volume:**  **75 mm³**  **Mass:**  **~0.08 g** | **202** | **5.6** | **9.5** | **3.4** | **8.48** | **0.68** | **NA** | **57** | **0** | **^34^** |
| Boron-10 nitride nanosheets | **127** | Murine mammary carcinoma (4T1) subcutaneously inoculated to BALB/c mice | **Volume:**  **50 mm³**  **Mass:**  **~0.05 g** | **90** | **37.8** | **23.2** | **2.4** | **3.17** | **0.16** | **2.3**  **× 10^12^** | **64** | **0** | **^35^** |
| anti-EGFR-Ce^10^B_6_ NPs | **NA** | Human squamous cell carcinoma (HTB-43) subcutaneously inoculated to BALB/c nude mice | **Volume:**  **21 mm³**  **Mass:**  **~0.02 g** | **56** | **15.0** | **151.0** | **4.2** | **50.00** | **1.25** | **3.6**  **× 10^12^** | **72** | **0** | **^36^** |
| Zr-TCPP MOFs | **100** | Human glioblastoma adenocarcinoma (U87MG) intracranially inoculated to BALB/c nude mice | **NA** | **3333** | **50.0** | **13.5** | **3.8** | **NA** | **NA** | **1.1**  **× 10^12^** | **NA** | **3/4** | **^37^** |
| PEG@BCNO NPs | **134** | Human mammary adenocarcinoma (MDA-MB-231) intracranially inoculated to BALB/c nude mice | **Volume:**  **100 mm³**  **Mass:**  **~0.10 g** | **40** | **NA** | **6.0** | **3.5** | **NA** | **NA** | **2.7**  **× 10^12^** | **59** | **0** | **^38^** |
| SP94-LB@BA-MSN NPs | **148** | Human hepatocellular carcinoma (HepG-2) subcutaneously inoculated to BALB/c nude mice | **Volume:**  **~33 mm³**  **Mass:**  **~0.03 g** | **333** | **4.0** | **8.0** | **5.9** | **11.25** | **0.38** | **NA**  **(No BNCT)** | **NA**  **(No BNCT)** | **NA**  **(No BNCT)** | **^39^** |
| Boron quantum dots | **NA** | Murine mammary carcinoma (4T1) subcutaneously inoculated to BALB/c mice | **Volume:**  **~80 mm³**  **Mass:**  **~0.08 g** | **50** | **9.3** | **27.4** | **3.5** | **18.4** | **1.47** | **3.4**  **× 10^12^** | **58** | **0** | **^40^** |
| BATBNs | **9** | Murine mammary carcinoma (4T1) subcutaneously inoculated to BALB/c mice | **Volume:**  **~100 mm³**  **Mass:**  **~0.10 g** | **50** | **9.3** | **66.1** | **5.7** | **35.5** | **3.55** | **3.4**  **× 10^12^** | **80** | **0** | **^41^** |
| BCDs-HSA | **164** | Murine melanoma (B16-F10) subcutaneously inoculated to C57BL/6 mice | **NA** | **210** | **14.9** | **33.5** | **NA** | **9.78** | **NA** | **NA** | **93**  **(Initial tumour volume**  **100 mm^3^)** | **0** | **^42^** |
| DND-PG-PBA-SucMe NPs | **57** | Murine colon adenocarcinoma (CT26) subcutaneously inoculated to BALB/c mice | **NA** | **400** | **7.4** | **14.3** | **6.7** | **9.66** | **NA** | **3.8**  **× 10^12^** | **50** | **0** | **^43^** |
| h-^10^BN-PG NPs | **36** | Murine colon adenocarcinoma (CT26) subcutaneously inoculated to BALB/c mice | **Volume:**  **100 mm³**  **Mass:**  **~0.10 g** | **399** | **58.0** | **80.0** | **0.5** | **6.92** | **0.69** | **3.6**  **× 10^12^** | **100**  **(Initial tumour volume**  **80 mm^3^)** | **3/3** | **^44^** |
| ^10^B_4_C-PG NPs | **73** | Murine colon adenocarcinoma (CT26) subcutaneously inoculated to BALB/c mice | **Volume:**  **~424 mm³**  **Mass:**  **~0.34 g** | **165** | **30.9** | **36.7** | **1.4** | **5.68** | **1.93** | **3.6**  **× 10^12^** | **75** | **0** | **^45^** |
| ^10^B_4_C-PG NPs | **37** | Murine colon adenocarcinoma (CT26) subcutaneously inoculated to BALB/c mice | **Volume:**  **~113 mm³** | **18** | **5.1** | **NA** | **NA** | **NA** | **NA** | **3.6**  **× 10^12^** | **86** | **1/3** | **○** |
| ^10^B_4_C-PG NPs | **45** | Murine colon adenocarcinoma (CT26) subcutaneously inoculated to BALB/c mice | **Volume:**  **~120 mm³**  **Mass:**  **~0.06 g** | **15** | **5.1** | **75.8** | **10.0** | **68.07** | **3.38** | **3.6**  **× 10^12^** | **93**  **（Initial tumour volume**  **140 mm^3^)** | **5/8** | **○** |
| ^10^B_4_C-PG NPs | **61** | Murine colon adenocarcinoma (CT26) subcutaneously inoculated to BALB/c mice | **Volume:**  **~120 mm³**  **Mass:**  **~0.06 g** | **12** | **5.1** | **82.0** | **8.9** | **73.58** | **4.03** | **3.6**  **× 10^12^** | **98**  **（Initial tumour volume**  **140 mm^3^)** | **7/8** | **○** |
| ^10^B_4_C-PG NPs | **89** | Murine colon adenocarcinoma (CT26) subcutaneously inoculated to BALB/c mice | **Volume:**  **~120 mm³**  **Mass:**  **~0.06 g** | **14** | **5.1** | **30.1** | **15.4** | **27.03** | **1.67** | **3.6**  **× 10^12^** | **66**  **（Initial tumour volume**  **140 mm^3^)** | **1/8** | **○** |
| ^10^B_4_C-PG NPs | **123** | Murine colon adenocarcinoma (CT26) subcutaneously inoculated to BALB/c mice | **Volume:**  **~120 mm³**  **Mass:**  **~0.05 g** | **14** | **5.1** | **55.9** | **13.9** | **50.15** | **2.38** | **3.6**  **× 10^12^** | **73**  **（Initial tumour volume**  **140 mm^3^)** | **2/8** | **○** |
| ^10^B_4_C-PG NPs | **61** | Murine mammary carcinoma (4T1) subcutaneously inoculated to BALB/c mice | **Volume:**  **~149 mm³**  **Mass:**  **~0.09g** | **12** | **5.2** | **51.5** | **9.3** | **49.85** | **3.20** | **3.6**  **× 10^12^** | **86**  **(Initial tumour volume**  **97 mm^3^)** | **1/3** | **○** |
| ^10^B_4_C-PG NPs | **61** | Murine skin fibrosarcoma (Meth-A) subcutaneously inoculated to BALB/c mice | **Volume:**  **~154 mm³** | **12** | **5.1** | **NA** | **NA** | **NA** | **NA** | **3.6**  **× 10^12^** | **100** | **5/5** | **○** |
| ^10^B_4_C-PG NPs | **61** | Murine melanoma (B16-F10) subcutaneously inoculated to C57BL/6 mice | **Volume:**  **~143 mm³**  **Mass:**  **~0.10 g** | **12** | **5.1** | **52.6** | **12.5** | **47.68** | **3.48** | **3.6**  **× 10^12^** | **77**  **(Initial tumour volume**  **94 mm^3^)** | **0** | **○** |
| ^10^B_4_C-PG NPs | **61** | Murine Lewis lung carcinoma (LLC) subcutaneously inoculated to C57BL/6 mice | **Volume:**  **~255 mm³**  **Mass:**  **~0.17 g** | **12** | **5.1** | **35.5** | **11.9** | **30.14** | **4.39** | **3.6**  **× 10^12^** | **73**  **(Initial tumour volume**  **71 mm^3^)** | **0** | **○** |
| ^10^B_4_C-PG NPs | **61** | Murine colon adenocarcinoma (CT26) subcutaneously inoculated to BALB/c mice | **Volume:**  **~169 mm³** | **6** | **2.5** | **NA** | **NA** | **NA** | **NA** | **3.6**  **× 10^12^ × 2 in two days** | **91** | **5/6** | **○** |
| ^10^B_4_C-PG NPs | **61** | Murine colon adenocarcinoma (CT26) subcutaneously inoculated to BALB/c mice | **Volume:**  **~122 mm³** | **6** | **2.5** | **NA** | **NA** | **NA** | **NA** | **3.6**  **× 10^12^ × 2 in one week** | **96** | **2/5** | **○** |

NA, not available; ○, present work; If only tumour volume was reported, the weight was approximated using an average tumour density^46^ of 1.0 g/cm³. Tumour weight was only precisely reported in our previous and present works.

**Supplementary Table 2**　Synthesis of Boron Carbide Nanoparticles by Ball Milling

| **Raw material** | **TEM size (nm)** | **Literature** |
| --- | --- | --- |
| Mg, B_2_O_3_,  polyvinyl chloride | 50 – 200 | J. Wang et al., *Ceramics International* 42(2016) 6969 |
| Mg, B_2_O_3_,  carbon powder | 50 – 350 | Gokmese et al., *International Advanced Researches and Engineering Journal* 03(2019) 195 |
| Mg, B_2_O_3_,  graphite powder | 10 – 80 | E. Mohammad Sharifi et al., *Advanced Powder Technology* 22 (2011) 354 |
| Mg, B_2_O_3_, graphite | 100 – 200 | F. Deng et al., *Materials Letters* 60 (2006) 1771 |
| Mg, B_2_O_3_, graphite | 5 – 50 | M.J. Nasr Isfahani et al., *Journal of Alloys and Compounds* 797 (2019) 1348 |

**Supplementary Table 3** Summary of the parameters in the ball-milling syntheses of ^10^B_4_C or B_4_C NPs with 35, 50, 80 and 110 nm sizes.

| NPs | Pretreatment  Duration  (min) | | Mass of reactant (g) | | | Total mass  (g) | Ball to Powder ratio | Rotation speed  (rpm) | Milling time  (h) | Post  treatment | Mass of product  (mg) | Yield  (%) |
| --- | --- | --- | --- | --- | --- | --- | --- | --- | --- | --- | --- | --- |
|  |  |  | ^10^B_2_O_3_  / B_2_O_3_ | Mg | Graphite |  |  |  |  |  |  |  |
| ^10^B_4_C(35) | ^10^B_2_O_3_ | Mg | 0.91 | 1.00 | 0.092 | 2.00 | 30:1  (new) | 800 | 8 | 10.0 g NaCl | 104 | 30.6 |
|  | 60 | 30 |  |  |  |  |  |  |  | 800 rpm, 1 h |  |  |
| ^10^B_4_C(50) | ^10^B_2_O_3_ | Mg | 0.90 | 1.00 | 0.089 | 1.99 | 30:1  (new) | 800 | 8 | 10.0 g NaCl | 112 | 32.9 |
|  | 20 | 30 |  |  |  |  |  |  |  | 800 rpm, 1 h |  |  |
| ^10^B_4_C(80) | ^10^B_2_O_3_ | Mg | 0.84 | 0.97 | 0.084 | 1.89 | 30:1  (new) | 800 | 8 | 9.5 g NaCl | 92 | 30.8 |
|  | – | 30 |  |  |  |  |  |  |  | 800 rpm, 1 h |  |  |
| ^10^B_4_C(110) | ^10^B_2_O_3_ | Mg | 0.90 | 1.00 | 0.089 | 1.99 | 30:1  (worn) | 800 | 8 | 10.0 g NaCl | 124 | 36.5 |
|  | – | 30 |  |  |  |  |  |  |  | 800 rpm, 1 h |  |  |
| B_4_C(35) | B_2_O_3_ | Mg | 0.95 | 1.04 | 0.096 | 2.10 | 30:1  (new) | 800 | 8 | 10.5 g NaCl | 204 | 55.9 |
|  | 60 | 30 |  |  |  |  |  |  |  | 800 rpm, 1 h |  |  |
| B_4_C(50) | B_2_O_3_ | Mg | 0.90 | 1.00 | 0.091 | 2.10 | 30:1  (new) | 800 | 8 | 10.5 g NaCl | 118 | 33.9 |
|  | 20 | 30 |  |  |  |  |  |  |  | 800 rpm, 1 h |  |  |
| B_4_C(80) | B_2_O_3_ | Mg | 1.01 | 1.10 | 0.101 | 2.21 | 30:1  (new) | 800 | 15 | 11.0 g NaCl | 185 | 47.7 |
|  | – | 30 |  |  |  |  |  |  |  | 800 rpm, 1 h |  |  |
| B_4_C(100) | B_2_O_3_ | Mg | 0.90 | 1.00 | 0.090 | 1.99 | 30:1  (new) | 800 | 6 | 10.0 g NaCl | 120 | 34.5 |
|  | – | 30 |  |  |  |  |  |  |  | 800 rpm, 1 h |  |  |

**Supplementary Table 4** Comparison particular tumor accumulation and therapeutic efficacy as a function of NP type and particle size.

| **NPs type** | **Hydrodynamic Size (nm)** | **Tumor model and animal** | **Time post-injection** | **Tumor stage** | **Tumor accumulation** | **Delivery**  **efficiency (%ID)** | **Therapeutic efficiency** | **Ref** |
| --- | --- | --- | --- | --- | --- | --- | --- | --- |
| **Polymer-based micelle** | **30, 50, 70, 100** | **Murine colon adenocarcinoma (CT26) subcutaneously inoculated to BALB/c nude mice** | **24 h** | **Volume:**  **~100 mm³**  **Mass:**  **~0.10 g** | **30 ≈ 70 > 50 > 100** | **1.00 ≈ 1.00 > 0.85 > 0.80** | **30 ≈ 100 > 50 ≈ 70** | **^47^** |
|  |  | **Human pancreatic adenocarcinoma (BxPC3) subcutaneously inoculated to BALB/c nude mice** | **24 h** | **Volume:**  **~100 mm³**  **Mass:**  **~0.10 g** | **30 > 50 > 70 ≈ 100** | **1.05 > 0.60 > 0.38 ≈ 0.38** | **30 > 50 > 70 ≈ 100** | **^47^** |
| **Polymer-based micelle** | **35, 100, 150** | **Human breast cellosaurus (Bcap-37) subcutaneously inoculated to BALB/c nude mice** | **24 h** | **Volume:**  **~100 mm³**  **Mass:**  **~0.10 g** | **150 > 100 > 35** | **1.46 > 1.20 > 0.15** | **NA** | **^48^** |
| **Polymer-based micelle** | **30, 70, 140** | **Human hepatocellular carcinoma (HepG2) subcutaneously inoculated to BALB/c nude mice** | **48 h** | **Volume:**  **~100 mm³**  **Mass:**  **~0.10 g** | **30 > 70 > 140** | **NA** | **30 > 140** | **^49^** |
| **PEGylated Gold** | **50, 60, 80, 100, 120** | **Human breast melanoma (MDA-MB-435) subcutaneously inoculated to athymic nude mice** | **24 h** | **Volume:**  **1000 ~ 2200 mm³**  **Mass:**  **1.00 ~ 2.20 g** | **60 > 50 > 80 > 100 ≈ 120** | **0.90 > 0.60 > 0.45 > 0.10 > 0.10** | **NA** | **^50^** |
|  |  | **Human breast melanoma (MDA-MB-435) subcutaneously inoculated to athymic nude mice** | **24 h** | **Volume:**  **500 ~ 1000 mm³**  **Mass:**  **0.50 ~ 1.00 g** | **60 ≈ 50 > 80 > 100 > 120** | **0.50 ≈ 0.50 > 0.15 > 0.09 > 0.05** | **NA** | **^50^** |
| **PEGylated Gold** | **20, 40, 60, 80, 100** | **Human breast melanoma (MDA-MB-435) subcutaneously inoculated to athymic nude mice** | **24 h** | **Volume:**  **~1000 mm³**  **Mass:**  **~1.00 g** | **60 > 80 > 100 > 40 > 20** | **1.10 > 0.85 > 0.75 > 0.66 > 0.01** | **NA** | **^51^** |
| **Gold@tiopronin** | **50, 100**  **(Core size)** | **Human breast cancer (MCF-7) subcutaneously inoculated to BALB/c nude mice** | **24 h** | **Volume:**  **~100 mm³**  **Mass:**  **~0.10 g** | **50 > 100** | **0.03 > 0.01** | **NA** | **^52^** |
| **PEGylated Drug–silica nanoconjugates** | **40, 70, 240** | **Human breast cancer (MCF-7) subcutaneously inoculated to athymic nude mice** | **24 h** | **Volume:**  **~170 mm³**  **Mass:**  **~0.17 g** | **70 > 40 > 240** | **0.28 > 0.22 > 0.15** | **70 > 40 > 240** | **^53^** |
|  |  | **Murine mammary carcinoma (4T1) subcutaneously inoculated to BALB/c nude mice** | **24 h** | **Volume:**  **~170 mm³**  **Mass:**  **~0.17 g** | **70 > 40 > 240** | **1.57 > 1.41 > 1.29** | **70 > 40 ≈ 240** | **^53^** |
| **Mesoporous organosilica** | **30, 50, 70, 100** | **Murine mammary carcinoma (4T1) subcutaneously inoculated to BALB/c nude mice** | **24 h** | **Volume:**  **100–200 mm³ Mass:**  **0.10 ~ 0.20 g** | **50 > 30 > 70 > 100** | **NA** | **NA** | **^54^** |
| **DOX@AZIF-8** | **80, 120** | **Murine mammary carcinoma (4T1) subcutaneously inoculated to BALB/c nude mice** | **12 h** | **Volume:**  **50 mm³**  **Mass:**  **~ 0.05 g** | **80 > 120** | **0.40 > 0.30** | **80 > 120** | **^55^** |
| **^10^B_4_C-PG** | **35, 45, 60, 90, 120** | **Murine colon adenocarcinoma (CT26) subcutaneously inoculated to BALB/c white mice** | **48 h** | **Volume:**  **~120 mm³**  **Mass:**  **~0.06 g** | **60 > 45 > 120 > 90** | **4.03 > 3.38 > 2.38 > 1.67** | **60 > 45 > 35 > 120 > 90** | **Present**  **work** |

NA, not available; If only tumor volume was reported, the weight was approximated using an average tumor density^46^ of 1.0 g/cm³. Tumor weight was only precisely reported in our previous and present works.

**References**

1	Kobayashi, T. & Kanda, K. Microanalysis system of ppm-order 10b concentrations in tissue for neutron capture therapy by prompt gamma-ray spectrometry. *Nucl. Instrum. Methods Phys. Res.* **204**, 525-531 (1983).

2	Kawai, K. et al. Cyclic rgd-functionalized closo-dodecaborate albumin conjugates as integrin targeting boron carriers for neutron capture therapy. *Mol. Pharm.* **17**, 3740-3747 (2020).

3	Nomoto, T. et al. Poly(vinyl alcohol) boosting therapeutic potential of *p*-boronophenylalanine in neutron capture therapy by modulating metabolism. *Sci. Adv.* **6**, eaaz1722 (2020).

4	Konarita, K. et al. Poly(vinyl alcohol) potentiating an inert d-amino acid-based drug for boron neutron capture therapy. *J. Control. Release* (2024).

5	Miyabe, J. et al. Boron delivery for boron neutron capture therapy targeting a cancer-upregulated oligopeptide transporter. *J. Pharmacol. Sci.* **139**, 215-222 (2019).

6	Mi, P. et al. Block copolymer-boron cluster conjugate for effective boron neutron capture therapy of solid tumors. *J. Control. Release* **254**, 1-9 (2017).

7	Tokura, D. et al. Active control of pharmacokinetics using light-responsive polymer-drug conjugates for boron neutron capture therapy. *J. Control. Release* **371**, 445-454 (2024).

8	Li, J. et al. A metabolically stable boron-derived tyrosine serves as a theranostic agent for positron emission tomography guided boron neutron capture therapy. *Bioconjug. Chem.* **30**, 2870-2878 (2019).

9	Nishimura, K. et al. Efficient neutron capture therapy of glioblastoma with pteroyl-closo-dodecaborate-conjugated 4-(p-iodophenyl)butyric acid (pbc-ip). *J. Control. Release* **360**, 249-259 (2023).

10	Kim, S., Mushtaq, S., Lee, K. C., Park, J. A. & Kim, J. Y. (64)cu-labeled boron-containing cyclic rgd peptides for bnct and pet imaging. *ACS Med. Chem. Lett.* **15**, 344-348 (2024).

11	Chen, J. et al. A bis-boron amino acid for positron emission tomography and boron neutron capture therapy. *Angew. Chem. Int. Ed. Engl.*, e202413249 (2024).

12	Mo, S. et al. The synthesis and evaluation of novel bpa derivatives for enhanced blood-brain barrier penetration and boron neutron capture therapy. *Chin. Chem. Lett.* (2024).

13	Koganei, H. et al. Development of high boron content liposomes and their promising antitumor effect for neutron capture therapy of cancers. *Bioconjug. Chem.* **24**, 124-132 (2013).

14	Lee, W. et al. Pegylated liposome encapsulating nido-carborane showed significant tumor suppression in boron neutron capture therapy (bnct). *Biochem. Biophys. Res. Commun.* **522**, 669-675 (2020).

15	Kueffer, P. J. et al. Boron neutron capture therapy demonstrated in mice bearing emt6 tumors following selective delivery of boron by rationally designed liposomes. *Proc. Natl. Acad. Sci. U. S. A.* **110**, 6512-6517 (2013).

16	Li, J. et al. Boron encapsulated in a liposome can be used for combinational neutron capture therapy. *Nat Commun* **13**, 2143 (2022).

17	Feng, B. et al. Delivery of sodium borocaptate to glioma cells using immunoliposome conjugated with anti-egfr antibodies by zz-his. *Biomaterials* **30**, 1746-1755 (2009).

18	Maruyama, K. et al. Intracellular targeting of sodium mercaptoundecahydrododecaborate (bsh) to solid tumors by transferrin-peg liposomes, for boron neutron-capture therapy (bnct). *J. Control. Release* **98**, 195-207 (2004).

19	Shi, Y. et al. Tracing boron with fluorescence and positron emission tomography imaging of boronated porphyrin nanocomplex for imaging-guided boron neutron capture therapy. *ACS Appl. Mater. Interfaces* **10**, 43387-43395 (2018).

20	Shi, Y. et al. Covalent organic polymer as a carborane carrier for imaging-facilitated boron neutron capture therapy. *ACS Appl. Mater. Interfaces* **12**, 55564-55573 (2020).

21	Xiong, H. et al. Amphiphilic polycarbonates from carborane-installed cyclic carbonates as potential agents for boron neutron capture therapy. *Bioconjug. Chem.* **27**, 2214-2223 (2016).

22	Ma, W. et al. Molecular engineering of aie-active boron clustoluminogens for enhanced boron neutron capture therapy. *Chem. Sci.* **15**, 4019-4030 (2024).

23	Dai, L. et al. Bpa‐containing polydopamine nanoparticles for boron neutron capture therapy in a u87 glioma orthotopic model. *Adv. Funct. Mater.* (2023).

24	Dai, L. et al. Boronophenylalanine‐containing polydopamine nanoparticles for enhanced combined boron neutron capture therapy and photothermal therapy for melanoma treatment. *Adv. Funct. Mater.* (2024).

25	Chen, J. et al. Remarkable boron delivery of irgd-modified polymeric nanoparticles for boron neutron capture therapy. *Int J Nanomedicine* **14**, 8161-8177 (2019).

26	Chan, W.-J. et al. Engineering a potent boron-10-enriched polymeric nanoparticle for boron neutron capture therapy. *Nanomedicine* **18**, 743-754 (2023).

27	Deng, S. et al. A pd-l1 sirna-loaded boron nanoparticle for targeted cancer radiotherapy and immunotherapy. *Adv. Mater.*, e2419418 (2025).

28	Yinghuai, Z. et al. Substituted carborane-appended water-soluble single-wall carbon nanotubes:  New approach to boron neutron capture therapy drug delivery. *J. Am. Chem. Soc.* **127**, 9875-9880 (2005).

29	Choi, G., Jeon, I. R., Piao, H. & Choy, J. H. Highly condensed boron cage cluster anions in 2d carrier and its enhanced antitumor efficiency for boron neutron capture therapy. *Adv. Funct. Mater.* **28** (2017).

30	Kuthala, N., Vankayala, R., Li, Y. N., Chiang, C. S. & Hwang, K. C. Engineering novel targeted boron-10-enriched theranostic nanomedicine to combat against murine brain tumors via mr imaging-guided boron neutron capture therapy. *Adv. Mater.* **29** (2017).

31	Kuthala, N., Shanmugam, M., Yao, C. L., Chiang, C. S. & Hwang, K. C. One step synthesis of ^10^b-enriched ^10^bpo_4_ nanoparticles for effective boron neutron capture therapeutic treatment of recurrent head-and-neck tumor. *Biomaterials* **290**, 121861 (2022).

32	Li, L. et al. On-demand biodegradable boron nitride nanoparticles for treating triple negative breast cancer with boron neutron capture therapy. *ACS Nano* **13**, 13843-13852 (2019).

33	Gao, Z. et al. Use of boron cluster-containing redox nanoparticles with ros scavenging ability in boron neutron capture therapy to achieve high therapeutic efficiency and low adverse effects. *Biomaterials* **104**, 201-212 (2016).

34	Xiong, H. et al. Doxorubicin-loaded carborane-conjugated polymeric nanoparticles as delivery system for combination cancer therapy. *Biomacromolecules* **16**, 3980-3988 (2015).

35	Li, L. et al. A boron-10 nitride nanosheet for combinational boron neutron capture therapy and chemotherapy of tumor. *Biomaterials* **268**, 120587 (2021).

36	Shanmugam, M., Kuthala, N., Kong, X., Chiang, C.-S. & Hwang, K. C. Combined gadolinium and boron neutron capture therapies for eradication of head-and-neck tumor using gd^10^b_6_ nanoparticles under mri/ct image guidance. *JACS Au* **3**, 2192-2205 (2023).

37	Wang, Z. et al. Multifunctional high boron content mofs nano-co-crystals for precise boron neutron capture therapy for brain glioma in situ. *Nano Today* **45** (2022).

38	Lan, K. W. et al. In vivo investigation of boron-rich nanodrugs for treating triple-negative breast cancers via boron neutron capture therapy. *Biomater Adv* **155**, 213699 (2023).

39	Tang, H. et al. Boron-containing mesoporous silica nanoparticles with effective delivery and targeting of liver cancer cells for boron neutron capture therapy. *ACS Appl. Mater. Interfaces* (2024).

40	Wang, M. et al. Tumor redox‐responsive minimalist b/fe nano‐chains for chemodynamically enhanced ferroptosis and synergistic boron neutron capture therapy. *Adv. Funct. Mater.* (2024).

41	Li, L. et al. Intratumoral transforming boron nanosensitizers for amplified boron neutron capture therapy. *Angew. Chem. Int. Ed. Engl.*, e202413232 (2024).

42	Zhong, T. et al. Human serum albumin-coated (10)b enriched carbon dots as targeted "pilot light" for boron neutron capture therapy. *Adv Sci (Weinh)*, e2406577 (2024).

43	Nishikawa, M. et al. Conjugation of phenylboronic acid moiety through multistep organic transformations on nanodiamond surface for an anticancer nanodrug for boron neutron capture therapy. *Bull. Chem. Soc. Jpn.* **94**, 2302-2312 (2021).

44	Zhang, Y. et al. Tumor eradication by boron neutron capture therapy with ^10^b-enriched hexagonal boron nitride nanoparticles grafted with poly(glycerol). *Adv. Mater.* **35**, e2301479 (2023).

45	Wang, Y. et al. Polyglycerol functionalized (10) b enriched boron carbide nanoparticle as an effective bimodal anticancer nanosensitizer for boron neutron capture and photothermal therapies. *Small*, e2204044 (2022).

46	De Vleeschauwer, S. I. et al. Observe: Guidelines for the refinement of rodent cancer models. *Nat. Protoc.* **19**, 2571-2596 (2024).

47	Cabral, H. et al. Accumulation of sub-100 nm polymeric micelles in poorly permeable tumours depends on size. *Nat Nanotechnol* **6**, 815-823 (2011).

48	Wang, J. et al. The role of micelle size in tumor accumulation, penetration, and treatment. *ACS Nano* **9**, 7195-7206 (2015).

49	Fan, W. et al. Role of micelle size in cell transcytosis-based tumor extravasation, infiltration, and treatment efficacy. *Nano Lett.* (2023).

50	Sykes, E. A. et al. Tailoring nanoparticle designs to target cancer based on tumor pathophysiology. *Proc. Natl. Acad. Sci. U. S. A.* **113**, E1142-1151 (2016).

51	Perrault, S. D., Walkey, C., Jennings, T., Fischer, H. C. & Chan, W. C. W. Mediating tumor targeting efficiency of nanoparticles through design. *Nano Lett.* **9**, 1909-1915 (2009).

52	Huo, S. et al. Superior penetration and retention behavior of 50 nm gold nanoparticles in tumors. *Cancer Res.* **73**, 319-330 (2013).

53	Tang, L. et al. Investigating the optimal size of anticancer nanomedicine. *Proc. Natl. Acad. Sci. U. S. A.* **111**, 15344-15349 (2014).

54	Zhang, J. et al. Size effect of mesoporous organosilica nanoparticles on tumor penetration and accumulation. *Biomater Sci* **7**, 4790-4799 (2019).

55	Duan, D. et al. Size-controlled synthesis of drug-loaded zeolitic imidazolate framework in aqueous solution and size effect on their cancer theranostics in vivo. *ACS Appl. Mater. Interfaces* **10**, 42165-42174 (2018).
